# Stretchable thin-film metal electronics enabled by multilayered nanomembranes

**DOI:** 10.1101/2025.10.30.685700

**Authors:** Dongjun Jung, Camille E. Cunin, Rajib Mondal, Karen K.L. Pang, Yeji Kim, Ethan Frey, Sara Winther, Jacob Beckham, Taylor M. Cannon, Claudia Cea, Aristide Gumyusenge, Polina Anikeeva

## Abstract

Metallic thin films are indispensable in flexible electronics, yet their brittleness under tensile strain has impeded their use in intrinsically stretchable devices. Here, we overcome this barrier with a multilayer platform of alternating nanomembranes of metal and porous elastomer assembled via exponential stacking. The porous elastomer layers anchor adjacent metal layers and facilitate vertical percolation. Under strain, they dissipate stress and laterally misalign cracks within successive metal layers, forming crack-bridging conductive pathways. This architecture achieves synergistic scaling of electrical and mechanical performance with increasing layer number, demonstrating bulk-like conductance at strains exceeding 700% across a wide range of metals. Using nanomembrane stacks of gold and platinum, we fabricated stretchable electrode arrays that enabled high-fidelity recordings and electrical stimulation of murine colonic electrophysiology *in vivo*.

## Main Text

Soft and stretchable electronics enable seamless integration with dynamic and deformable surfaces, empowering the development of electronic skins (*1–2*), smart fabrics (*3, 4*), soft robotics (*5*), and implantable biomedical devices (*6, 7*) (Fig. 1A). These systems demand stretchable conductors that retain metallic conductivity (*8, 9*) and low interfacial impedance (*10*) under substantial strains (*11*). Metals intrinsically meet the electrical demands (*12*), yet existing stretchable metallic conductors, whether based on nanomaterials (*13*) or liquid metals (*14*), are limited by the low processing scalability or narrow material choice. Thin-film metals can be produced at scale, but exhibit low fracture strains (Fig. 1B), motivating the patterning of metal films into serpentine or wavy architectures to achieve structural elasticity. While indispensable in a wide range of applications (*15*), structural elasticity poses limitations to device miniaturization and hinders intimate integration to the deforming tissues. Engineered microcracks have been shown to enable conduction at strains up to 300% without structural elasticity (*16*), but this approach compromises conductance and is restricted to gold (Au) and silver (Ag) (*17*) (supplementary text, section S1). Consequently, there remains a need for a platform that achieves metallic conductivity in thin films while exhibiting intrinsic elasticity to empower applications in stretchable electronics.

**Fig. 1.**
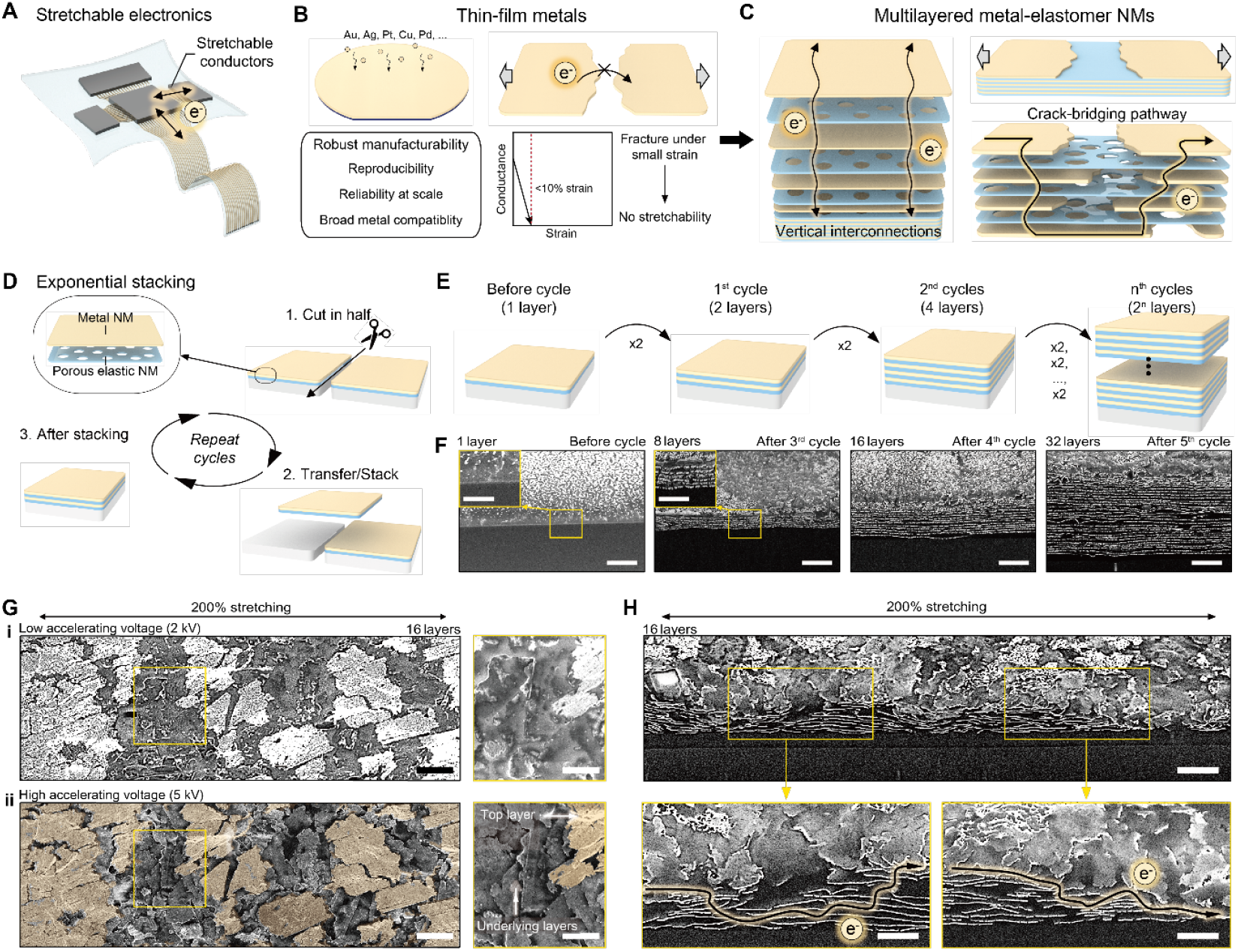
Multilayered metal-elastomer nanomembranes fabricated via exponential stacking. (**A**) An illustration of a stretchable electronic system incorporating stretchable thin-film metal conductors. (**B**) Conventional thin-film metals offer robust manufacturability, but they fracture under small strain, resulting in loss of electrical conductance. (**C**) An illustration of a multilayer metal–elastomer nanomembrane architecture, showing vertical vias between metal layers and crack-bridging pathways under strain. (**D**) An illustration of the exponential stacking process. (**E**)Schematic showing configurations before and after the first, second, and n^th^ stacking cycles. (**F**)Cross-sectional SEM images of Au multilayer structures before stacking (1 layer) and after 3, 4, and 5 cycles (8, 16, and 32 layers, respectively). Scale bars: 2 µm; (insets) 1µm. (**G**) Top-view SEM images of a 16-layer Au architecture acquired at (i) low and (ii) high accelerating voltages. Right images show magnified SEM views. The highlighted region in the bottom image corresponds to the top metal layer observed in the top image. Scale bars: (left) 10 µm, (right) 5 µm. (**H**) Cross-sectional SEM images of a16-layer Au stack under 200% strain. The orange-highlighted lines illustrate crack bridging pathways. Scale bars: (top) 5 µm, (bottom) 2 µm.

### Multilayered metal–elastomer nanomembranes fabricated via exponential stacking

We present a multilayer architecture composed of hybrid metal–elastomer nanomembranes (NMs) that enables the synergistic scaling of electromechanical performance (Fig. 1C). In this design, multiple metal NMs are vertically interleaved with porous elastomer NMs that promote interfacial adhesion between metal layers (*18*). We hypothesized that the porosity of the elastomer NMs would enable direct contact between adjacent metal NMs, creating a dense network of vertical vias (*14*). Furthermore, under strain, elastomer layers are anticipated to dissipate mechanical stress and arrest propagation of cracks within the adjacent metal layers, allowing to build crack-bridging pathways through vertical percolations at strains that would otherwise yield loss of conductivity in fractured metal films (*17*).

To test the hypothesis of synergistic scaling of electrical and mechanical performance in metal-elastomer NM stacks, we developed an exponential stacking strategy that enables efficient and scalable fabrication of multilayered architectures. First, a metal NM deposited via electron beam (or sputtering) onto polytetrafluoroethylene (PTFE) sheet is transferred onto a porous elastomer NM produced by spin-coating, resulting in a single NM bilayer (fig. S1). This bilayer then undergoes repeated stacking cycles, where the existing stack is cut in half, and one half is transferred onto the other (Fig. 1D and fig. S2). With each successive step, the number of layers doubles, as illustrated by cross-sectional scanning electron microscopy (SEM) images of 8, 16, 32, and 64 bilayers of 75±4 nm gold (Au) and 196±11 nm polyisobutylene (PIB) NM produced via 3, 4, 5, and 6 stacking cycles, respectively (Figs. 1E and F, and fig. S3).

We then tested the hypothesis that under strain the elastomer (PIB) would dissipate stress arresting crack propagation between the adjacent metal (Au) layers, resulting in crack misalignment and formation of crack-bridging vertical percolation pathways. Using 16-layer Au-PIB stacks as testbeds, we applied SEM to investigate the structure of the NM multilayers under 200% strain. At a low accelerating voltage (2 kV), SEM images capture only the top metal layer, demonstrating macroscopic cracks that would otherwise interrupt electrical continuity (Fig. 1G and fig. S4). SEM imaging at higher accelerating voltages (5 or 10 kV) penetrates the elastomer layers and reveals underlying metal layers through the cracks in the top layer, corroborating misalignment of cracks. Cross-sectional SEM further showed continuous metal-elastomer interfaces throughout the multilayered architecture under 200% strain (Fig. 1H). The misalignment of metal layer cracks, in combination with robust interlayer adhesion under strain, collectively enable crack-bridging percolation pathways that preserve electrical continuity across macroscopic cracks.

### Synergistic scaling enables unconstrained performance across a broad range of metals

Stretchable metallic conductors have been confined to liquid eutectic gallium-indium alloys, Au, and Ag (*19*). A variety of metals used in high-performance wafer-bound electronics, including Pt (*20*), Pd (*21*), and Cu (*22*), embrittle or oxidize rapidly at the nanoscale, precluding their use in stretchable form. Our multilayered architecture circumvents this challenge by creating crack-bridging percolation pathways that preserve electrical continuity even after macroscale fractures develop (fig. S5).

To demonstrate generality of multilayer NM stacking across different metals, we fabricated structures comprising 1-64 metal NM layers (2-128 layers in total, including elastomer NMs) using Au, Ag, Cu, Pt, and Pd, reaching a total metal thickness of 4.80±0.03 µm (Fig. 2A). Cross-sectional SEM images confirmed direct contact between adjacent metal layers through pores in the elastomer NMs, facilitating vertical charge transport. For all tested metals, sheet conductance increased monotonically with the number of layers (Fig. 2B). Specifically, 64-layer stacks of Au, Ag, Pt, Cu, and Pd exhibited sheet conductance of 23.9±1.7, 27.0±0.7, 3.3±0.2, 8.8±0.6, and 3.3±0.1 S/sq, corresponding to 12.2%, 8.9%, 7.3%, 3.1%, 7.4%, of the sheet conductance expected for bulk metals with 4.8 µm thickness equivalent to the integral thickness of the metal layers within NM stacks.

**Fig. 2.**
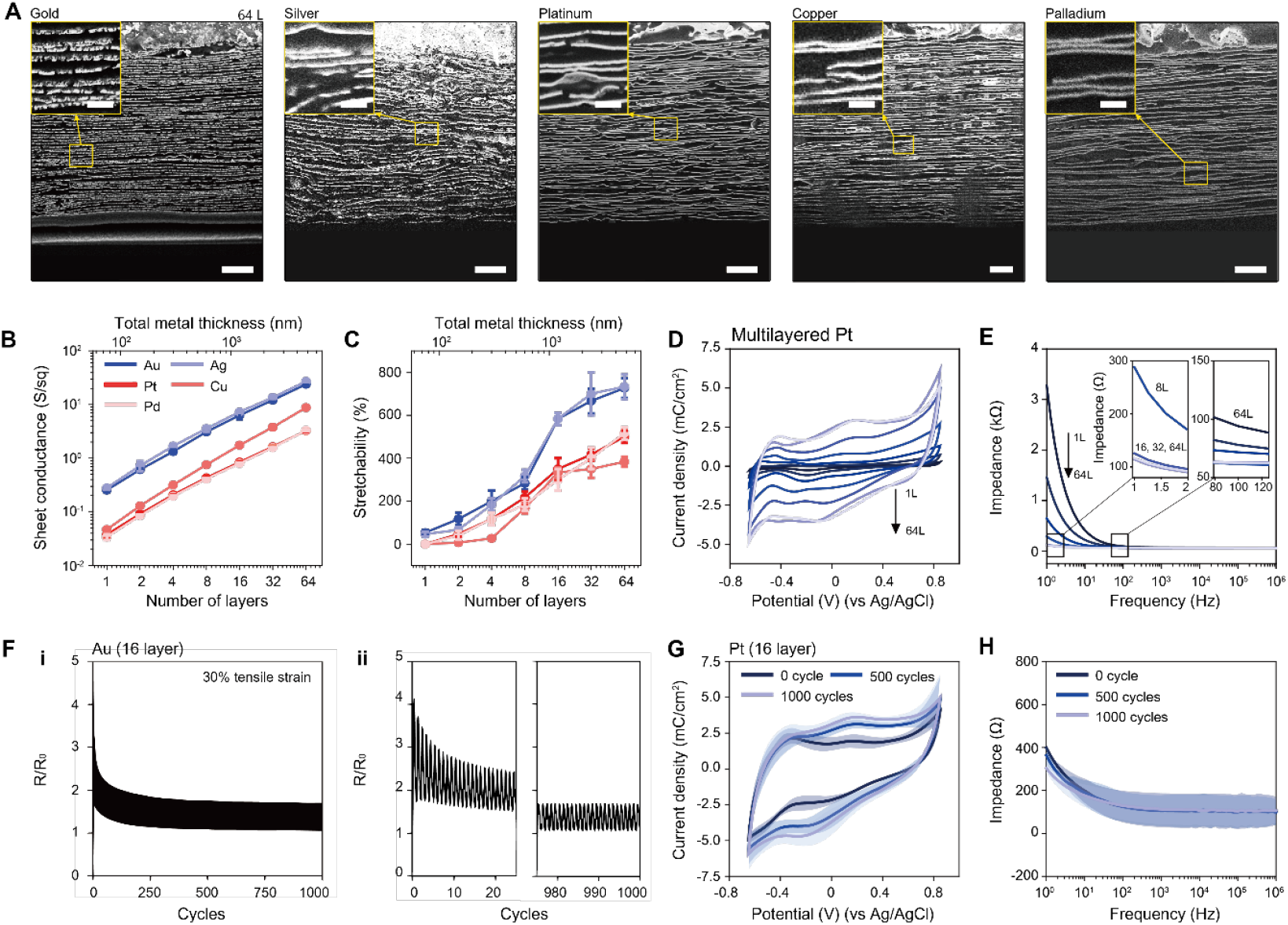
Synergistic scaling of electrical and mechanical performance in multilayered architectures across a broad range of metals. (**A**) Cross-sectional SEM images of 64-layer architectures employing NMs of gold (Au), silver (Ag), platinum (Pt), copper (Cu), and palladium (Pd). Insets: magnified SEM views. Scale bars: 2 µm; (insets) 500 nm. (**B** and **C**) (B) Sheet conductance and (C) stretchability of each metal as a function of the number of layers under optimized conditions (Methods). Markers and error bars represent mean ± s.d. (n = 3). (**D** and **E**) Representative (D) cyclic voltammetry curves and (E) electrochemical impedance spectra (EIS) of Pt multilayered stacks with varying numbers of layers. Insets show impedance near 1 Hz and 100 Hz. Electrode area: 4 mm × 5 mm. (**F**) (i) Resistance variation of a 16-layer Au stack under 1,000 cycles of 30% strain. (ii) The expanded views of the initial and final 25 cycles. (**G** and **H**) (G) Cyclic voltammograms and (H) EIS of a 16-layer Pt stack measured after 0, 500, and 1,000 cycles of 30% strain. Electrode area: 1 mm × 5 mm. Curves represent mean values, and shaded area indicate ± s.d. (n = 3).

Notably, stretchability, which here is defined as the strain value at which the resistance of a sample exceeds 1 kΩ (Methods), also scales monotonically with the number of NM layers (and thus metal thickness) (figs. S6A to E). For 64-metal layer stacks of Au, Ag, Pt, Cu, and Pd, stretchability reaches 727±46%, 733±58%, 500±29%, 383±24%, and 517±29%, respectively (Fig. 2C). Approaches reliant on individual microcracked thin metal films on elastomer substrates do not deliver comparable stretchability: Au rapidly loses stretchability at thicknesses over 100 nm, while Pt exhibits negligible stretchability at any thickness (fig. S6F) (*17*).

Among previously underexplored metals, Pt is particularly attractive for bio-interfacing electronics, owing to its biochemical inertness and favorable charge injection and charge storage properties commonly leveraged in clinic for electrical stimulation and recording of physiology (*23, 24*). Previously, Pt was additively integrated into stretchable platforms as Pt nanoparticles or Pt black coatings, which limited performance and long-term electrochemical reliability (*25*). Our multilayered architecture enables the use of continuous Pt films while its porous structure enlarges electrochemical surface area (ECSA) (fig. S7A). Increasing the number of Pt NMs in multilayer stacks up to 16 yields a significant decrease in impedance and an increase in cathodic charge storage capacity (CSC_c_), while stacking additional Pt NMs yields diminishing returns (Figs. 2D, E, and figs. S7B to D). The same trend appears in double-layer capacitance (C_dl_) (fig. S7E) (*26*): C_dl_ rises from 0.20±0.02 mF cm^−2^ for 1 layer to 6.61±1.14 mF cm^−2^ for 16 layers and then only modestly increases to 7.10±1.13 mF cm^−2^ at 64 layers, suggesting limited fluid access to deeper metal NM layers. The 20 mm^2^ square 16-layer Pt electrode exhibits an impedance of 132±17 Ω at 1 Hz and 60.7±0.7 Ω at 1 kHz and a CSC_c_ of 27.3±8.1 mC cm^-2^.

Finally, we evaluated the reliability of electrical performance of NM multilayer stacks under repeated strain. Stacks comprising 16-layers of Au and PIB NMs were subjected to 1,000 cycles of 30% uniaxial strain while monitoring resistance variation, while 16-layer Pt-PIB stacks were tested under the same conditions using impedance spectroscopy and cyclic voltammetry. In Au NM stacks, resistance initially increased but gradually returned to near its original value (increase of 8.61±2.28%) over successive cycles (Fig. 2F, fig. S8A). This near-complete recovery can likely be attributed to the viscoelasticity of the PIB elastomer NMs, which facilitate stress relaxation and enables formation of vertical percolation pathways (*27*). In Pt stacks, during cycling, impedance decreased by 23.6±6.9% and CSC_c_ increased by 62.8±36.4% (Figs. 2G, H and figs. S8B and C). This can likely be attributed to the enlargement of the pores under strain, which changes geometry of the structure. Post hoc SEM analysis of the cycled NM stacks revealed no signs of fracture or delamination, confirming that the PIB elastomer dissipates mechanical stress, maintains structural integrity, and preserves electrical performance under repeated deformation (figs. S8D and E).

### Optimizing interfacial adhesion and elastomer thickness

Realizing the full potential of the multilayered architecture demands precise control over interfacial adhesion and elastomer layer thickness. Low interfacial energy may yield local delamination and the formation of interlayer gaps that disrupt vertical electrical interconnections (Fig. 3A and fig. S9) and reduce electrical performance. The interlayer gaps may further act as stress concentrators under deformation, promoting crack initiation and propagation leading to large-scale fractures (*28*), which significantly decreases conductance under strain and causes irreversible degradation of electrical performance. In contrast, strong interfacial adhesion minimizes the interlayer gaps and ensures structural integrity, thereby promoting vertical percolation and facilitating crack-bridging pathways, essential for synergistic scaling of conductance and stretchability (Fig. 3B).

**Fig. 3.**
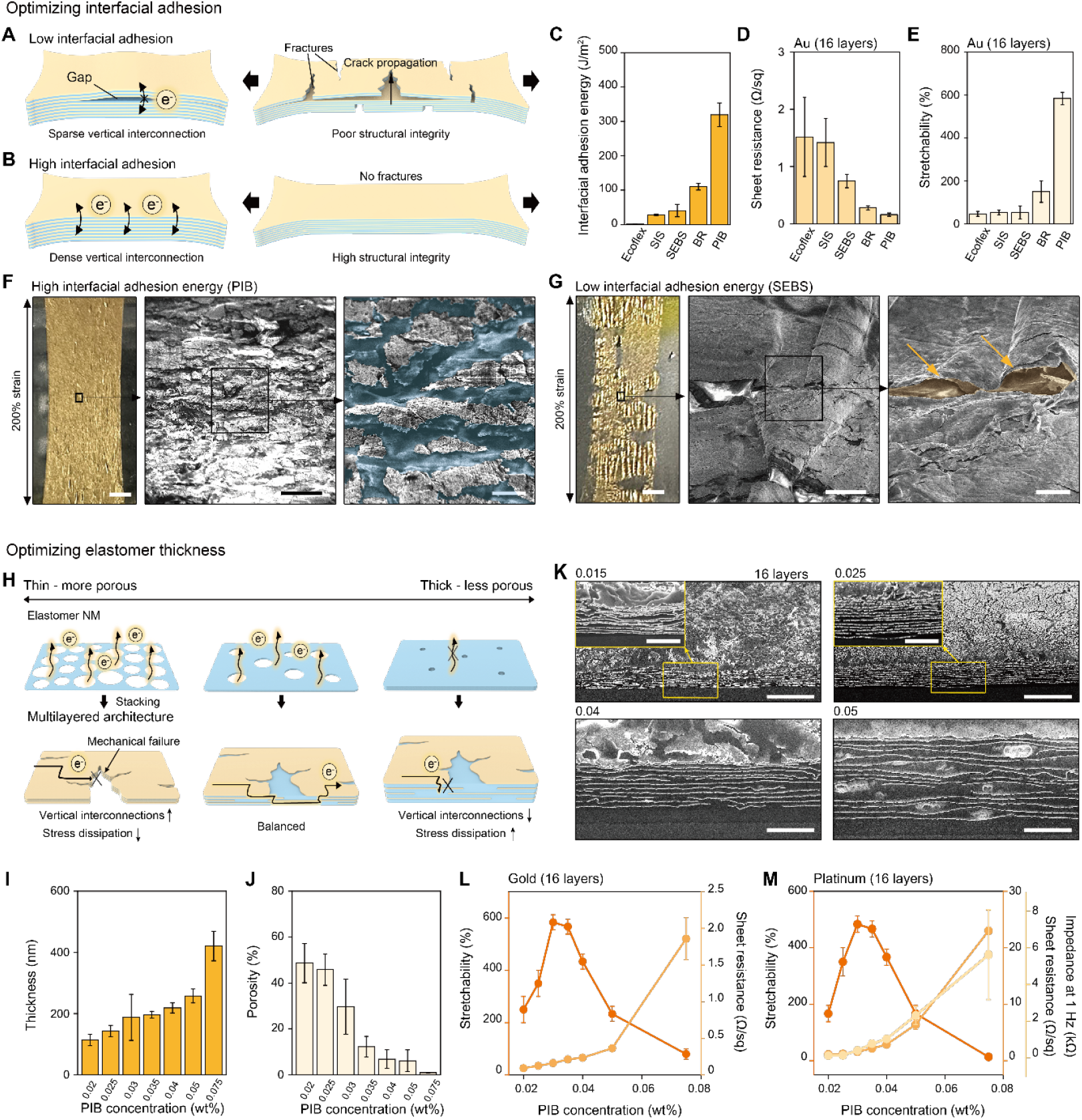
Optimizing interfacial adhesion and elastomer thickness for the multilayered architecture. (**A**) An illustration of the potential effects of low interfacial adhesion on structural integrity under strain. Gaps impede vertical electrical interconnections and lead to fracture under strain. (**B**) An illustration of the potential role of high interfacial adhesion in promoting vertical electrical interconnections and facilitating structural integrity under strain. (**C**) Interfacial adhesion energy between elastomers and gold films. (**D** and **E**) (D) Sheet resistance and (E) stretchability of 16-layer Au stacks interleaved with NMs of different elastomers. (C-E) Bars and error bars represent mean ± s.d. (n = 3). (**F** and **G**) (left) Optical and (middle, right) SEM images of 16-layer Au stack using (F) PIB NM and (G) SEBS NM under 200% strain. The blue-shaded area in (F) corresponds to the PIB layer exposed between microscopic Au cracks. The arrow and orange-shaded region in (G) indicate the onset of delamination and a macroscopic fracture. Scale bars: (left) 2 mm, (middle) 20 µm, and (right) 5 µm. (**H**) An illustration of the effect of PIB thickness on porosity, stress dissipation, and crack-bridging pathways in a multilayer architecture. (**I)** Measured thickness of PIB NMs spin-coated at various PIB concentrations (0.02 – 0.075 wt%) in hexane. (**J**) Quantified porosity percentage of PIB nanomembranes at various PIB concentrations. (I and J) Bars and error bars represent mean ± s.d. (n = 3). (**K**) Cross-sectional SEM images of 16-layer Au stacks using PIB NMs with varying thicknesses, corresponding to PIB concentrations of 0.015, 0.025, 0.04, and 0.05 wt%. Insets: magnified SEM views. Scale bars: 5 µm, (insets) 2 µm. (**L**) Stretchability and sheet resistance of 16-layer Au stacks as a function of PIB concentrations. (**M**) Stretchability, sheet resistance, and impedance at 1 Hz for 16-layer Pt stacks as a function of PIB concentration. (L and M) Markers and error bars represent mean ± s.d. (n = 3).

To evaluate the role of interfacial adhesion, we assessed five elastomers commonly used in bioelectronics and soft robotics: Ecoflex (00-30), poly(styrene-isoprene-styrene) (SIS), poly(styrene-ethylene-butylene-styrene) (SEBS), polybutadiene (BR), and PIB, by measuring their interfacial adhesion energies to Au films via a 180° peeling test (Fig. 3C). Elastomer NMs with a uniform thickness of ~200 nm (fig. S10) were integrated into 16-layer Au stacks to evaluate sheet resistance and stretchability. Sheet resistance decreased with increasing interfacial adhesion (Fig. 3D) from 1.52 Ω/sq for Ecoflex (1.2 J/mm^2^ adhesion energy) to 0.16 Ω/sq for PIB (338.6 J/mm^2^ adhesion energy). An even more pronounced dependence on adhesion was observed for stretchability: Ecoflex, SIS, and SEBS exhibited limited stretchability of 47±12, 53±12, and 53±31%, respectively, BR showed moderate performance of 150±50%, and only PIB achieved stretchability of 583±29 % (Fig. 3E). Optical microscopy and SEM images revealed that the Au-PIB NM multilayers maintained interfacial integrity with no evidence of delamination or catastrophic fractures under strain (Fig. 3F), whereas the other elastomers showed notable signs of interfacial failure and crack formation localized to delaminated regions (Fig. 3G and fig. S11). Although increasing the thickness of the elastomers helped suppress mechanical fracture, interfacial delamination persisted, ultimately limiting stretchability as defined above by maintaining resistance ≤ 1 kΩ (fig. S12). These findings highlight that interfacial adhesion, most effectively provided by PIB, is critical for maintaining structural robustness and conductivity under strain in the multilayered NM architectures.

We next investigated the effect of PIB thickness on the electromechanical performance of the multilayered NM architectures. As the thickness of the elastomer NM decreases, vertical polymer chain entanglement is reduced, limiting packing density and polymer chain mobility (*29*). This hinders uniform condensation of the polymer networks and promotes pore formation (Fig. 3H). The porosity in elastomer NMs facilitates vertical interconnections between metal layers, thereby improving conductance. However, the reduction in polymer chain entanglement compromises mechanical robustness of the elastomer NMs, reducing their ability to dissipate stress and leading to low fracture toughness (fig. S13). Conversely, thicker elastomer NMs offer improved mechanical robustness but exhibit lower porosity, which limits vertical percolation and degrades electrical properties. This tradeoff between the mechanical and electrical functions motivates optimization of the elastomer NM thickness.

To optimize PIB thickness, we employed high-speed spin coating at 6,000 rpm, which promotes rapid solvent evaporation and pore formation (*30*). The thickness of the resulting PIB films was controlled by varying the concentration of PIB in hexane (0.02 to 0.075 wt%). This process yielded PIB NMs ranging from 121±6 nm to 415±61 nm in thickness, with porosity decreasing as thickness increased (Figs. 3I, J, and fig. S14). At concentrations below 0.02 wt%, the films were too thin to maintain continuity (fig. S14). We evaluated electrical and mechanical performance in 16-layer Au and Pt NM stacks (Fig. 3K). For both metals, electrical characteristics: resistance, impedance, and CSC_c_, progressively degraded with increasing PIB thickness (Figs. 3L, M, and fig. S15). In contrast, stretchability showed a non-monotonic trend. Only NMs with thickness values ranging between 144-215 nm (corresponding to 0.025 to 0.04 wt%) allowed for multilayers with stretchability over 300% (as defined by resistance ≤ 1 kΩ). This suggests that sufficient vertical interconnections and stress dissipation can be simultaneously achieved only within a narrow elastomer thickness regime.

Ultraviolet (UV) laser patterning permits inexpensive patterning of NM stacks with consistent feature dimensions down to ~50 µm. Motivated by applications in (bio)electronics where Au electrodes are used as interconnects, while Pt electrodes establish robust electrochemical interfaces, we patterned 16-layer Au NM stacks into 3 cm-long, 50 µm-wide 16-layer Au “traces” and 16-layer Pt NM stacks into 100 µm × 100 µm square “contacts” (fig. S16). Au traces possessed an average resistance of 60.5±1.9 Ω (CV = 3.2%) and Pt contacts exhibited an average impedance of 199±17 kΩ at 1 Hz (CV = 8.3%).

### High-resolution gastrointestinal electrophysiology enabled by the multilayered metal-elastomer nanomembranes

The gastrointestinal (GI) tract is an electrically excitable organ whose coordinated smooth muscle contractions are essential for propulsion of nutrients, fluids, and waste products contained in its lumen (*31*). Disruptions to these electrical rhythms manifest in dysmotility and visceral pain characteristic of GI disorders such as irritable bowel syndrome (IBS) (*32*), colitis (*33*), and gastroparesis (*34*), and are common comorbidities for neurological and psychiatric conditions including Parkinson’s disease (*35*) and autism (*36*). Consequently, high-fidelity recording and targeted modulation of GI electrophysiology holds substantial diagnostic and therapeutic potential (*37*). However, existing GI physiology probes require invasive implantation into the GI wall or lumen, at least in part, due to the inability to reliably operate under the large deformations imposed by GI peristalsis while causing minimal disruption to tissue integrity.

Due to their bulk-like electrochemical performance and resilience to strain, NMs offer a versatile platform for GI bioelectronics. Here, we engineered an eight-channel stretchable NM electrode array tailored for serosal interfacing with a murine colon (Figs. 4A, B). To leverage favorable conductance in the Au NM multilayers with interface impedance and capacitance of the Pt NM multilayers, 16-layer Pt NM stacks were transferred onto the distal tips of 16-layer Au NM stacks (both stacks employed PIB as the elastomer NMs) (figs. S17A). After the transfer, the Au and Pt NM arrays were patterned using a UV laser (figs. S17B). The Au conductive traces were patterned into serpentine lines, a geometry that minimizes piezoresistive fluctuations (fig. S6), thereby assuring consistent signal-to-noise ratio and stimulation during peristalsis. For electrical insulation, the array was conformally encapsulated with SEBS layers (fig. S17A, C). A bioadhesive hydrogel (*38*) ensured intimate integration with the colon serosa without sutures (Figs. 4C, D, and Movie S1). The synergistic benefits of the array architecture and electrode properties coupled with an intimate tissue interface manifested in low *in vivo* impedance of 2.42±0.70 kΩ at 1 kHz across all seven channels (ch. 0 for stimulation and ch. 1-6 for recording) (figs. S17D, E). Among eight channels, the most proximal electrode was designated as the reference (Fig. 4C).

**Fig. 4.**
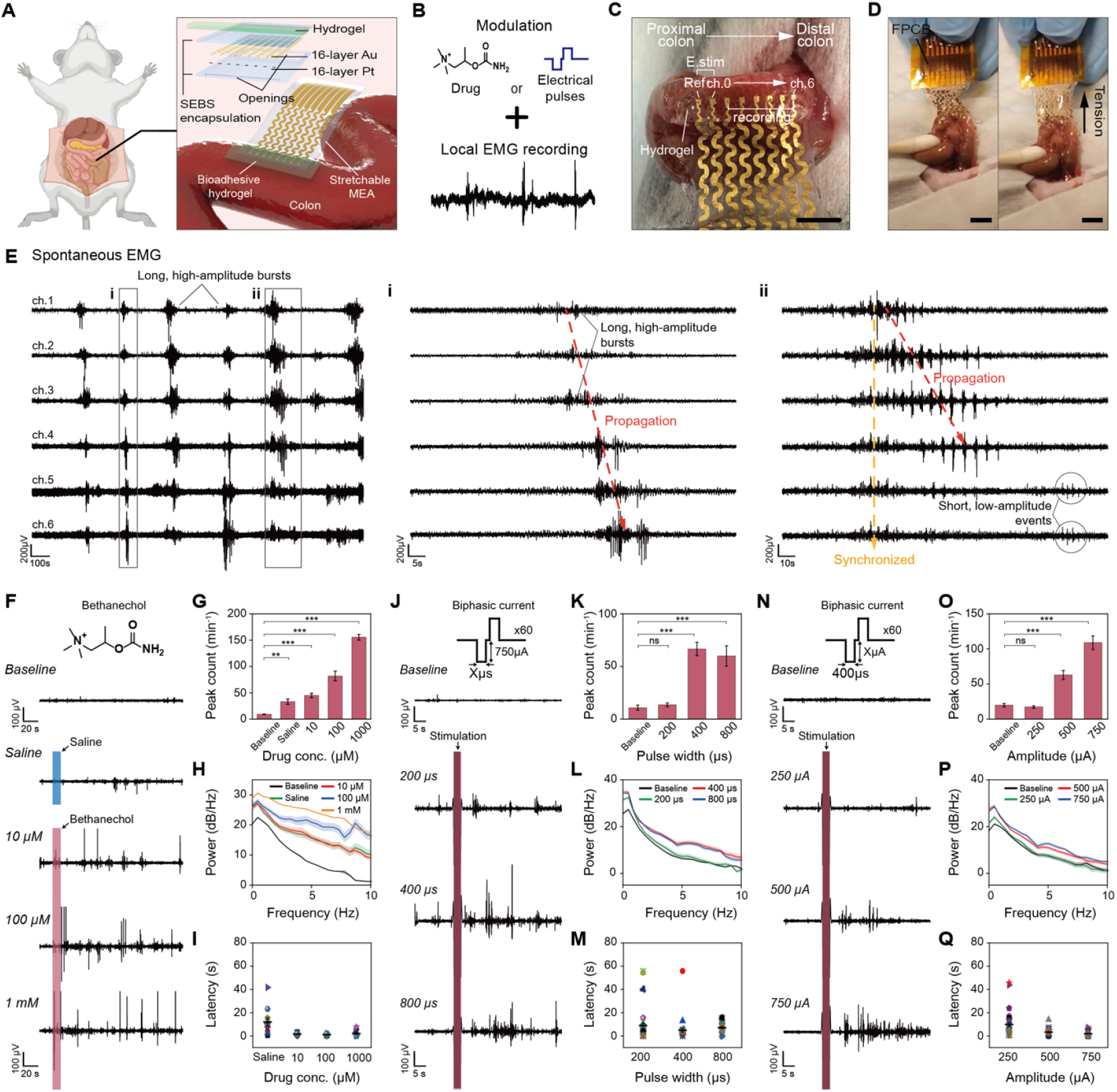
Stretchable nanomembrane electrode arrays enable high-fidelity recording and modulation of colonic electrophysiology in mice. (**A**) Schematic of the device adhered to the colon, along with an exploded view showing layers of the device. (**B**) Electrical and pharmacological stimulation performed concurrently with local EMG recording. (**C**) A photograph of the device placement in vivo. Scale bar: 5 mm. (**D**) Photographs showing robust adhesion of the device to the colonic surface via a hydrogel adhesive. Scale bars: 5 mm. (**E**) Representative spontaneous EMG recordings across channels 1–6. Insets (i) and (ii) show magnified traces. (**F** to **I**) EMG responses to bethanechol injections at varying concentrations: (F) representative EMG signals, (G) quantified peak counts per segment, (H) power spectral densities (0–10 Hz), and (I) latencies under different injection conditions (n = 12). (**J** to **M**) EMG responses to electrical stimulation with varying pulse widths: (J) representative signals, (K) peak counts, (L) power spectral densities (0–10 Hz), and (M) latencies (n = 36). (**N** to **Q**) EMG responses to stimulation with varying pulse amplitudes: (N) representative signals, (O) peak counts, (P) power spectral densities (0–10 Hz), and (Q) latencies (n = 36). Bars and lines denote means; error bars and shaded regions denote ± s.e.m. (F, J, N) Pink shaded regions indicate the injection or stimulation time.

We applied NM electrode arrays to record spontaneous serosal electromyography (EMG) activity in the murine colon in vivo. Despite substantial, continuous tissue deformation, the arrays resolved electrophysiological events consistent with prior *ex vivo* observations of colonic motility patterns. We observed long, high-amplitude bursts recurring every few minutes, resembling neurogenic bursts triggered by enteric neurons, and short, lower-amplitude events occurring every ~10 s, similar to muscle activity controlled by smooth-muscle pacemaker cells (Fig. 4E and fig. S18) (*39, 40*). Notably, high-amplitude bursts coincided with pronounced colonic deformation (Movie S2) and appeared either synchronized across channels or spatiotemporally propagating, indicative of coordinated smooth-muscle activity. Across recordings, these bursting events recurred with a duration of 31.0±21.6 s and an inter-burst interval of 283±201s, consistent with the previous *ex vivo* description of neurogenic bursts (fig. S19) (*39*).

We then assessed the colonic responses to increasing concentrations of cholinergic agonist bethanechol, which increases smooth-muscle activity (*41*). At 5-min intervals, we delivered 20 µL boluses of: saline (vehicle control), followed by 10 µM, 100 µM, and 1000 µM solutions of bethanechol onto the serosal surface (Fig. 4F, figs. S20 to S22). As expected, EMG responses gradually increased with increasing bethanechol concentration. EMG peak counts per minute (counted as crossings of a threshold of 3 standard deviations above baseline signal, 3σ) for baseline, saline, 10 µM, 100 µM, and 1 mM were 9.2±1.4, 33.2±17.1, 44.9±14.2, 81.9±32.1, and 155.6±18.5, respectively (Fig. 4G). Corresponding 0-2Hz band power values were 9.1±2.0, 12.9±3.5 13.3±3.1,14.5±3.0, 19.2±0.9 dB Hz^−1^ (Fig. 4H). Because vehicle application can evoke transient responses, we delivered additional saline control boluses within the same animals (fig. S23). Although saline produced a modest increase in power confined to < 2 Hz frequency band, the response latency (12.0±11.1 s) was significantly longer as compared to bethanechol injections (1.5±0.9 s at 10 µM, 1.1±0.8 s at 100 µM, and 2.1±2.9 s at 1000 µM) (Fig. 4I). Notably, there were no observed changes in the 2–10 Hz band power (fig. S23B to E), supporting that robust low-frequency enhancements observed for ≥ 100 µM bethanechol reflect drug-specific effects rather than injection artifacts.

Leveraging the favorable charge-injection characteristics of Pt NM contacts, we next evaluated local electrical stimulation to modulate colonic motility. Stimulation was delivered between the reference and channel 0 using biphasic, charge-balanced pulses arranged in bursts (burst rate, 20 Hz; burst duration, 3 s; figs. S17F and S24). We first fixed the current amplitude at 750 µA and varied pulse width (200, 400, and 800 µs) (Figs. 4J to M and figs. S25 to S27). Relative to baseline (10.9±12.6), peak counts per minute (3σ threshold) increased to 13.6±8.2, 66.6±40.2, and 59.9±26.6 for 200 µs, 400 µs, and 800 µs, respectively. The 0–2 Hz power rose from 12.7±4.2 dB Hz^−1^ (baseline) to 16.2±5.9, 18.4±5.8, and 18.2±5.8 dB Hz^−1^. Among these pulse widths, 400 µs yielded the largest and most consistent increases with the shortest latency (4.8±9.1 s), and we therefore fixed the pulse width at 400 µs for subsequent amplitude sweeps (250, 500, 750 µA) (Figs. 4N to Q and figs. S28 to S30). As anticipated, responses scaled with amplitude, with 750 µA producing the largest increases in peak rate, spectral and band power, along with the most consistent reduction in latency. Notably, these stimulation conditions reliably evoked high-amplitude bursts accompanied by visible colonic deformation (Movie S3). The consistency of responses to pharmacological (1 mM bethanechol, figs. S31 to S37) and electrical (biphasic, 10 3s-long 20 Hz bursts, 400 µs pulse width, ±750 µA amplitude, figs. S38 to S44) manipulations was then assessed in n=3 additional mice. Time-frequency analyses in both paradigms revealed selective enhancement of 0-50 Hz power consistent with smooth-muscle activity (*42, 43*). High-fidelity recordings from the NM array further resolved spatially distinct EMG responses to both pharmacological and electrical modulation (figs. S45, S46).

These findings illustrate the potential of the NM bioelectronics to empower mechanistic studies of GI neuro-muscular circuits orchestrating motility in physiological and pathological conditions (*44–47*). Notably, our platform is compatible with intact *in vivo* physiology and eliminates the need for insertion into the gut wall or lumen thus preserving neuromuscular junctions and intrinsic and extrinsic innervations, which work synergistically to coordinate GI function (*47–51*).

## Conclusions and outlook

By interleaving thin-film metals with elastomers within multilayer architectures, we convert these rigid materials into stretchable conductors. Exponential stacking of alternating metal and porous elastomer nanomebranes doubles the layer count at each step, breaking the long-standing trade-off between metal thickness (and thus conductance) and stretchability. Crack-bridging pathways formed throughout the architecture preserve electrical continuity even after macroscopic metal fractures, enabling synergistic electromechanical scaling across a broad range of metals, including Pt and Pd, which were previously unattainable in stretchable form. Our fabrication process uses standard thin-film deposition techniques and laser patterning, making it readily compatible with wafer-scale manufacturing. By converting brittle thin-film metals from a limitation into a foundation for stretchable systems, our platform closes the gap between merely flexible and truly stretchable electronics and paves the way toward integrated, imperceptible, tissue-like electronic systems that may redefine next-generation diagnostics and bioelectronic medicines.

## Author contributions

D.J., C.E.C., A.G. and P.A. designed the study and analyzed the results. D.J. and C.E.C. conceived key ideas, performed exponential stacking, fabricated devices, conducted electrical and mechanical characterization, organized data, and prepared figures with input from P.A. and A.G. R.M. and K.K.L.P. optimized surgical procedures for colonic implantation. D.J., C.E.C., R.M., and K.K.L.P. carried out surgeries, applied electrical and pharmacological modulation, recorded colonic electrophysiology, and analyzed data with input from P.A. Y.K. and E.F. supported electrochemical characterization. S.W. supported nanomembranes optical characterization. D.J., C.E.C., R.M., K.K.L.P., Y.K., E.F., J.B., T.M.C., C.C., A.G., and P.A. contributed to the writing of the manuscript.

## Data and materials availability

All data supporting the conclusions of this manuscript are available within the manuscript and supplementary materials.

## Supplementary Materials

Materials and Methods

Supplementary Text

Figs. S1 to S48

Table S1

Movies S1 to S3

