## Supplementary Information for "Stretchable thin-film metal electronics enabled by multilayered nanomembranes"

**The PDF file includes:**

Materials and Methods
Supplementary Text
Figs. S1 to S48
Table S1
Captions for Movies S1 to S3

**Other Supplementary Materials for this manuscript include the following:**

Movies S1 to S3

### Materials and Methods

#### Materials

Ecoflex™ (00-30) was purchased from Smooth-On, and poly(styrene-ethylene-butylene-styrene) (SEBS, H1041) was supplied by Asahi Kasei. Polystyrene-block-polyisoprene-block-polystyrene (SIS, styrene 22 wt%), polybutadiene (BR, cis Mw~200,000), polyisobutylene (PIB, Mw ~500,000), and all other materials and solvents were purchased from Sigma Aldrich unless otherwise specified.

#### Methods

##### Metal deposition

PTFE sheets (Chemfluor) were cut into 6-inch wafer shapes. Gold and platinum films were deposited using an e-beam evaporator (Temescal, FC2000). Silver, palladium, and copper films were also deposited using an e-beam evaporator (AJA International Inc., ATC 20×20×36). All metal layers were deposited to a thickness of 75 nm at a rate of 0.4 Å $s^{-1}$ under a base pressure in the $10^{-6}$ Torr range (fig. S47).

##### Preparation of various elastomer NMs

Solutions of SIS, SEBS, BR, and PIB were prepared by dissolving the polymers in hexane. Ecoflex solution was prepared by mixing parts A and B at a 1:1 ratio and diluting with hexane. Elastomer solutions were spin-coated onto PTFE wafers at 6000 rpm, targeting a ~200 nm thickness. The optimized concentrations determined were 0.025 wt% for SEBS, 0.035 wt% for SIS, 0.1 wt% for BR, 0.03 wt% for PIB, and 0.2 wt% for Ecoflex (fig. S10).

To measure membrane thickness precisely, the NMs were transferred onto $SiO_2$ wafers (University Wafer, Inc., part number 2026) using a polyvinyl butyral (PVb, Sigma Aldrich, P110010) shuttle. The PVb solution (1:10 in ethanol,) was drop-cast onto the elastomer NMs (20 µl per $cm^2$ and dried in oven at 60 °C for 10 min. Once the PVb solidified, the elastomer NM/PVb was detached, and transferred onto the wafer on a 60 °C hot plate. The PVb was then dissolved in ethanol for 30 min to complete the transfer. Thicknesses were verified by profilometry (Bruker, Dektak-XT).

##### Preparation of PIB NMs with varying thickness

PIB solutions were prepared by dissolving PIB in hexane at concentrations ranging from 0.02 to 0.075 wt%. Solutions were spin-coated at 6000 rpm onto PTFE wafers, and the film thicknesses were measured with a profilometer after transferring the elastomer membranes onto $SiO_2$ wafers using a PVb shuttle. Optimal PIB concentrations were established at 0.03 wt% for Au, Ag, and Cu, and 0.025 wt% for Pt and Pd. This differentiation was due to Au, Ag, and Cu typically being used as interconnects requiring high mechanical robustness combined with electrical performance, whereas Pt and Pd were selected for electrodes demanding critical electrochemical performance, which sharply decreased from 0.025 wt% to 0.03 wt% (fig. S15).

##### Exponential stacking

Single-layer hybrid nanomembrane (NM) were first prepared by spin-coating PIB solutions at 6000 rpm onto PTFE wafers. Metal NMs on PTFE wafers were rinsed with ethanol and transferred using the PVb shuttle. Strong adhesion between the PIB and metal NMs ensured retention of the metal layers after PVb removal.

For exponential stacking, the wafer bearing a single-layer hybrid NM was cut in half; one half was transferred via the PVb shuttle onto the other half. After dissolving PVb in ethanol, the process was repeated until the desired number of layers was obtained. For copper-elastomer NMs, the stacked films were treated with HCl (37%) for 1 min to remove oxide layers and enhance interlayer contact.

Interfacial adhesion energy measurements

Titanium (10 nm) and gold (100 nm) films were deposited onto polyester substrates (U.S. plastic, 44093). Elastomer membranes (SIS, SEBS, BR, PIB at 0.1 wt% in hexane and Ecoflex at a 1:1 ratio) were spin-coated onto PTFE substrates at 1000 rpm, transferred onto PET substrates, and secured with epoxy (Gorilla glue, GOR4200101). The resulting elastomer-PET substrates were then attached to the gold film surfaces on a hot plate at 60 °C. A 180-degree peel test was performed via Instron 8848 Microforce Tester equipped with a 50 N load cell. Adhesion force was defined as the peak force prior to fracture. The work of adhesion was calculated as the area under the force–displacement curve up to fracture and normalized by the membrane area (1 cm × 1 cm) to yield the adhesion energy.

Tensile testing

Freestanding 16-layer gold stacks with varying PIB concentrations (0.02, 0.035, 0.05, 0.075 wt% in hexane) were fabricated and cut into rectangular shapes (2 mm × 5 mm). Tensile tests were performed using an Instron 8848 Microforce Tester equipped with a 10 N load cell at a strain rate of 0.1 $s^{-1}$. Data smoothing was conducted using an FFT filter with a three-point window in Origin software (OriginLab, Origin 9.0).

Microscopy

Optical microscopy images were obtained using a Nikon Eclipse LV150N microscope. Scanning Electron Microscopy (SEM) top-view images were captured with a Zeiss Merlin microscope. Cross-sectional SEM images were acquired using an FEI Helios Nanolab 600 dual-beam FIB-SEM, while tilted at 52°. The cross section was prepared by ion milling. SEM images of cyclic-test samples (16-layer Au and Pt stacks before and after 1000 cycles at 30% strain, 1 s per cycle) were captured using the Zeiss Merlin microscope.

Porosity analysis

Porosity was quantified using ImageJ (NIH) from n = 3 brightfield optical images acquired at different, randomly selected membrane locations. A color threshold was applied to distinguish pores from the matrix based on hue, saturation, and brightness. Final porosity was calculated by averaging the pore area fraction of the three regions.

Electrical measurements

Samples for sheet conductance and sheet resistance measurements were prepared with dimensions of 5 mm × 10 mm. Liquid metal (Gallium-Indium eutectic, Sigma Aldrich, 495425) was applied to copper wires (Lovimag, CFT-1IN), which were then attached to the samples with a fixed 5 mm gap to ensure consistent length and width (5 mm each). Sheet resistance was measured using a four-wire probe method with a multimeter (Agilent, 34410A), and sheet conductance was determined as the reciprocal of the measured sheet resistance.
Electrochemical properties were evaluated using a potentiostat (Gamry, Interface 1010E). Pt stacks were cut into 6 mm × 10 mm pieces and mounted onto glass substrates. One end of each Pt stack

was connected to the copper wire using the liquid metal, and the copper wire was covered with polyimide (PI) tape (Dupont, Tape, 18-1S), leaving a rectangular 4 mm × 5 mm window exposed. The assembly was then immersed in phosphate buffered saline (PBS) so that only the exposed region was in contact with the solution. A Pt mesh counter electrode (Sigma Aldrich, 929883) and an Ag/AgCl reference electrode (Sigma Aldrich, BASMF2052) were also immersed in the PBS solution to establish a three-electrode electrochemical cell setup, with the Pt stack (via the copper wire connection) serving as the working electrode. The working and counter electrode were positioned 1 cm apart.

Cyclic voltammetry (CV) was conducted at a scan rate of 100 mV/s. The charge storage capacity ($CSC_c$) was obtained by integrating the negative current over time. Impedance spectra were recorded using potentiostatic electrochemical impedance spectroscopy (EIS) method. The electrochemically active surface area (ECSA) was estimated from double-layer capacitance ($C_{dl}$) extracted via Randles-circuit fitting using Gamry software.

Coupled electrical and mechanical measurements

(i) stretchability tests.

Hybrid metal–elastomer NM stacks (5 mm × 10 mm) on PTFE were transferred onto VHB tape (3M 4910). Owing to the stacks' integrity, transfer required no additional shuttles. Samples were mounted on a manual tensile stage (Toauto; B07BMJC3WF_CA NARF) with an initial length of 5 mm. Liquid metal (Ga–In eutectic) was applied to both ends to match the stage gap precisely. Resistance during extension at 0.1 mm $s^{-1}$ was measured by a four-wire method (Agilent 34410A). Stretchability was defined as the strain at which resistance exceeded 1 kΩ.

(ii) cyclic stretching (Au stacks).

16-layer gold stacks (5 mm × 10 mm) were transferred onto VHB tape and mounted on a motorized tensile stage (Zaber X-LSM100A-E03; initial length 5 mm). Electrical contacts were made with copper leads using Ga–In liquid metal. Current was continuously recorded with a potentiostat (Gamry) during 1,000 cycles at 30% strain, 1 s per cycle.

(iii) cyclic stretching with electrochemical recording (Pt stacks)

16-layer Pt stacks (1 mm × 10 mm) were transferred onto VHB tape and mounted on the same tensile stage with an initial length of 5 mm. Electrical connection was made using copper tape with the liquid metal. Polyimide (PI) tape was applied at both ends so that the 1 mm × 5 mm window exactly spanned the initial 5-mm gap and remained the portion subjected to extension during stretching. The electrochemical setup matched that described above (three-electrode cell in PBS; Pt stack working, Pt mesh counter, Ag/AgCl reference). The cyclic stretching to 30% strain was performed for 1,000 cycles at 1 s per cycle; electrochemical impedance spectroscopy and cyclic voltammetry were measured at cycles 0, 500, and 1,000.

Nanomembrane electrode array fabrication

Au and Pt were used as interconnects (traces) and tissue-interfacing electrodes (pads), respectively. Sixteen-layer Au and Pt stacks (with PIB interlayers) on the PTFE substrate were cut into rectangles with areas of 2 cm × 5 cm and 1 mm × 18 mm, respectively. The Pt stack was then transferred onto the Au stack using the PVb shuttle, precisely positioned in the electrode regions (figs. S17A, B). The combined Au-Pt stack was patterned using UV-laser ablation (LPKF, U4), employing the parameters: frequency, 200 kHz; power 0.3 W; mark speed, 500 mm/s; repetition 2. The platinum electrodes were patterned into 500 µm × 500 µm squares, and the Au interconnects were defined into serpentine geometries (500 µm width and 500 µm radius) to account for the

observed modest piezoresistive behavior (Fig. 2F), which does not significantly affect bulk conduction, but may influence signal-to-noise ratio in sensitive electrophysiological recordings.
Encapsulation layers were prepared from SEBS membranes spin-coated (10 wt% SEBS in hexane) onto PTFE substrates. The SEBS membranes were then cut into 2 cm × 5 cm rectangles and patterned via UV-laser ablation (frequency, 25 kHz; power 3 W; mark speed, 1000 mm/s; repetition 5). The top encapsulation layer was designed to fully cover the NM electrode array, while the bottom encapsulation layer was patterned to create precise openings for tissue-interfacing electrodes and interconnects to a flexible printed circuit board (FPCB).
Layer integration proceeded sequentially. First, the NM electrode array was transferred onto the top SEBS encapsulation layer using the PVb shuttle, followed by alignment and transfer of the bottom SEBS layer such that its openings precisely matched the Pt electrode sites (fig. S17C). Complete NM electrode arrays were subsequently bonded to flexible PCBs (FPCBs) for interfacing with electrophysiological recording and stimulation system (fig. S17D)
FPCBs were fabricated by patterning copper on polyimide film (Pulsar, G10/FR4) using the UV laser system (frequency, 25 kHz; power 3 W; mark speed, 250 mm/s; repetition 1). On the array's interconnection end, Au traces were patterned to align with the corresponding FPCB contact pads. The patterned end of the NM array was transferred and aligned onto the FPCB, secured with the PI tape. The robust adhesion between the NM electrode array and the FPCB was ensured by applying heat (130 °C) using a mini-iron (K Kernow, LC-708).

Synthesis and fabrication of bioadhesive hydrogel layers
Bioadhesive hydrogel was prepared according to previously published protocols (*42*) with minor modifications. Poly(vinyl alcohol) (PVA, Mw ~ 89,000–98,000) was dissolved in deionized water (DI) at a concentration of 7 wt%. From this PVA solution, 1 mL was taken and mixed with 350 μL acrylic acid, 2 mg α-ketoglutaric acid, and 0.5 mg poly(ethylene glycol) dimethacrylate (PEGDMA, Mn ~750). Nitrogen gas ($N_2$) was purged through the solution for 10 minutes, followed by the addition of 30 mg acrylic acid N-hydroxysuccinimide ester (AA-NHS). The resulting solution was cast into a glass mold with a defined gap of 100 μm and cured under UV LED exposure (395 nm, 50 W) for 30 minutes. The upper glass mold was removed, and the hydrogel was vacuum dried for at least 12 hours and then used in a dry state. The hydrogel films were then attached to the top SEBS encapsulation layers of the NM electrode arrays on a 60 °C hot plate (Fig. 4A).

*In vivo* recordings and modulation in anesthetized mice
All animal procedures were approved by the Massachusetts Institute of Technology Committee on Animal Care (MIT CAC, Protocol #2306000538) and conducted in accordance with institutional and federal guidelines for the care and use of laboratory animals.
Colonic electromyography (EMG) activity recordings were performed in adult C57BL6/J mice (n = 5, 8–10-week-old males) under isoflurane anesthesia (induction at 3%, maintenance at 1–1.5% in 100% oxygen). Once anesthetized, mice were placed supine on a heating pad to maintain core body temperature at 37°C. A midline laparotomy was performed to expose the colon for direct device interfacing. Depth of anesthesia was confirmed throughout the procedure by monitoring respiration and toe-pinch reflex.
The NM electrode arrays were affixed to the serosal surface of the proximal colon using the bioadhesive hydrogel. After positioning the device with the hydrogel on the tissue, 200 μL of PBS was applied onto the hydrogel to facilitate the adhesion. The proximal-most electrode on the array

was designated as reference electrode, and a platinum iridium oxide alloy wire electrode (Thermo Scientific Chemicals, 43019-BS) was placed subcutaneously to serve as electrical ground. Recordings were acquired at 30 kHz using the Ripple Neuro Pico, Grapevine Neural Interface system and Trellis software (Ripple LLC, Salt Lake City, UT, USA).

(i) Baseline recording with saline addition

Before each recording session, 100 µL of PBS was evenly dispensed over the exposed serosal surface to keep the tissue moist. Colonic activity was recorded for 10 min as the saline-on baseline. After 10 min, an additional 100 µL of PBS was uniformly applied across the exposed tissue surface, and recording continued for 20 min (total ≈ 30 min) to assess the incremental saline effect.

(ii) Electrical stimulation parameter optimization

After re-wetting with 100 µL of PBS, electrical stimulation was then delivered via channel 0 and reference of the NM microelectrode array. Stimulation consisted of 3-s trains of biphasic current pulses (20 Hz; 60 pulses per train; 750 µA per phase), repeated every 60 s (*46, 47*). Three pulse-width conditions were tested—200 µs, 400 µs, 800 µs per phase—with corresponding interphase intervals of 40, 80, 160 µs, respectively. Each condition comprised three 3-s trains delivered at 60-s intervals, with 5-min washout between conditions (e.g., 5:00, 6:00, 7:00 min for 200 µs; 13:00, 14:00, 15:00 min for 400 µs; 21:00, 22:00, 23:00 min for 800 µs). Recording was continuous before, during, and after stimulation (total ≈ 30 min). A second series was then performed (after re-wetting the tissue with 100 µL of PBS) to vary current amplitude at a fixed pulse width (400 µs; interphase interval: 160 µs): 250 µA, 500 µA, 750 µA.

(iii) Pharmacological dose-response characterization

Following electrical-stimulation sessions (and after re-wetting with 100 µL of PBS), pharmacological stimulation was performed on the same animal and device. After a 5-min baseline, successive 20 µL applications were pipetted onto the serosa 3–5 mm from the reference electrode over 5 s at 5:00, 10:00, 15:00, 20:00 min: PBS (vehicle) at 5:00, then bethanechol chloride (Sigma-Aldrich) at 10 µM, 100 µM, and 1 mM (in PBS). A final 10 min recording (20:00–30:00 min) captured responses following the 1 mM dose.

(iv) Electrical stimulation

With parameters fixed (and after re-wetting with 100 µL of PBS), we acquired a 600-s baseline of spontaneous colonic EMG, then delivered serosal stimulation via channel 0 (reference electrode completing the circuit): biphasic, 750 µA per phase, phase width 400 µs, interphase 80 µs, 20 Hz burst. Each burst lasted 3 s (60 pulses/burst) and was applied 10 times with 60-s interburst intervals. Recording was continuous for 1800 s, spanning baseline, interburst, and post-stimulation periods to capture immediate and sustained effects.

(v) Pharmacological stimulation

Using the same device/animal (and after re-wetting with 100 µL of PBS), we recorded a 600-s baseline, then locally applied 20 µL of bethanechol chloride (1 mM in PBS) to the proximal colon, 3–5 mm from the reference electrode and over 5 s. EMG activity was recorded continuously before, during, and after application.

Electrophysiological data analysis

Raw electrophysiological signals were analyzed using custom codes written in MATLAB (Mathworks, MATLAB R2024a). Raw electrophysiological data were loaded via Ripple Grapevine's API and custom functions in MATLAB, with a sampling frequency of 30 kHz for all channels. A series of notch filters were applied to each channel to remove 60 Hz noise (U.S. power line interference) and its harmonics up to 420 Hz. Each notch filter was implemented as a second-order Butterworth bandstop filter with a narrow stopband of ±1 Hz around the target frequency and a quality factor of 5. For each burst window, data across all channels were zeroed within a 5-s window extending 1 s before and 1 s after the burst to eliminate artifacts. For drug stimulation, data were zeroed out within a 5-s window corresponding to the period of drug administration.

(i) Filtering of raw signals into the EMG band

Raw electrophysiological traces (with previously identified bursts or drug window removed) were band-pass filtered to isolate EMG-band activity. A 2nd-order Butterworth filter was applied with cutoff frequencies of 1 Hz and 50 Hz, and zero-phase filtering 'filtfilt' was used to prevent phase distortions.

(ii) EMG power

To quantify the temporal evolution of EMG activity, the rectified EMG-band signals were used to compute power over time. Rectification ensures that all voltage fluctuations contribute positively to the signal energy. Power was calculated as the moving average of the squared rectified signal (RMS approach), using a 250 ms sliding window to capture rapid EMG fluctuations while smoothing high-frequency noise. A step size of 50 ms was used for the moving average to balance temporal resolution with computational efficiency, effectively downsampling the power signal to 20 Hz. The 250 ms window was chosen to capture physiologically relevant EMG fluctuations without excessive smoothing, and the 50 ms step provided a sufficient temporal sampling rate to resolve rapid changes in muscle activity while keeping the dataset manageable for visualization.

(iii) Time-resolved peak rates

Time-resolved EMG peak rates were computed from the smoothed power envelope of rectified EMG-band signals for each channel. The power envelope was calculated as a moving mean of the squared rectified signal using a 250 ms window, which provides sufficient smoothing to reduce high-frequency noise while preserving physiologically relevant fluctuations. Peaks were detected within consecutive, non-overlapping 1 s windows, chosen to balance temporal resolution and statistical reliability in counting discrete bursts. Baseline activity during the first 180 s was used to calculate the mean ($\mu$) and standard deviation ($\sigma$) of the power envelope for each channel. Detection thresholds were defined as $\mu + k \cdot \sigma$, with k = 3, 5, and 10. These multiple thresholds allow identification of small, moderate, and high-amplitude events relative to baseline, capturing a spectrum of EMG activity while accounting for channel-specific variability. For each window, peaks exceeding each threshold were identified using MATLAB's 'findpeaks' function, and the number of peaks per bin was recorded as the time-resolved event rate (events $s^{-1}$). Burst windows identified from prior analyses were shaded in black to indicate ongoing activity.

(iv) Time–frequency representations (spectrograms)

Spectrograms were computed using short-time Fourier transforms on notch-filtered signals from each channel. Two types of analyses were performed: low-frequency EMG-band activity (0–

50 Hz) and broadband activity (0–15 kHz). For the low-frequency EMG-band activity (0–50 Hz), signals were downsampled from 30 kHz to 200 Hz to satisfy the Nyquist criterion while reducing computational load, since the highest frequency of interest was 50 Hz. Spectrograms were generated using a 500 ms Hamming window with 450 ms overlap (90%) and a 1024-point FFT. This combination provided adequate frequency resolution (~2 Hz) for EMG-band power analysis while maintaining reasonable temporal precision. High overlap ensured smooth transitions between windows and minimized spectral leakage. For the broadband activity (0–15 kHz), full sampling at 30 kHz was retained to preserve high-frequency bursts. Spectrograms were computed using a 1-s Hamming window with 50% overlap and a 1024-point FFT. The longer window improved frequency resolution for high-frequency components, while the moderate overlap reduced computational load for the large dataset. This configuration allowed clear visualization of bursts across the full frequency spectrum without losing temporal context.
For both analyses, power (P) was expressed in decibels ($10 \cdot \log_{10}(P)$) and visualized using a perceptually uniform 'parula' colormap with a consistent dynamic range (–10 to 20 dB) across channels. Burst windows, identified from prior analysis, were shaded in red.

(v) Power spectral densities (PSDs)
PSDs were computed in MATLAB using Welch's method for each segment and electrode (2-s windows, 50% overlap, NFFT = 2^nextpow2(N)). For condition comparisons, we defined the stimulation windows as noted above. PSDs were computed per electrode within each window and averaged across electrodes to obtain a mean baseline PSD and a mean PSD for each stimulation window. Band power in the 0–2, 2–5, and 5–10 Hz ranges was obtained by integrating the linear PSD within each frequency band. The power was displayed over the 0–10 Hz range. Following prior physiological and signal-processing studies, the spectrum was divided into three frequency bands—0–2 Hz (myogenic/low-frequency), 2–5 Hz (neurogenic central band), and 5–10 Hz (high-frequency band including the ~7 Hz component of the proximal colon)—to compute and compare both band power (area) and mean power (*49*).

(vi) Latency to first 3 σ burst
For each stimulation onset, the latency to the first envelope peak exceeding the 3 σ threshold was measured. Latency was defined as the time difference between stimulation onset ($t_s$) and the first detected high-amplitude peak ($t\_{first}$), i.e., latency = $t\_{first} - t_s$.

(vii) Segmentation of analysis windows (for peak counts, spectral and band power, and latency).
- Pharmacological ladder: the response window was a 290-s period starting 5 s after injection onset; the baseline window was a 290-s period ending 5 s before injection onset
- Electrical-stimulation ladder (pulse-width and amplitude series): the response window was a 50 s period beginning 5 s after stimulation onset (i.e., 2 s after the 3-s train ended). The baseline window was a 150-s period ending 2 s before stimulation onset. The 150-s baseline was split into three 50-s segments to match the number and duration of response segments. These windows were chosen to equalize segment counts and durations across conditions for peak-count analysis
-Electrical stimulation with fixed parameters (interbursts): ten 50-s interburst segments per condition were extracted from 5 s after the train onset to 5 s before the next stimulation. The baseline window was a 180-s period ending 2 s before stimulation onset.
- Drug stimulation with fixed parameters (1 mM bethanechol): the baseline window was a 290-s

period ending 5 s before injection; the post-drug window was a 290-s period starting 5 s after injection.
These windows were chosen to minimize artifacts from stimulation/injection.

(viii) Statistical analysis
To assess stimulation- or drug-induced changes, linear mixed-effects models (LMMs) were fitted in MATLAB (fitlme, Statistics and Machine Learning Toolbox) on two complementary measures: (a) segment-level peak counts and (b) segment-averaged PSDs. Condition (e.g., baseline, saline, or stimulation intensity) was modeled as a fixed effect, with random intercepts for Animal and Electrode (nested within Animal) to account for repeated measures. Fixed-effect coefficients were tested via F-tests (Satterthwaite approximation). All $p$ values are reported in Table S1.

### Supplementary Text

#### Text S1

#### Limitation of current deposition-based method (thin-film-on-elastomer)

To prevent fracture, microcracks are induced in thin-film metals by depositing them directly onto elastomer membranes. These microcracks dissipate applied strain and help preserve the integrity of metal networks under deformation. However, this strategy faces two critical limitations.

First, strain-accommodating microcracks are only formed in ultrathin metal films (< 100 nm). When thicker films are deposited, these cracks become filled, diminishing their ability to dissipate stress. For example, SEM images show well-developed microcracks in 75 nm-thick gold films, which enable stress dissipation under strain (figs. S48A, B). In contrast, 200 nm-thick gold films exhibit filled cracks with minimal dissipation points, resulting in fracture under strain (fig. S48C and D). Thus, the limited thickness constrains the conductance.

Second, this approach is highly affected by mechanical properties of metals. In ultrathin metal films (<100 nm) where dislocation density is low, the degree of dislocation entanglement becomes critical for resisting stress concentration. Metals with low stacking-fault energy, such as Au and Ag, facilitate partial dislocation and energy dissipation, thereby preventing fracture (figs. S48A, B). In contrast, metals with higher stacking-fault energy, such as Pt and Pd, exhibit poor dislocation entanglement and are prone to concentrate stress and fracture even under very small strains. This fracture severs the conductive pathways, leading to loss of electrical functionality (fig. S48E to H). Collectively, these limitations underscore not only the fundamental trade-off between electrical performance and stretchability but also the restricted metal compatibility of current thin-film metal strategy (fig. S6F).

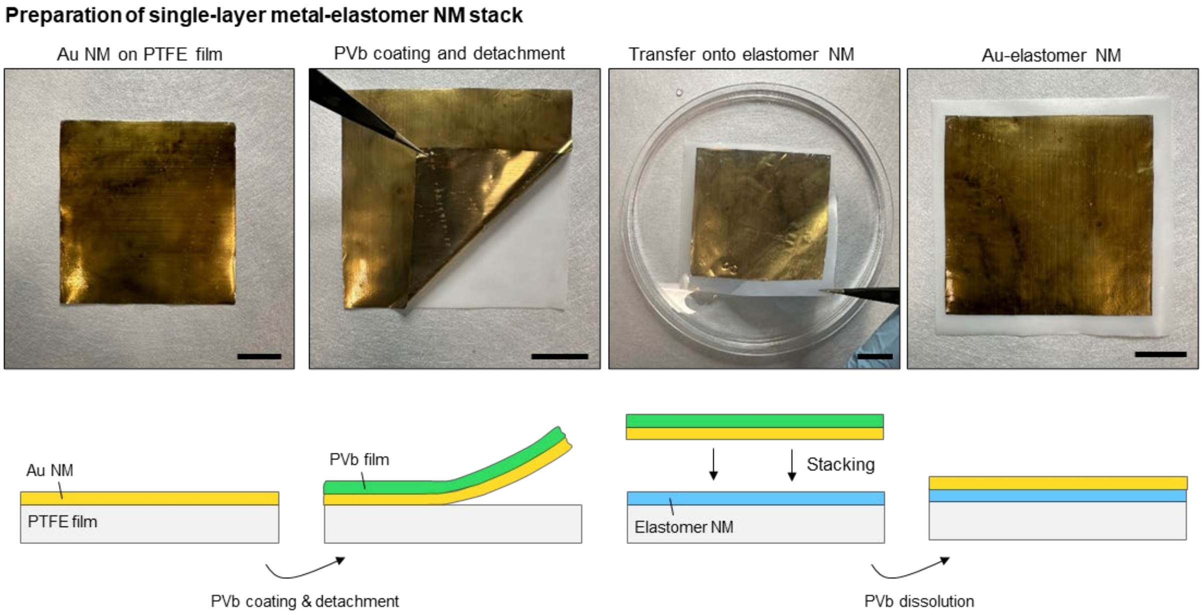


**Fig. S1. Photographs and illustrations showing the preparation of a single-layer Au–elastomer NM stack.** Scale bars: 2 cm.

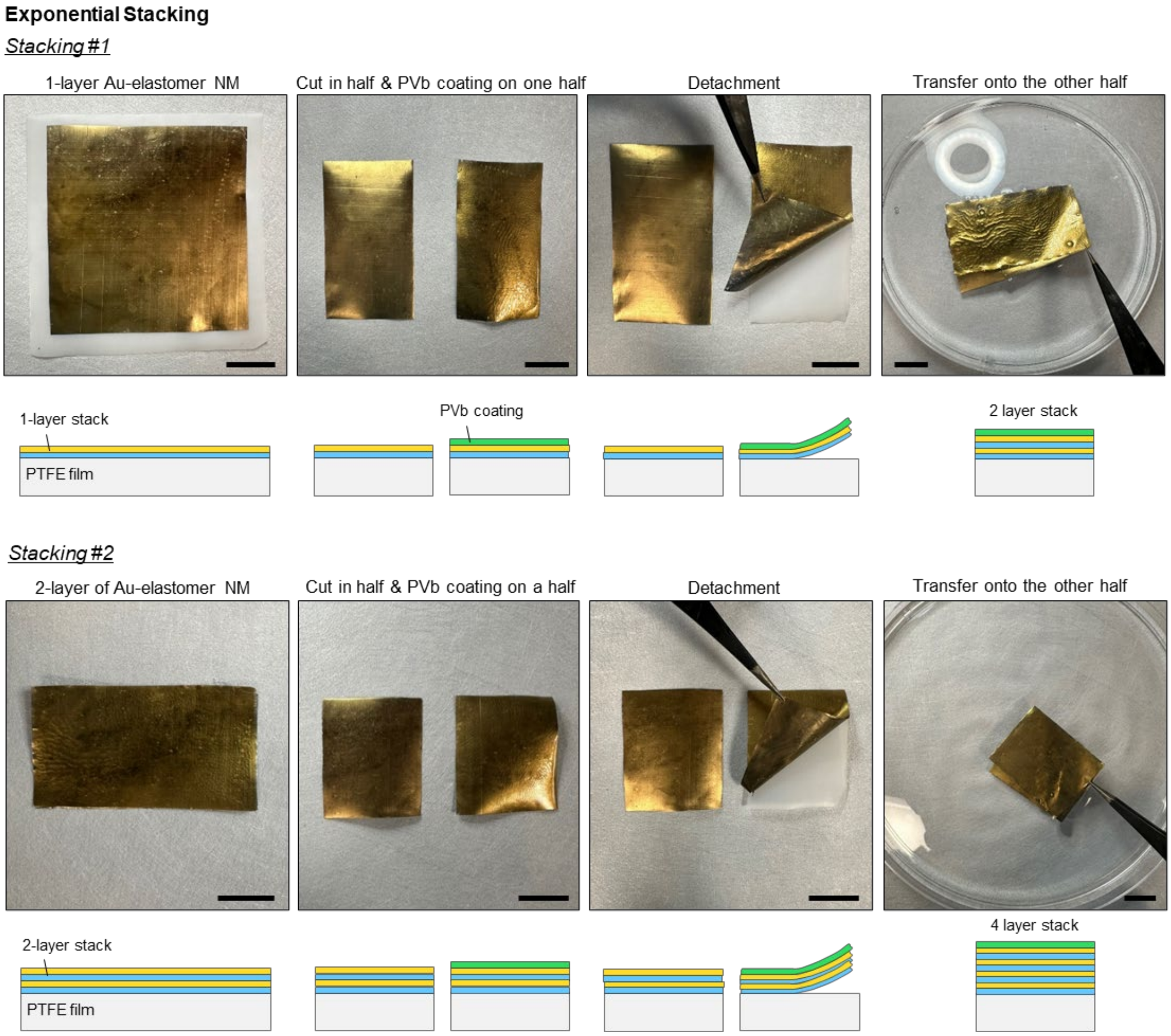


**Fig. S2. Photographs and illustrations showing the process of exponential stacking.** Scale bars: 2 cm.

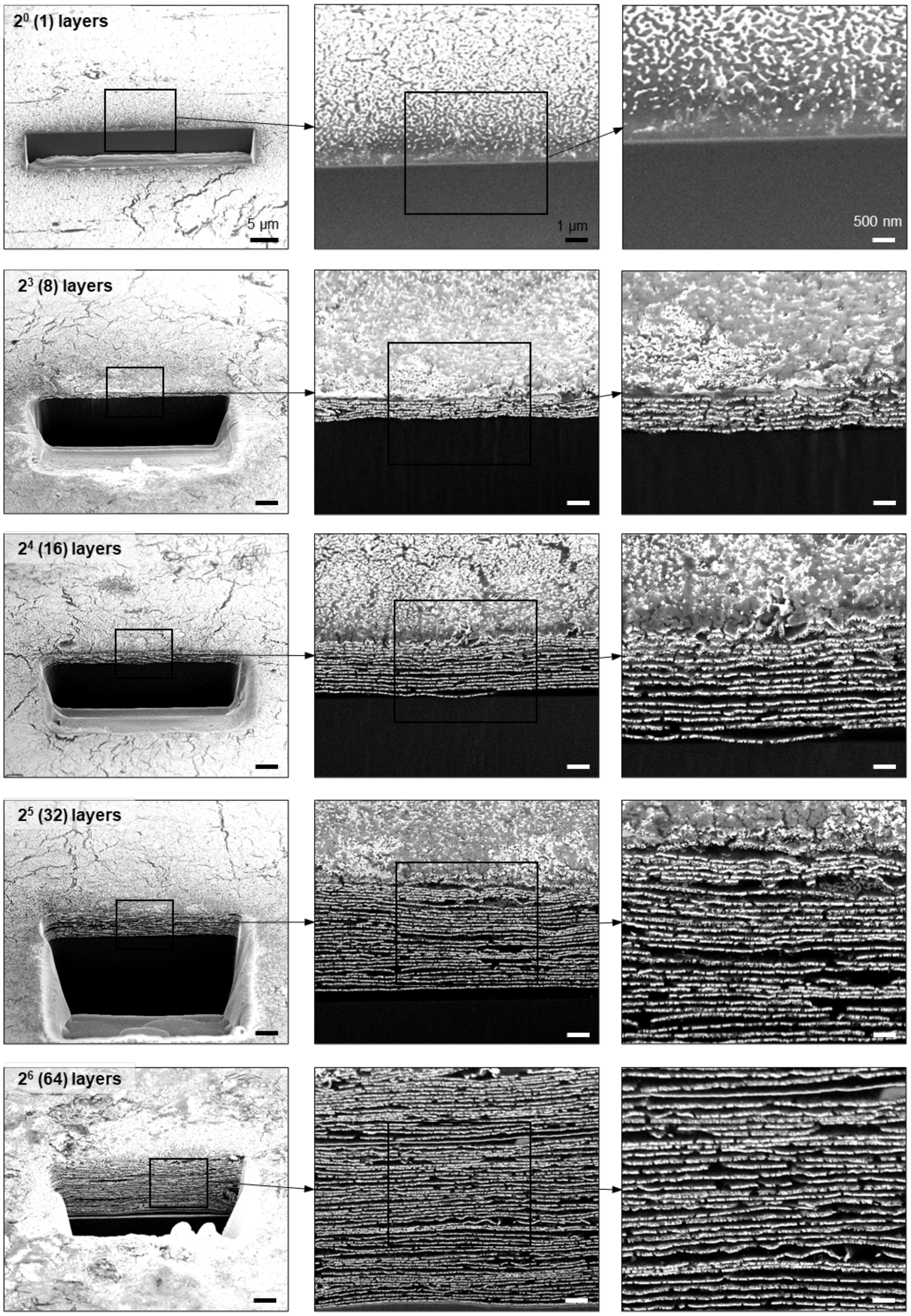


**Fig. S3. Cross-sectional SEM images of multilayered Au architectures** prior to stacking (1 layer) and after 3, 4, and 5 stacking cycles (8, 16, and 32 layers, respectively). Scale bars: (left) 5 µm, (middle) 1 µm, (right) 500 nm.

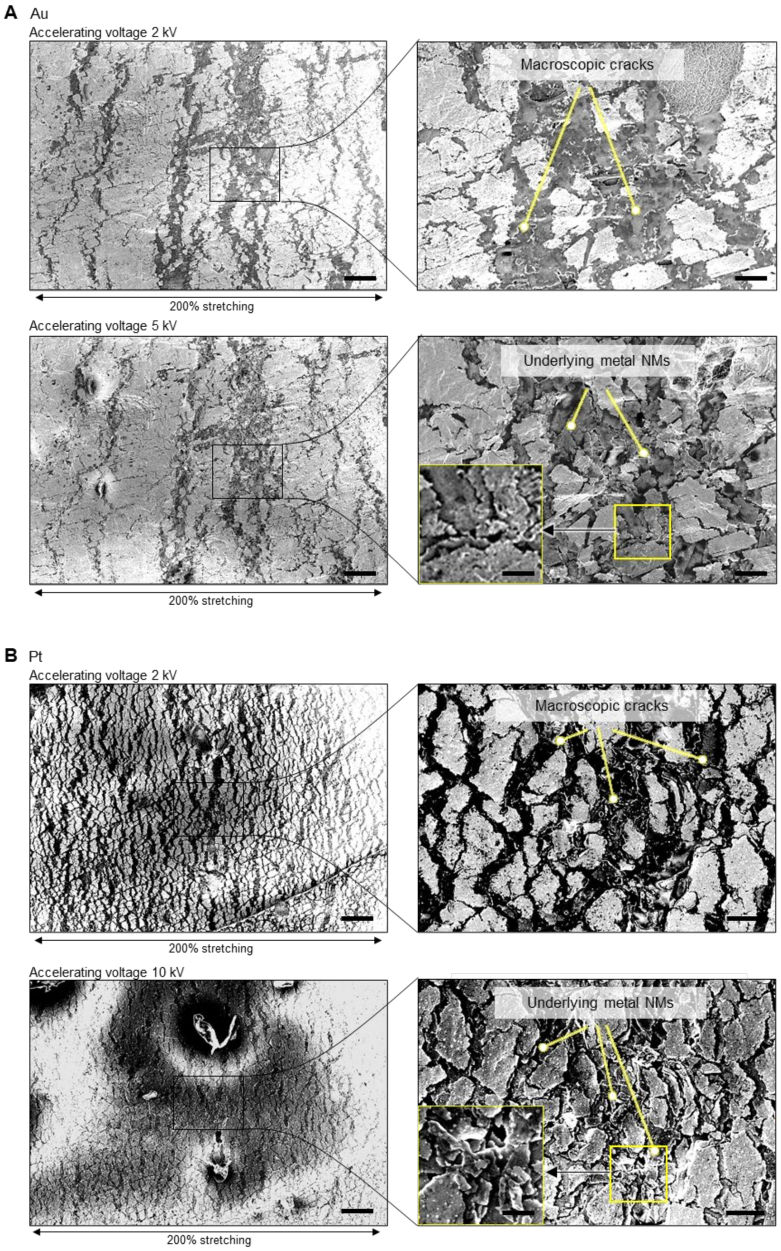


**Fig. S4. SEM characterization of crack-bridging pathways under 200% strain.** (**A**) SEM images of a 16-layer Au stack acquired at the same location with accelerating voltages of (top) 2 kV and (bottom) 5 kV, shown at (left) low and (right) high magnifications. (**B**) SEM images of a 16-layer Pt stack acquired at the same location with accelerating voltages of (top) 2 kV and (bottom) 10 kV, shown at (left) low and (right) high magnifications. Higher accelerating voltages reveal underlying metal layers through the cracks in the top layer, indicating vertical misalignment of cracks. Insets show magnified views of the representative regions marked with rectangles. Scale bars: (left) 50 μm, (right) 10 μm, (inset) 5 μm.

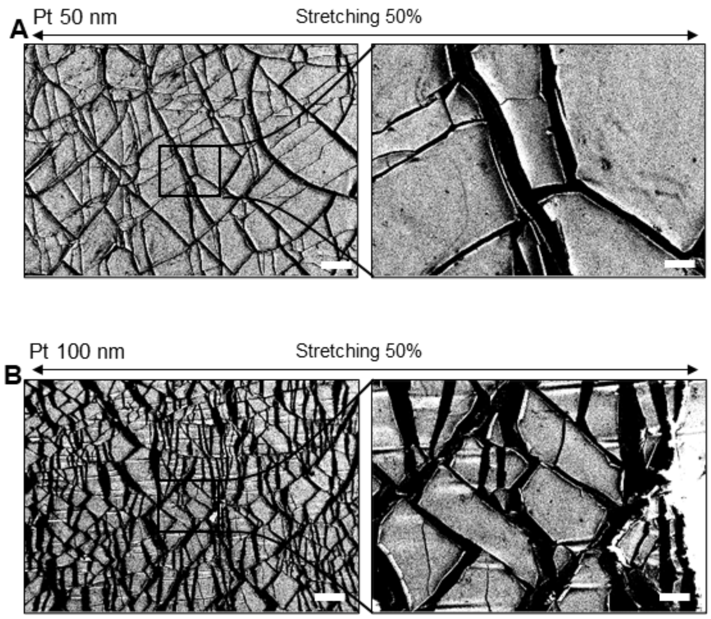


**Fig. S5. SEM analysis of Pt films under 50% strain.** (**A** and **B**) (left) Low- and (right) high-magnification SEM images of (A) 50 nm-thick and (B) 100 nm-thick Pt films deposited on poly(styrene-ethylene-butylene-styrene) (SEBS) rubber and stretched to 50% strain. High-magnification images correspond to regions indicated in the left panels. Scale bars: (left) 100 μm, (right) 20 μm.

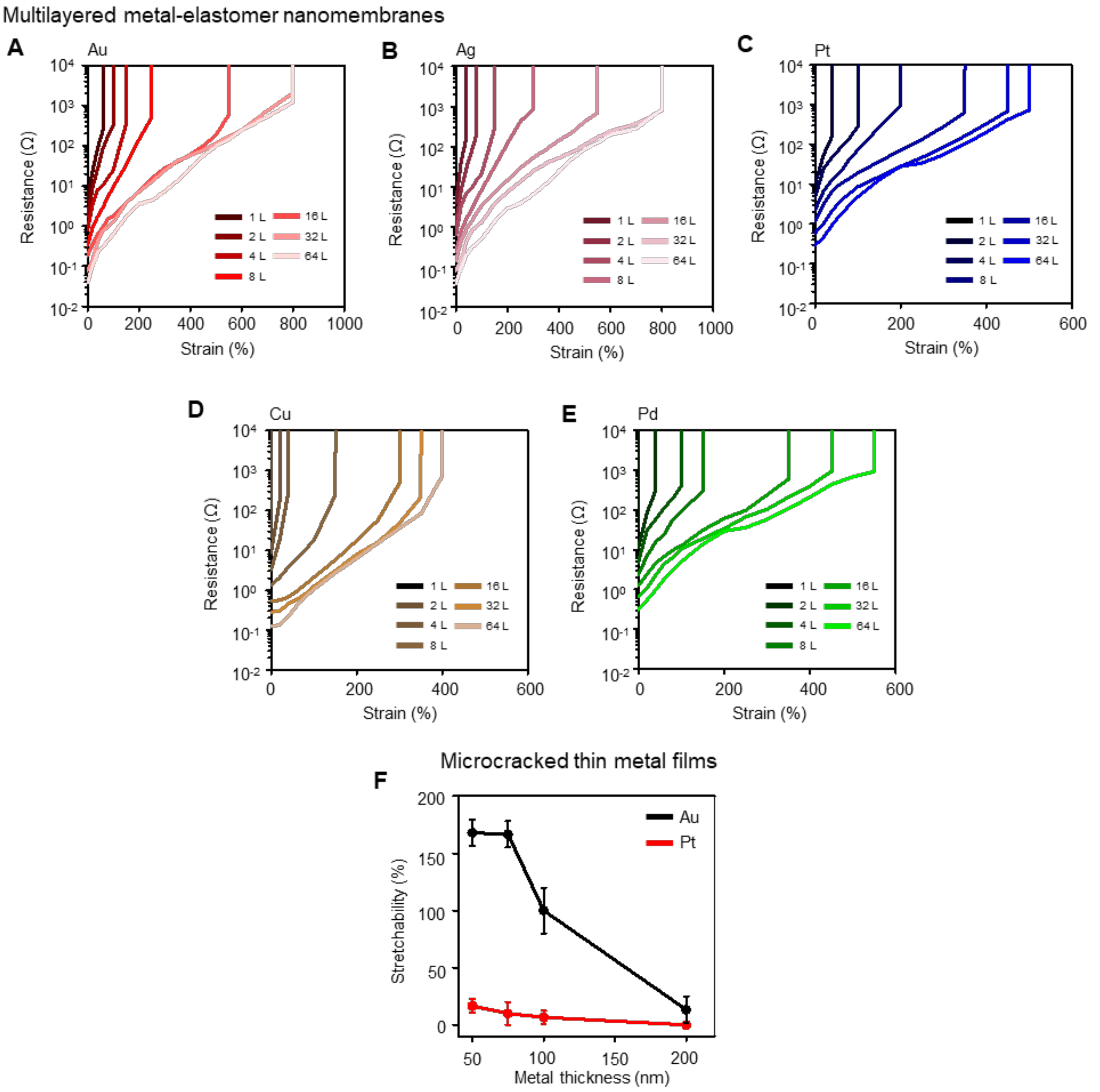


**Fig. S6. Strain-dependent electrical performance as a function of layer number across different metals and stretchability of thin-film-on-elastomer depending on metal thickness.** (**A** to **E**) Representative resistance curves under strain for multilayer stacks of (A) Au, (B) Ag, (C) Pt, (D) Cu, and (E) Pd as a function of the number of layers. (**F**) Stretchability of Au and Pt microcracked thin films depending on metal thickness. The films are fabricated by depositing them on SEBS elastomer. Markers and error bars represent mean ± s.d. (n = 3).

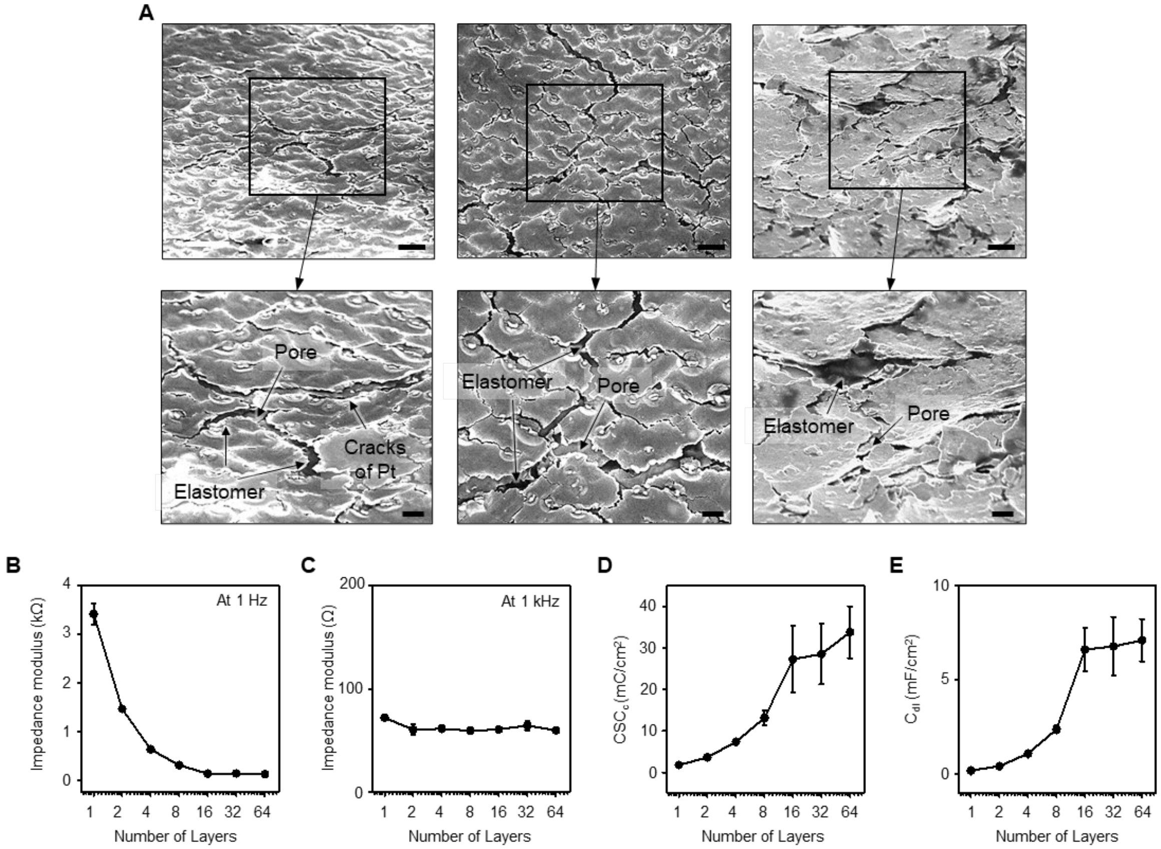


**Fig. S7. Electrochemical characterization of Pt multilayer stacks.** (**A**) Tilted-view SEM images (at 52°) showing the porous morphology of the elastomer NM beneath cracks in the Pt layer. Scale bars: (top) 5 µm, (bottom) 2 µm. (**B** to **D**) Impedance at (B) 1 Hz and (C) 1 kHz, and (D) CSCc of the platinum stacks versus the number of layers. Electrode size: 4 mm × 5 mm. (**E**) Double-layer capacitance ($C_{dl}$) estimated from EIS using a Randles-circuit model. Markers and error bars represent mean ± s.d. (n = 3).

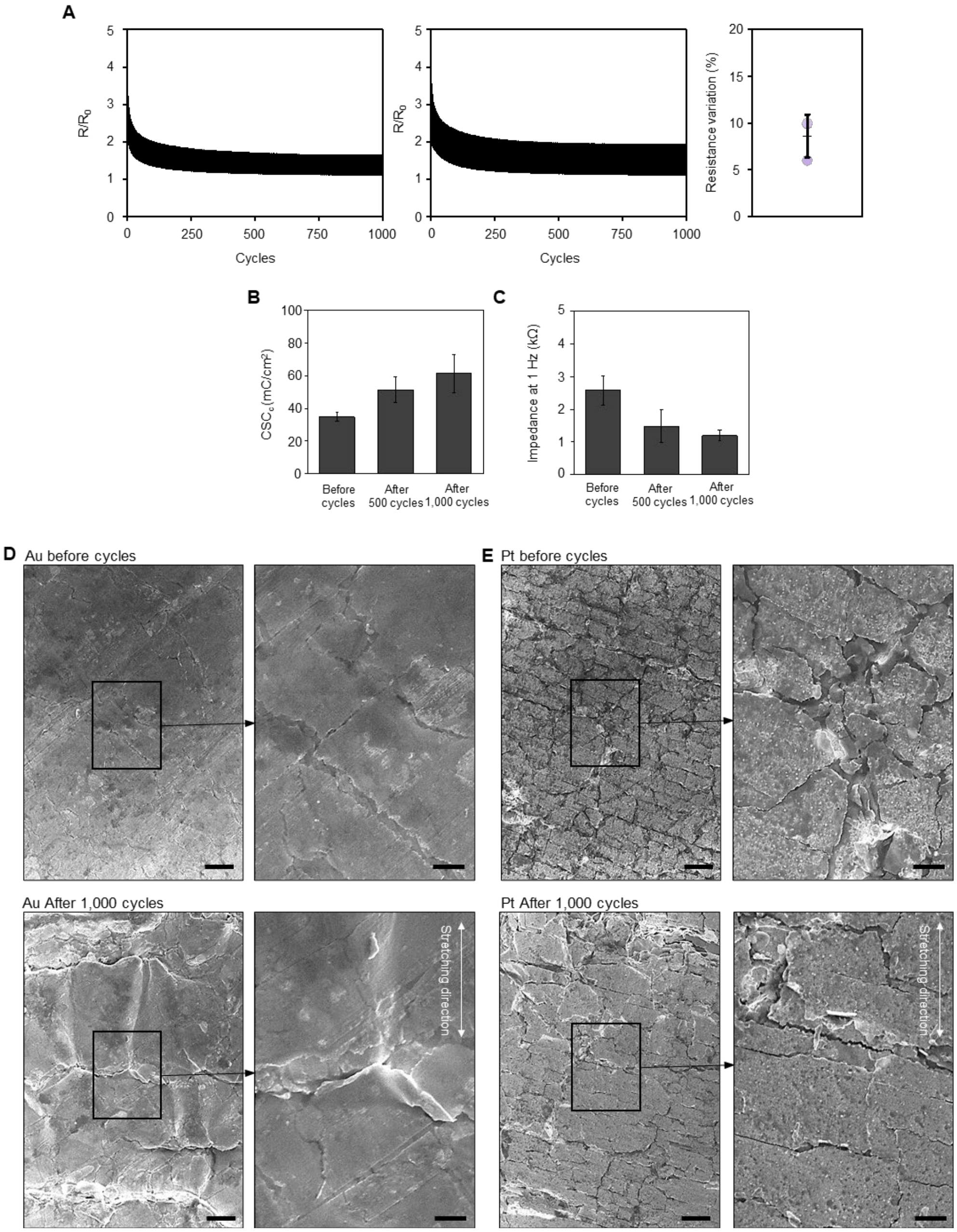


**Fig. S8. Electrical performance of 16-layer Au and Pt stacks under cyclic strain.** (**A**) Resistance variations of 16-layer Au stacks under 1,000 cycles of 30% strain. The right plot presents the average resistance variation and standard deviation after 1000 cycles. Bar and error bars represent mean ± s.d. (n = 3). (**B** and **C**) $CSC_c$ and impedance at 1 Hz after 0, 500, 1,000 cycles condition. Electrode size: 1 mm × 1.5 mm. Bars and error bars represent mean ± s.d. (n = 3). (**D**) SEM images of a 16-layer Au stack at (top) before and (bottom) after 1,000 cycles of 30% strain. Scale bars: (left) 20 µm, (right) 5 µm. (**E**) SEM images of a 16-layer Pt stack at (top) before and (bottom) after 1,000 cycles of 30% strain. Scale bars: (left) 20 µm, (right) 5 µm.

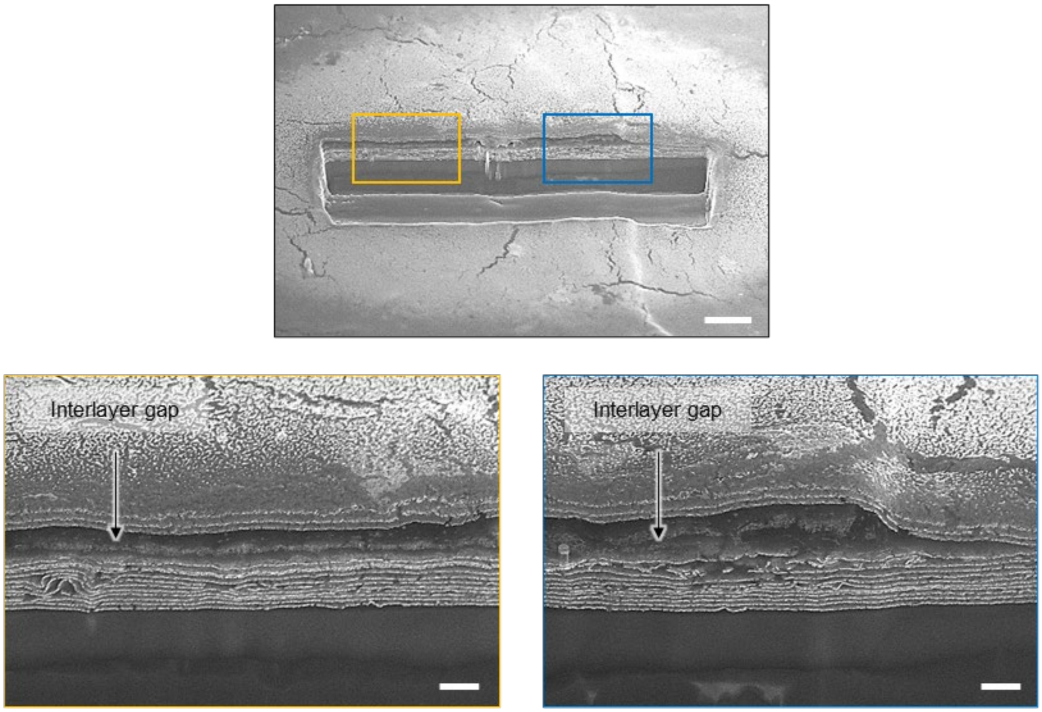


**Fig. S9. SEM analysis of interlayer gaps in a 16-layer Au stack with SEBS nanomembranes.** Cross-sectional SEM images at (top) low magnification and at (bottom) high magnification, collected from two locations in the top image. Scale bars: (top) 10 µm, (bottom) 2 µm.

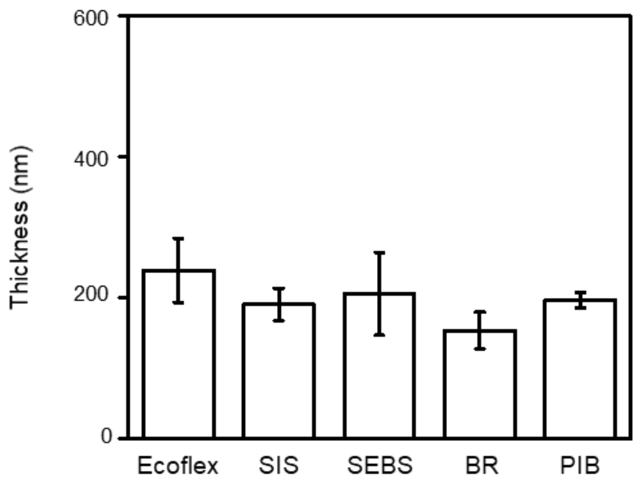


**Fig. S10. Thicknesses of different elastomer nanomembranes.** ~200 nm-thick films were obtained by adjusting the concentration of each elastomer in hexane, using the same spin-coating speed of 6,000 rpm. Bars and error bars represent mean ± s.d. (n = 3).

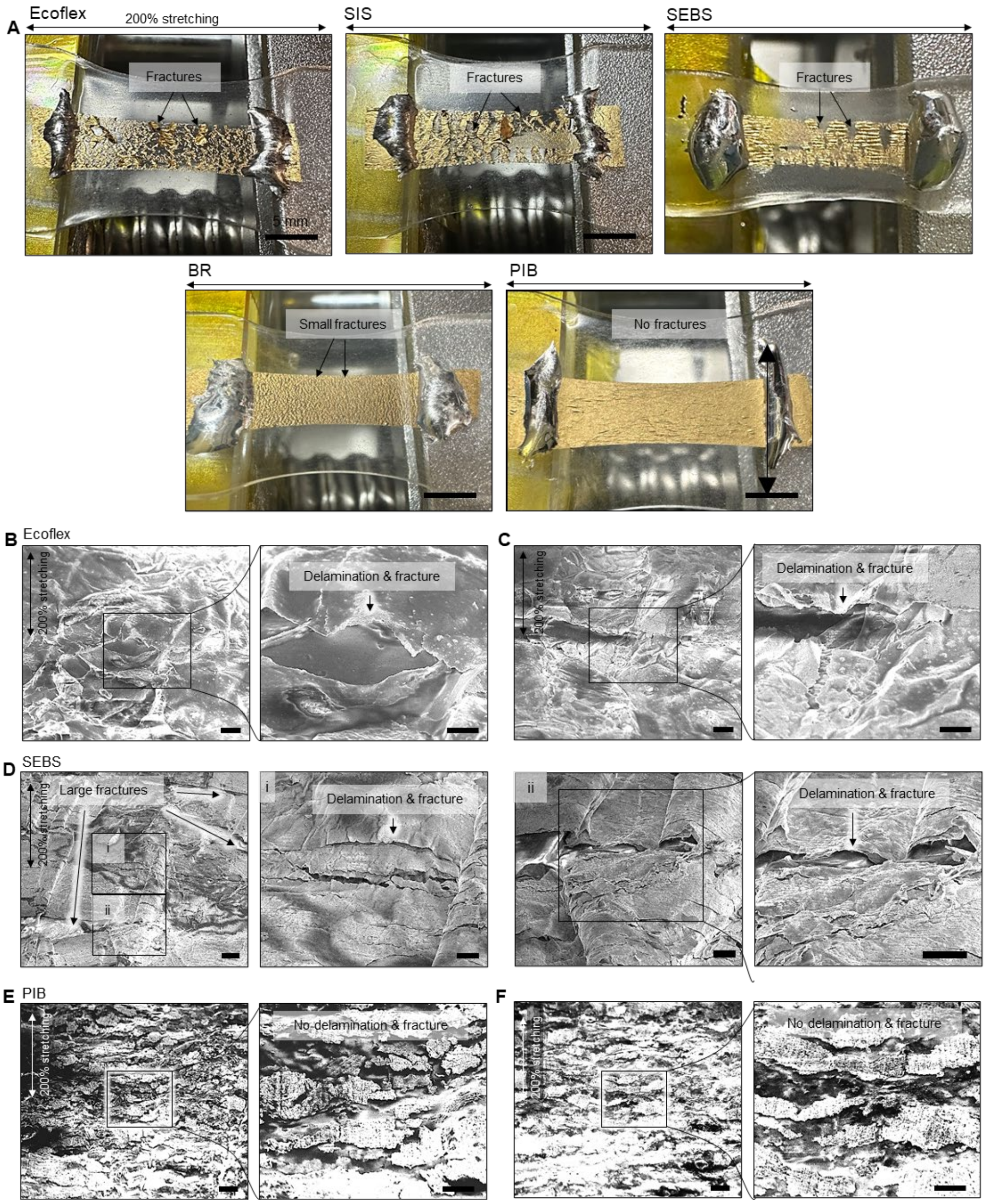


**Fig. S11. Structural integrity of 16-layer Au stacks using different elastomers under 200% strain.** (**A**) Photographs showing the stacks under deformation. (**B** and **C**) Tilted-view SEM images (at 52°) showing delamination and fracture in a 16-layer Au stack using Ecoflex. Scale bars: (left) 10 µm, (right) 5 µm. (**D**) Tilted-view SEM images showing delamination and fracture in a 16-layer Au stack using SEBS. Scale bars: (left) 20 µm, (i and ii) 5 µm, (ii, right) 5 µm. (**E** and **F**) Tilted-view SEM images showing high structural integrity in a 16-layer Au stack using PIB. Scale bars: (left) 10 µm, (right) 5 µm.

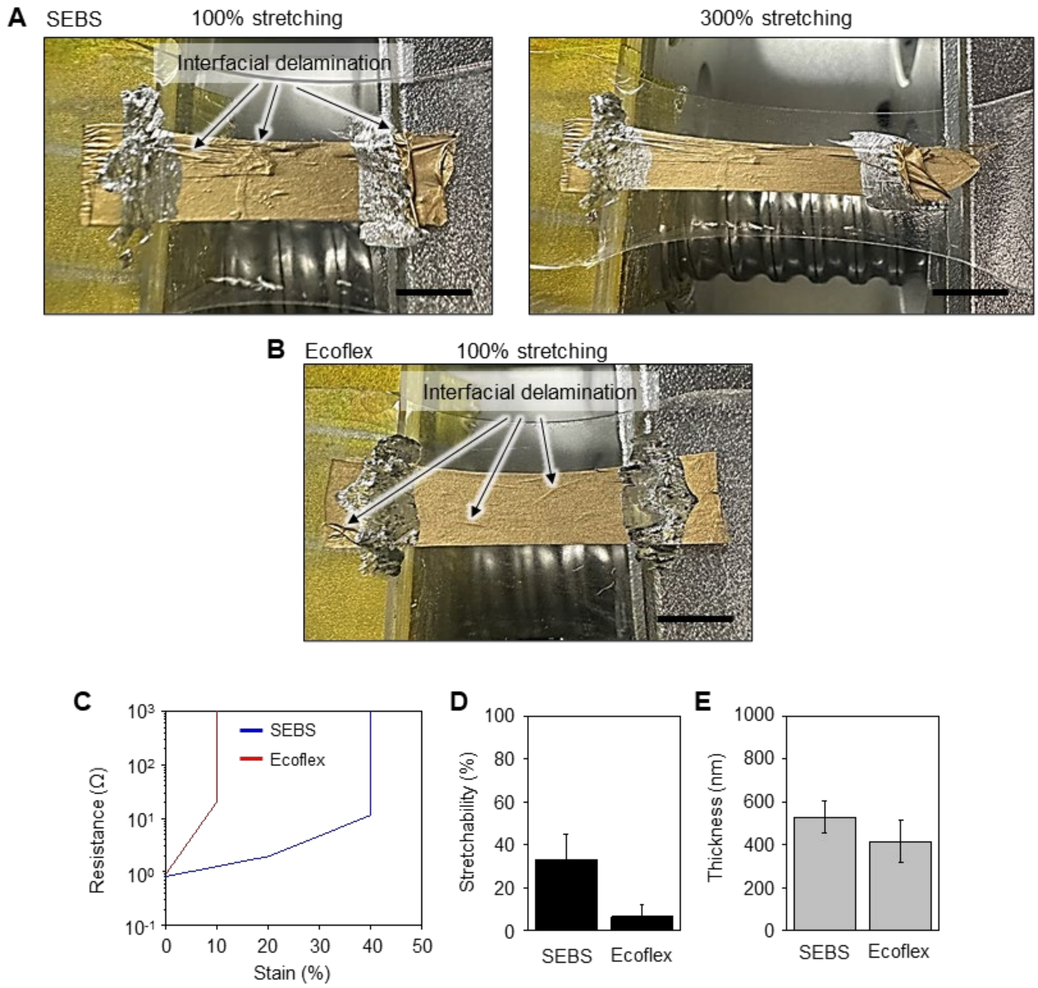


**Fig. S12. Structural integrity and stretchability of thicker elastomer nanomembranes.** (**A** and **B**) Photographs showing interfacial delamination in 16-layer Au stacks using (A) SEBS and (B) Ecoflex under deformation. SEBS and Ecoflex were used at concentrations of 0.075 wt% and 0.5 wt%, respectively. (**C**) Resistance of 16-layer Au stacks using SEBS and Ecoflex under strain. (**D**) Stretchability of 16-layer Au stacks using SEBS and Ecoflex. (**E**) Thickness of the SEBS and Ecoflex. (D and E) Bars and error bars represent mean ± s.d. (n = 3).

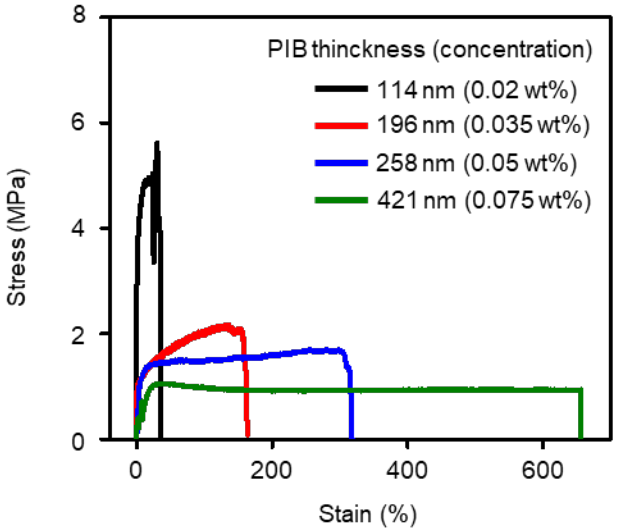


**Fig. S13. Stress-strain curves of 16-layer Au stacks using PIB NMs with varying thicknesses.**

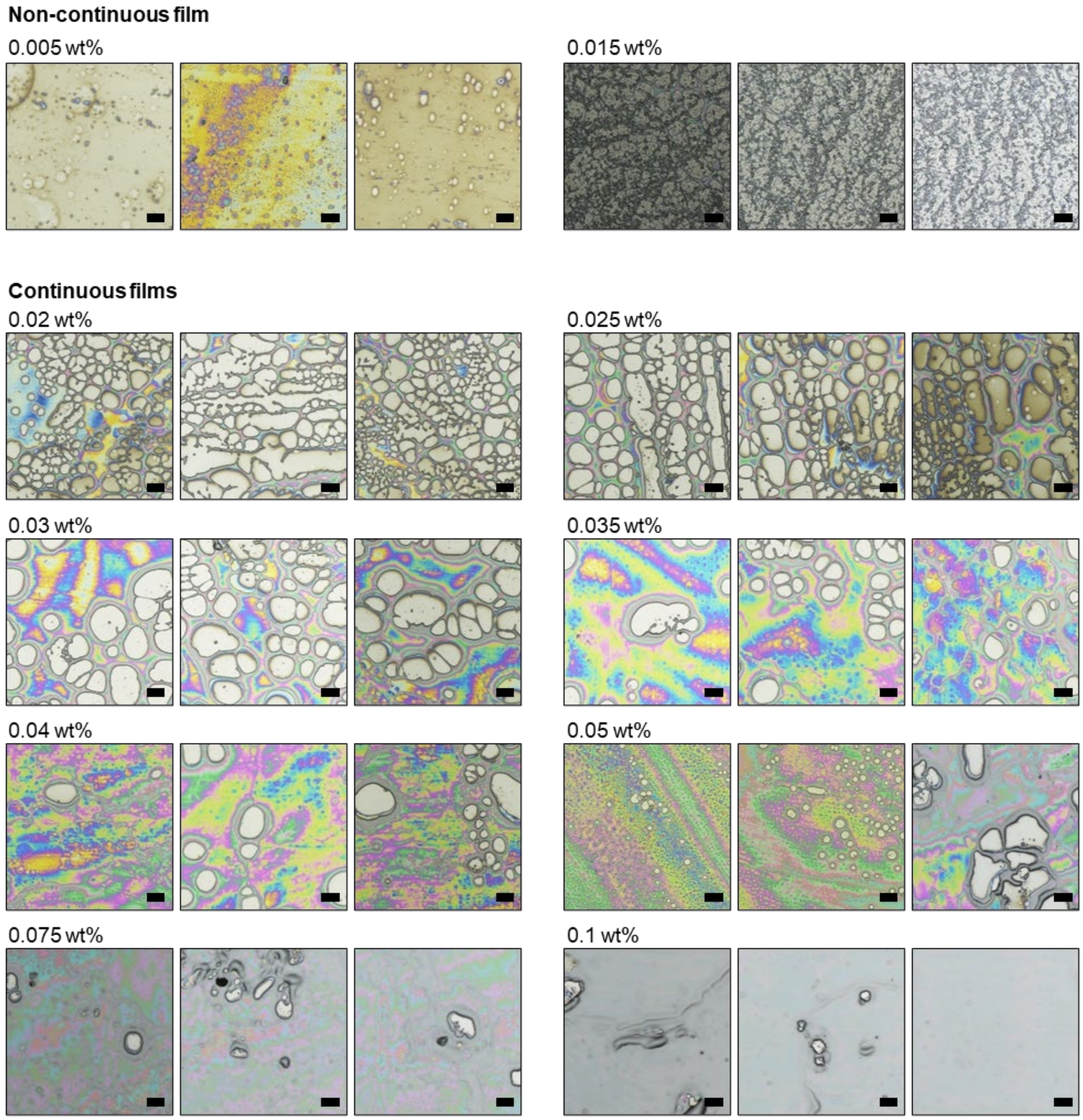


**Fig. S14. Porosity variation in PIB nanomembranes as a function of the PIB concentration in hexane prior to spin coating.** Brightfield microscope images from three different locations at each concentration showing porosity differences. All scale bars, 20 µm.

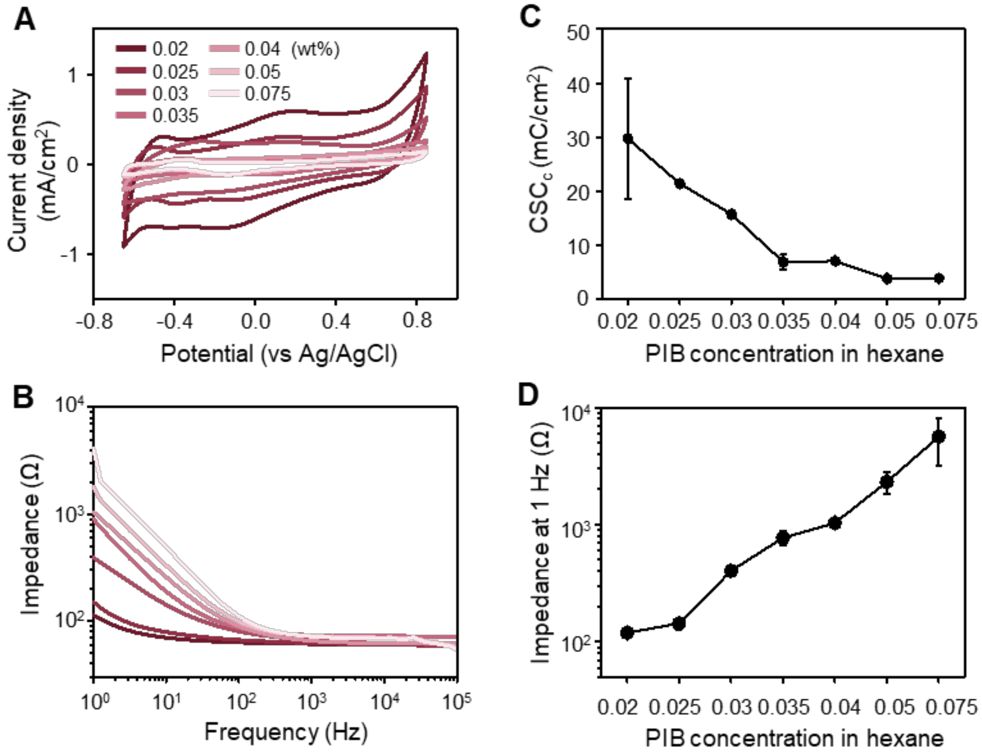


**Fig. S15. Electrochemical performance of 16-layer Pt stacks using PIB nanomembranes with varying thicknesses.** (**A** and **B**) Representative (A) cyclic voltammetry and (B) impedance spectra curves of 16-layer Pt stacks using PIB nanomembranes with different concentrations. (**C** and **D**) (C) $CSC_c$ and (D) impedance at 1 Hz of 16-layer Pt stacks using PIB nanomembranes with different concentrations. Electrode size 4 × 5 $mm^2$. Markers and error bars represent mean ± s.d. (n = 3).

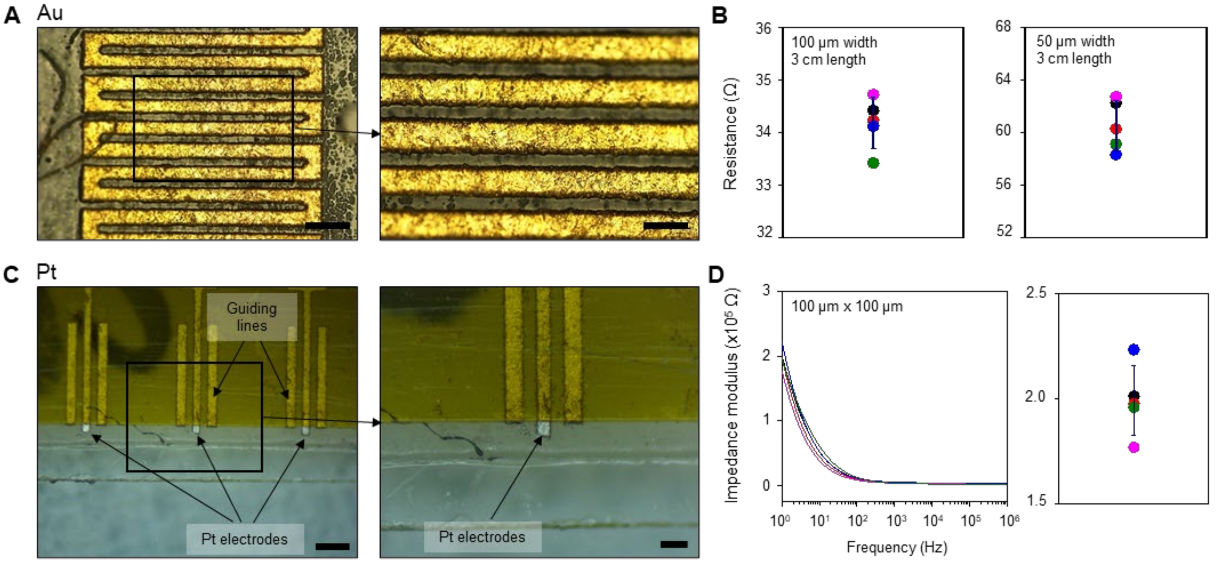


**Fig. S16. Electrical performance of Au nanomembranes traces and Pt nanomembranes electrodes produced by UV laser patterning.** (**A**) Microscope images showing clear line patterns with a width of 50 µm in 16-layer Au stacks after UV laser patterning. Scale bars: (left) 200 µm, (right) 100 µm. (**B**) Resistance of five patterned lines in 16-layer Au stacks with widths of (left) 100 µm and (right) 50 µm, and a length of 3 cm. (**C**) Microscope images showing electrode patterns with a width and length of 100 µm in 16-layer Pt stacks after UV laser patterning. Scale bars: (left) 500 µm, (right) 200 µm. (**D**) (left) Impedance spectra and (right) impedance at 1 Hz of five patterned electrodes with a width and length of 100 µm from 16-layer Pt stacks. (B and D) Markers and error bars represent mean ± s.d. (n = 5).

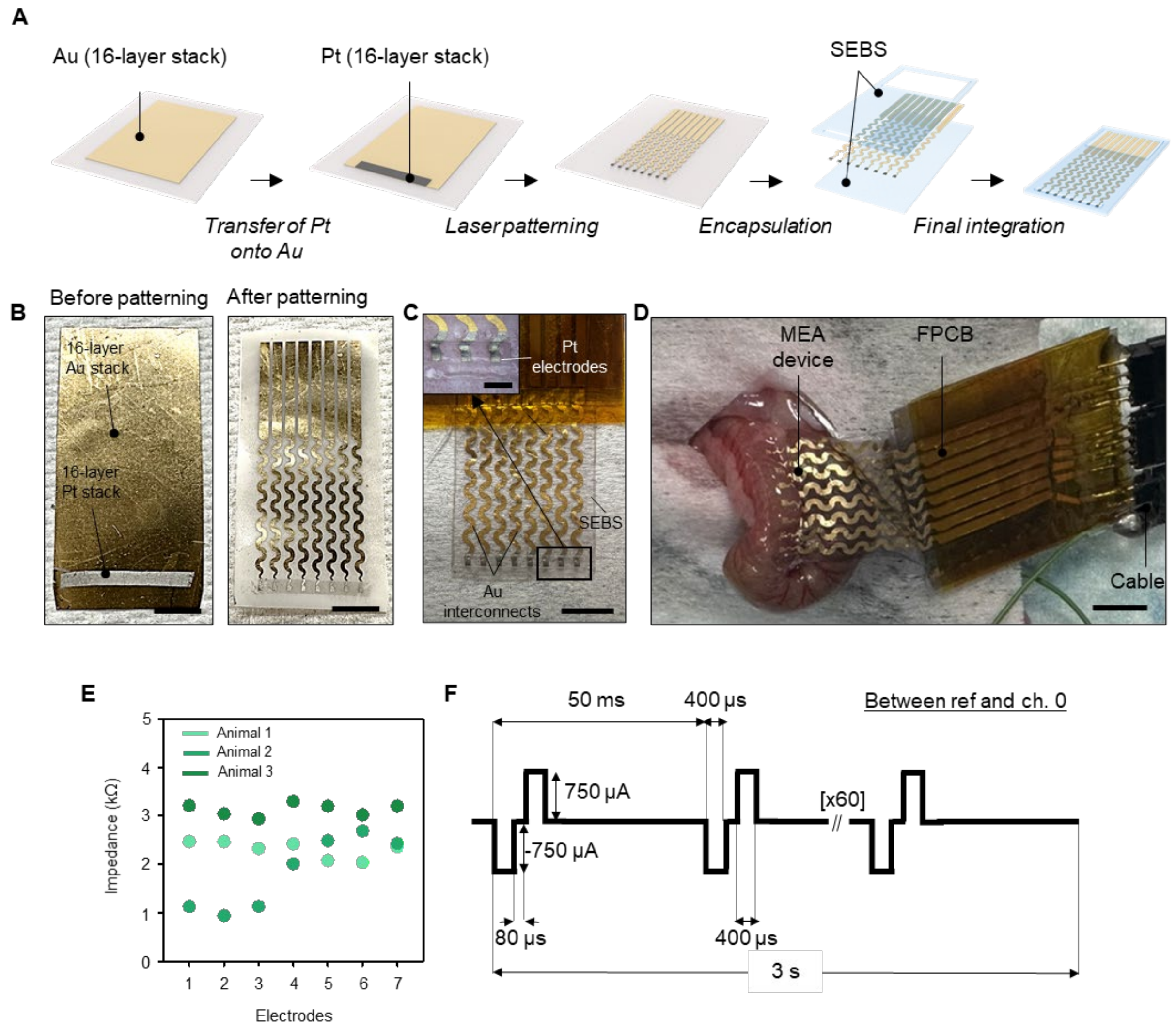


**Fig. S17. Fabrication and in vivo application of the nanomembrane electrode arrays.** (**A**) An illustration of the fabrication process. (**B**) Photographs of nanomembranes before and after patterning. Scale bars: 5 mm. (**C**) A photograph of a nanomembrane electrode array. Scale bars: 5 mm, (inset) 1.5 mm. (**D**) A photograph of the experimental setup used for recordings of murine colonic EMG activity. The stretchable nanomembrane electrode array is connected to the flexible printed circuit board (FPCB), which interfaces with the electrophysiology recording system (Ripple) via a cable. Scale bar: 1 cm. (**E**) Impedance values (at 1 kHz) of seven individual electrodes measured across three different mice. (**F**) Parameters of the electrical pulse burst used for stimulation.

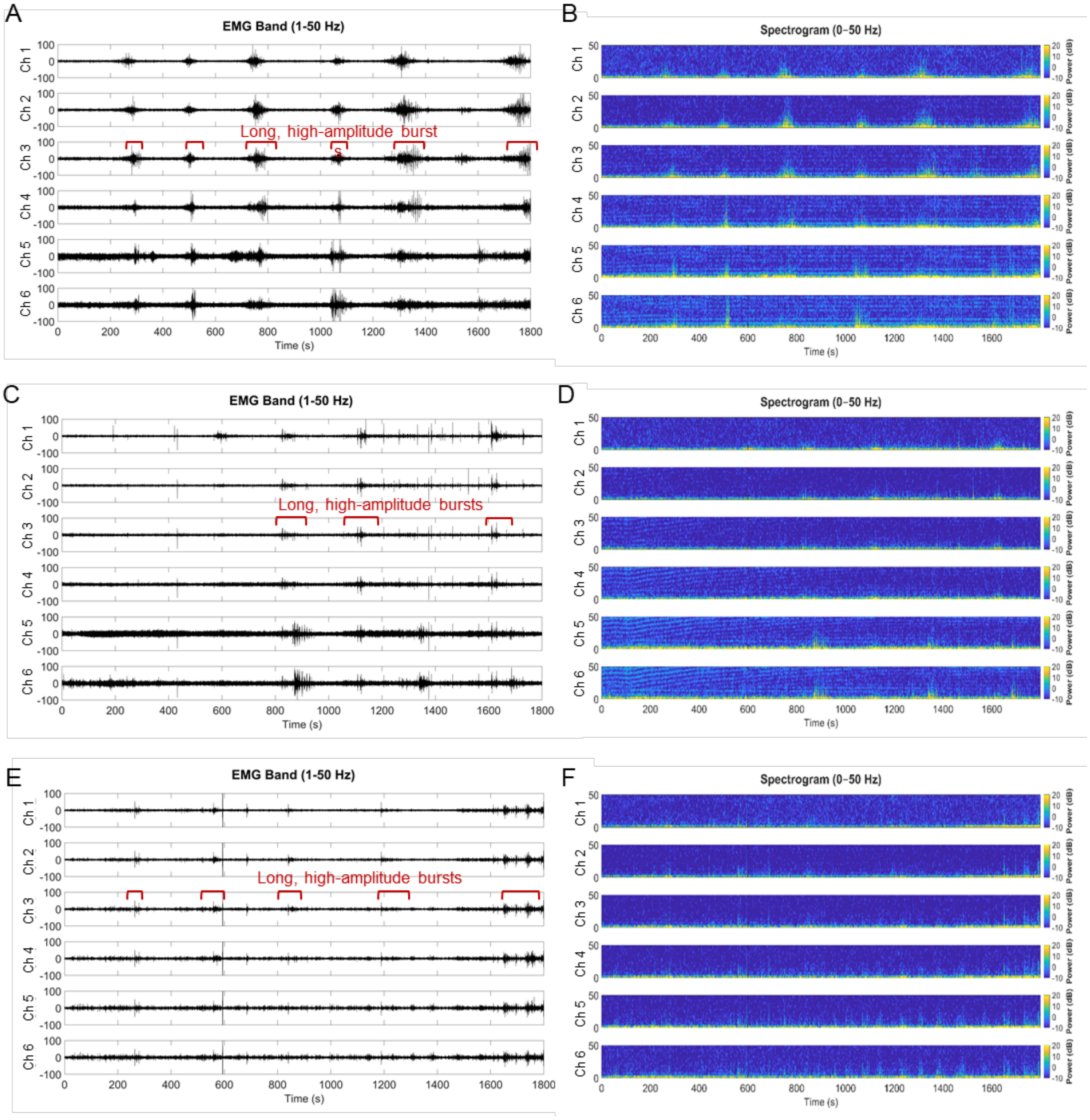


**Fig. S18. Spontaneous electrophysiological responses across channels 1–6 in three animals.** (**A, C, E**) EMG-band (1–50 Hz) signals and (**B, D, F**) corresponding EMG spectrograms in animals 1, 2, and 3, respectively.

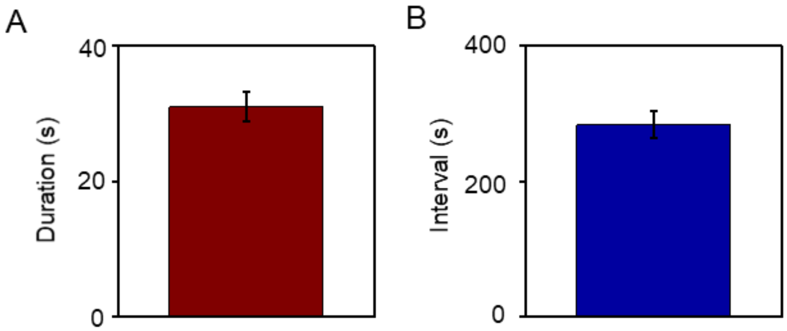


**Fig. S19. (A) Duration and (B) Interval of neurogenic bursts.** Bars and error bars represent mean ± s.e.m. (n=101)

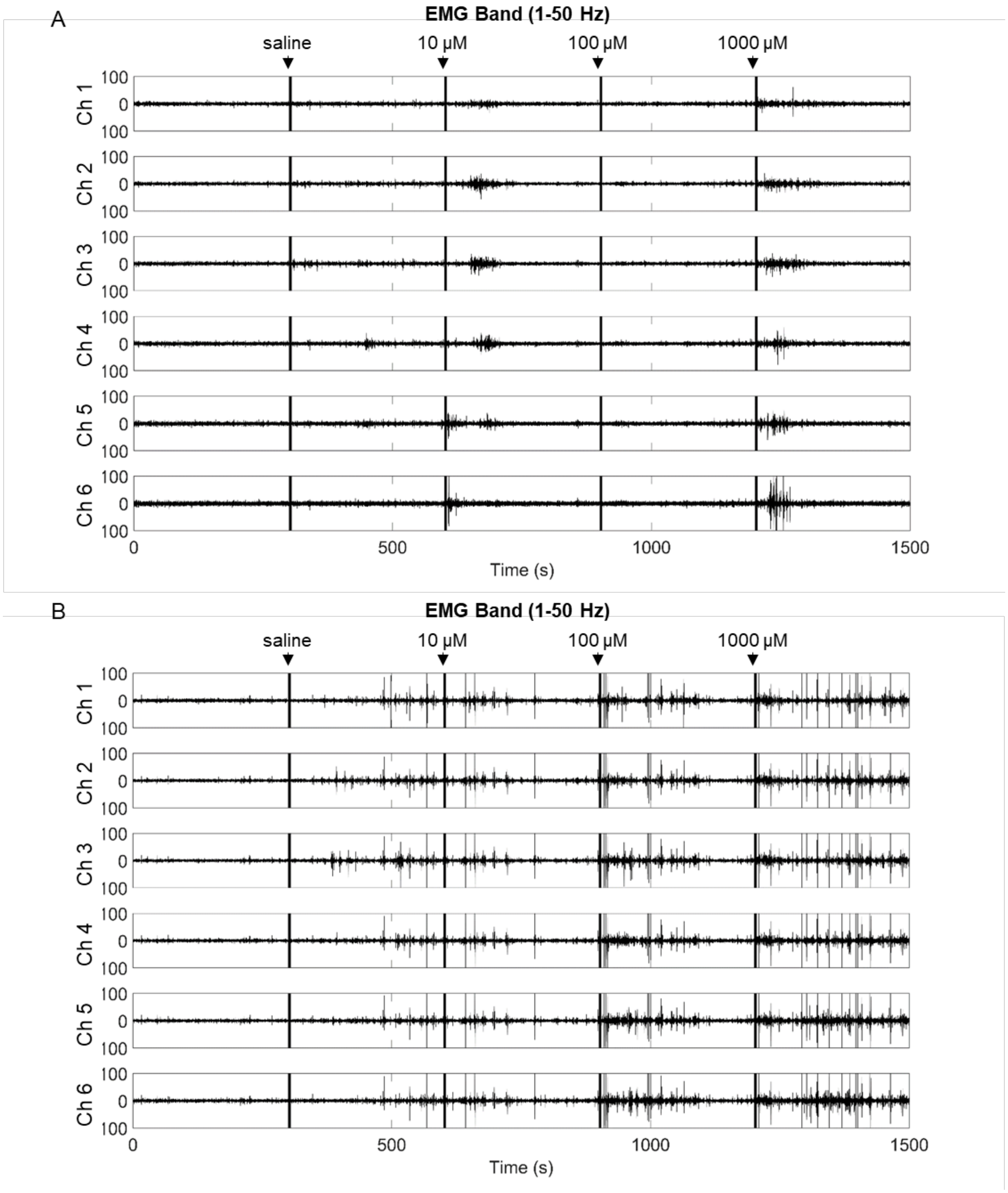


**Fig. S20. Electrophysiological responses to saline and varying concentrations of bethanechol across channels 1–6 in two animals.** (**A**) EMG-band (1–50 Hz) signals from Animal 1. (**B**) EMG-band (1–50 Hz) signals from Animal 2. Black bars indicate successive additions of 20 µL saline followed by 20 µL of 10 µM, 100 µM, and 1 mM bethanechol.

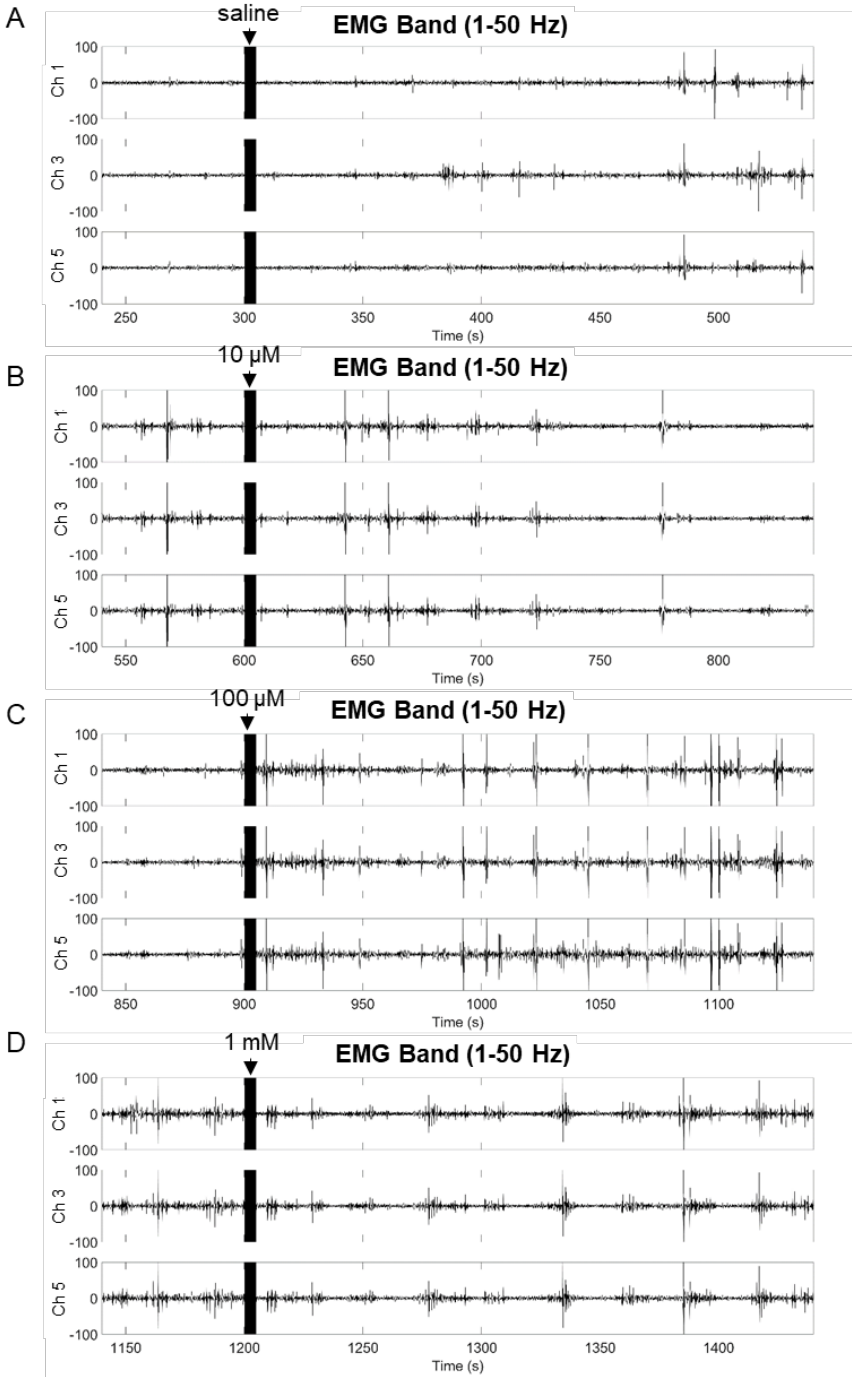


**Fig. S21. Electrophysiological responses to saline and varying concentration of bethanechol across channels 1, 3, and 5 in Animal 2.** EMG-band (1–50 Hz) signals in response to (**A**) 20 µL of saline, (**B**) 20 µL of 10 µM, (**C**) 100 µM, and (**D**) 1 mM bethanechol. Black bars indicate the times of addition.

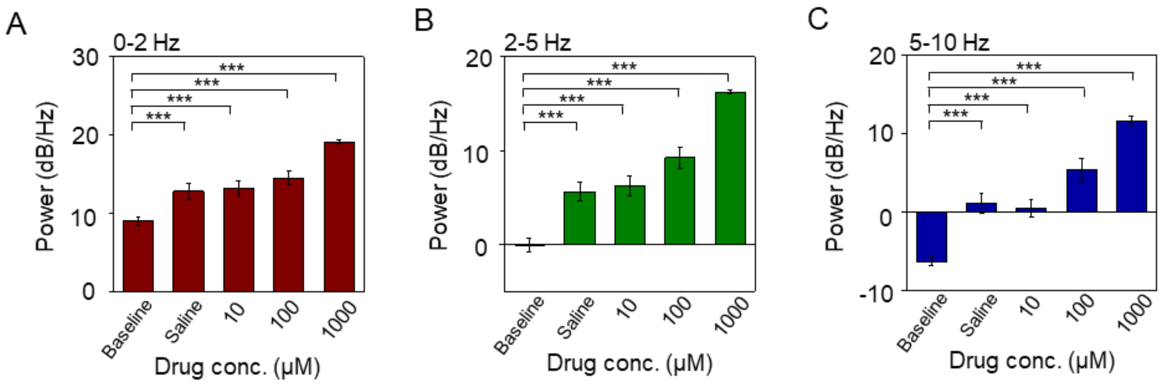


**Fig. S22. Power analysis of electrophysiological responses to saline and varying concentrations of bethanechol.** Averaged power in the (**A**) 0–2 Hz, (**B**) 2–5 Hz, and (**C**) 5–10 Hz frequency bands comparing responses to additions of 20 µL saline, and 20 µL of 10 µM, 100 µM, and 1 mM bethanechol. ***$p < 0.001$. Bars and error bars denote mean ± s.e.m. ($n = 12$).

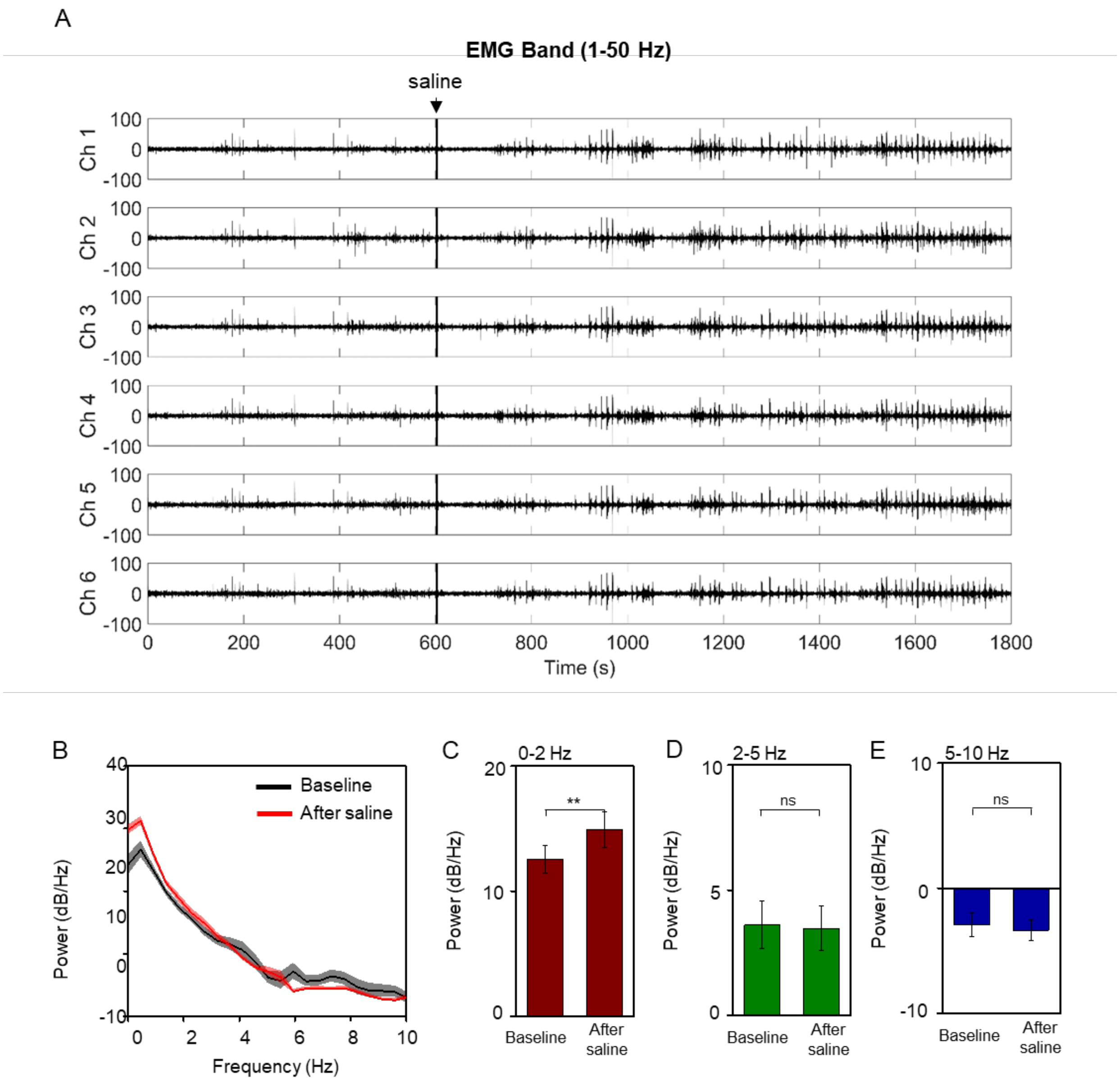


**Fig. S23. Electrophysiological responses to saline addition in two animals.** (**A**) EMG-band (1–50 Hz) signals across channels 1–6 in animal 1. (**B**) Power spectral analysis showing no significant differences before and after saline addition in animals 1 and 2. Averaged power in the (**C**) 0–2 Hz, (**D**) 2–5 Hz, and (**E**) 5–10 Hz frequency bands before and after saline addition. The black bar indicates the timing of 100 µL saline delivery. ** $p < 0.01$; ns, not significant ($p > 0.05$). Bars and error bars denote mean ± s.e.m. ($n = 12$).

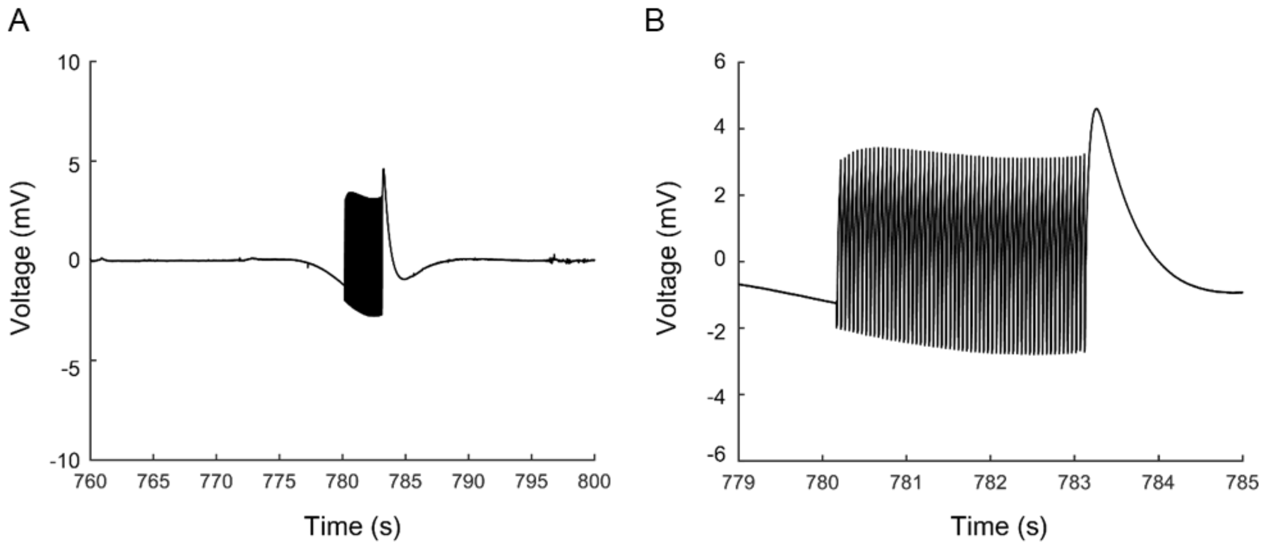


**Fig. S24. Stimulation artifact.** (**A**) Raw electrophysiological signal during electrical stimulation, showing the stimulation artifacts associated with pulse bursts from channel 1 in Animal 3. (**B**) Zoom-in on the first pulse burst artifact.

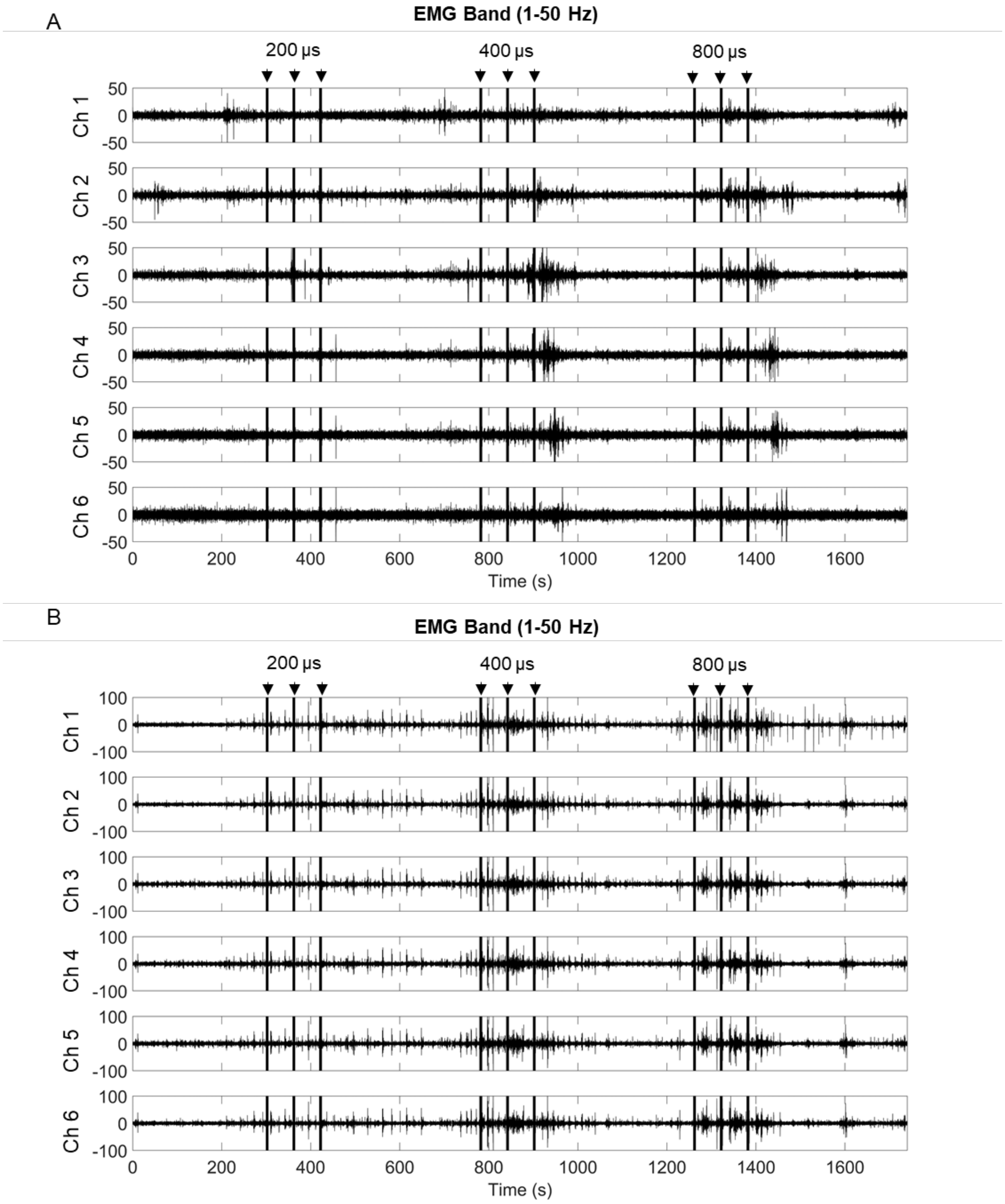


**Fig. S25. Electrophysiological responses across channels 1–6 to pulse-width variation in two animals.** (**A**) Animal 1; (**B**) Animal 2. Traces show EMG-band (1–50 Hz) signals. Each condition comprised three repetitions of 3 s trains of 60 bipolar pulses at 750 μA with pulse widths of 200, 400, or 800 μs. Black bars indicate stimulation epochs.

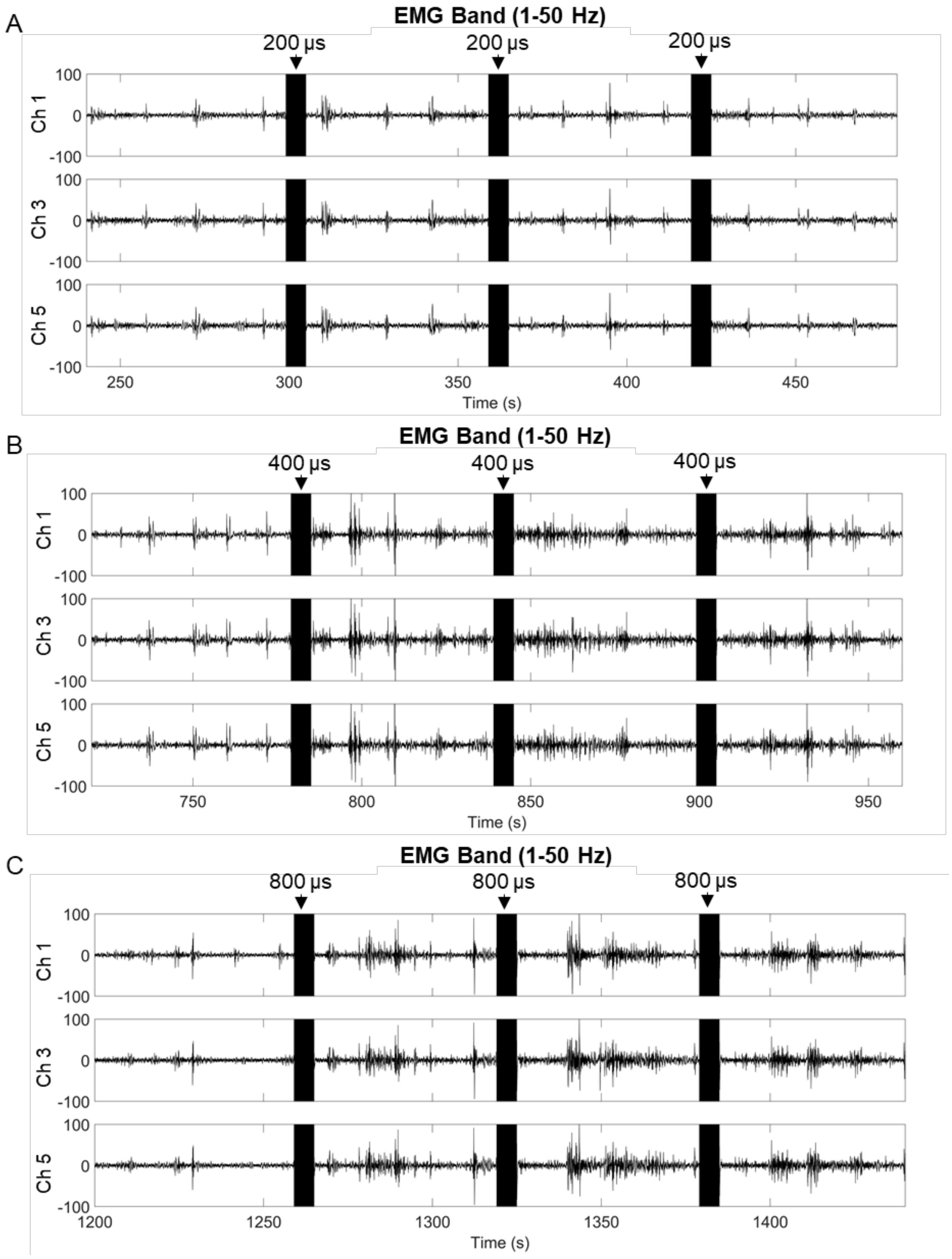


**Fig. S26. Electrophysiological responses on channels 1, 3, and 5 in Animal 2 under varying pulse widths.** EMG-band (1–50 Hz) traces are shown for three repetitions of 3 s trains of 60 biphasic pulses at 750 µA with pulse widths of (A) 200 µs, (B) 400 µs, and (C) 800 µs. Black bars denote stimulation epochs.

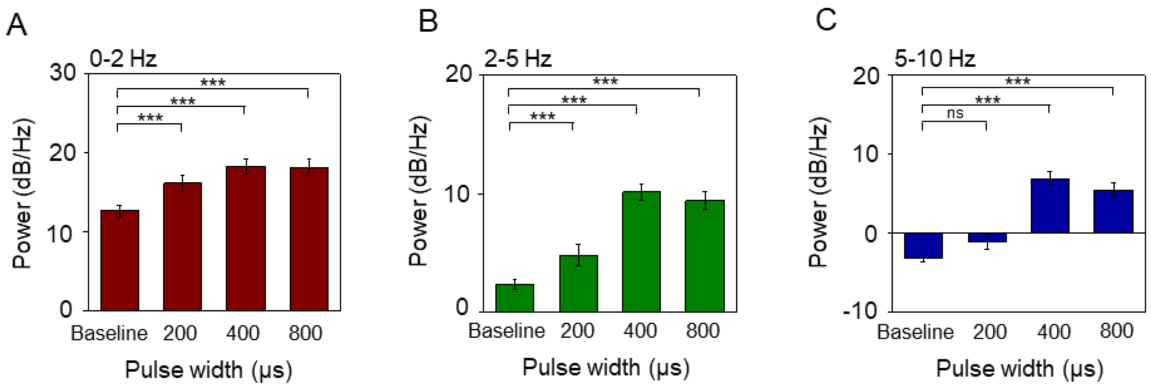


**Fig. S27. Power analysis of electrophysiological responses with varying pulse widths.** Averaged power in the (**A**) 0–2 Hz, (**B**) 2–5 Hz, and (**C**) 5–10 Hz frequency bands comparing baseline and responses to three repetitions of 60 biphasic pulses with pulse widths of 200 µs, 400 µs, and 800 µs. ns, not significant ($p > 0.05$); ***$p < 0.001$. Bars and error bars denote mean ± s.e.m. ($n = 36$).

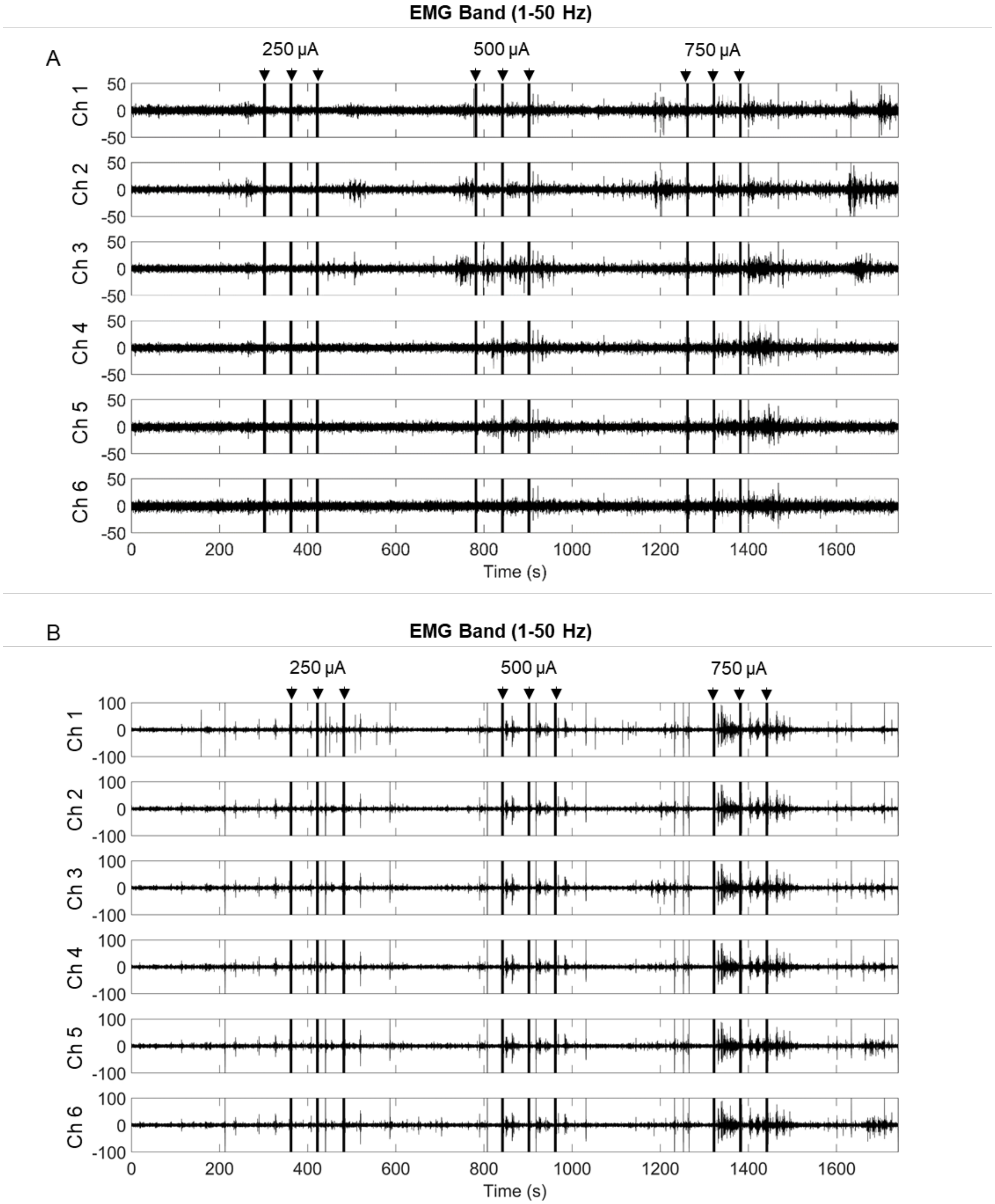


**Fig. S28. Electrophysiological responses across channels 1–6 under varying pulse amplitudes in two animals.** (**A**) EMG-band (1–50 Hz) signals in response to pulse bursts in Animal 1 (**B**) EMG-band (1–50 Hz) signals in response to pulse bursts in Animal 2. Each condition comprised three repetitions of 3 s trains of 60 bipolar pulses (400 µs pulse width, 80 µs interpulse interval) with amplitudes of 250, 500, and 750 µA. Black bars denote stimulation epochs.

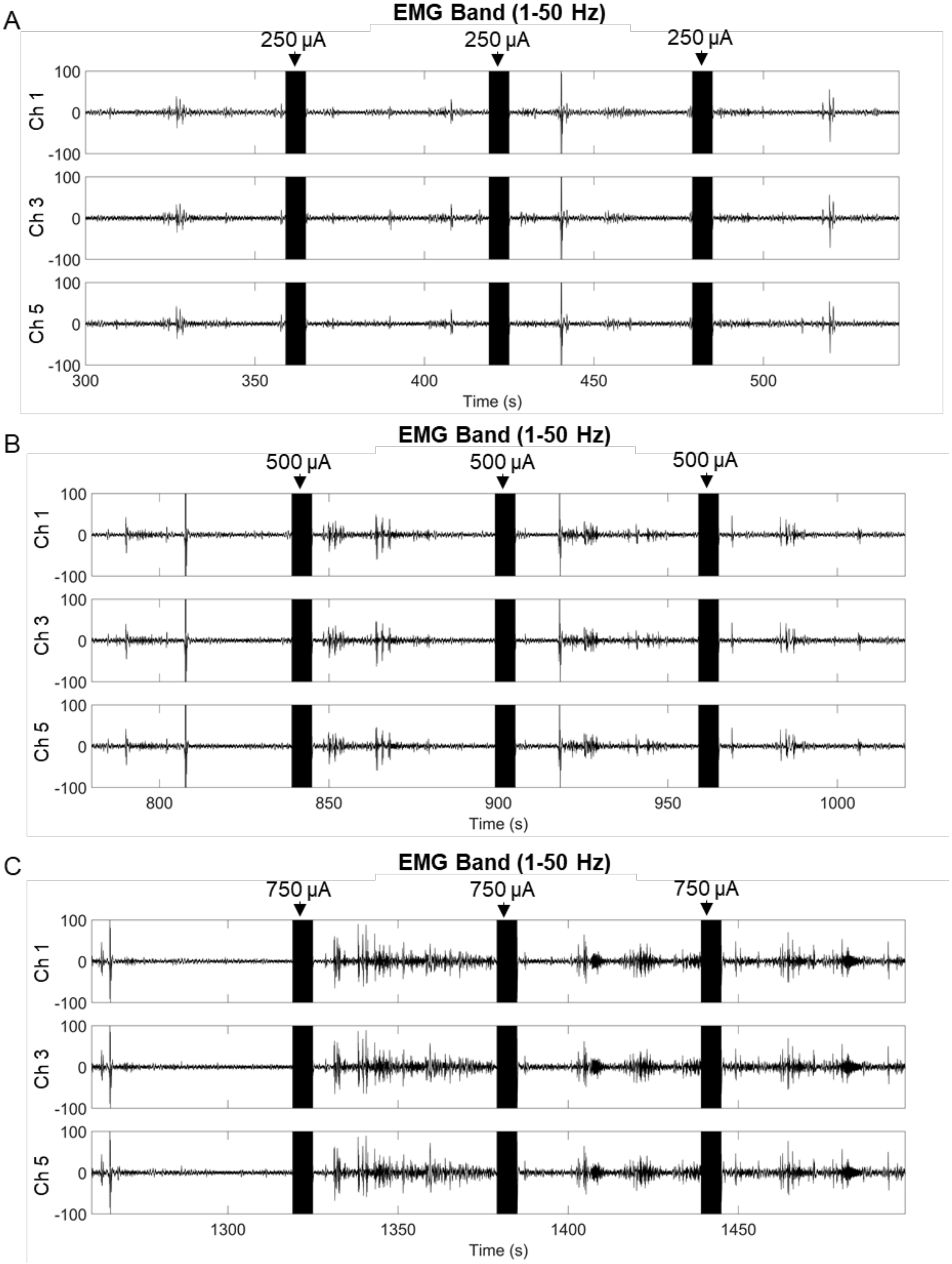


**Fig. S29. Electrophysiological responses across channels 1, 3, 5 with varying pulse amplitudes in Animal 2.** EMG-band (1–50 Hz) signals in response to 3 repetitions of 60 biphasic pulses with pulse amplitudes of (A) 250 µA, (B) 500 µA, and (C) 750 µA. Black bars indicate stimulation epochs.

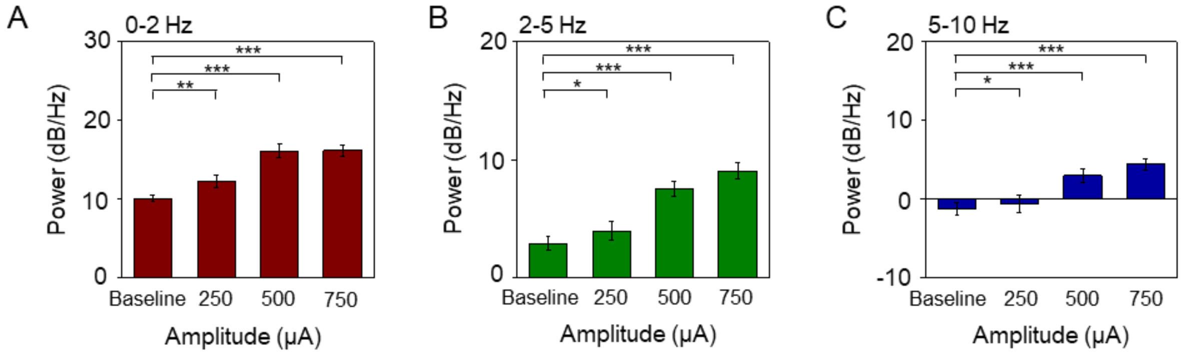


**Fig. S30. Power analysis of electrophysiological responses with varying pulse amplitudes.** Averaged power in the (**A**) 0–2 Hz, (**B**) 2–5 Hz, and (**C**) 5–10 Hz frequency bands comparing baseline and responses to three repetitions of 60 biphasic pulses with pulse amplitudes of 250 μA, 500 μA, and 750 μA. *$p < 0.05$; **$p < 0.01$; ***$p < 0.001$. Bars and error bars denote mean ± s.e.m. ($n = 36$).

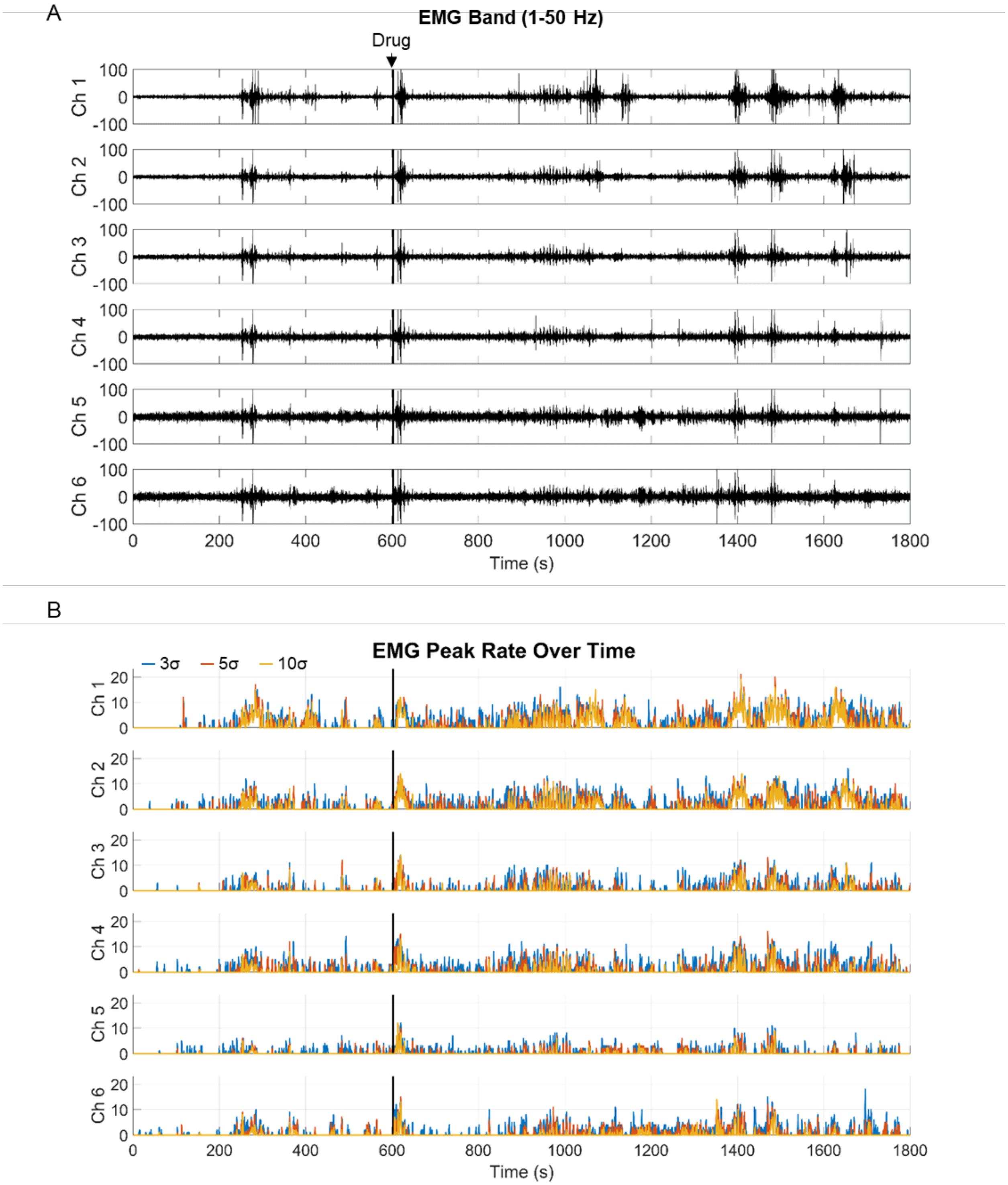


**Fig. S31. Electrophysiological recordings during pharmacological stimulation in Animal 3 (channels 1–6).** (**A**) EMG-band (1–50 Hz) signals before and after drug injection. (**B**) EMG peak rate across time. Black region marks drug delivery.

A
B
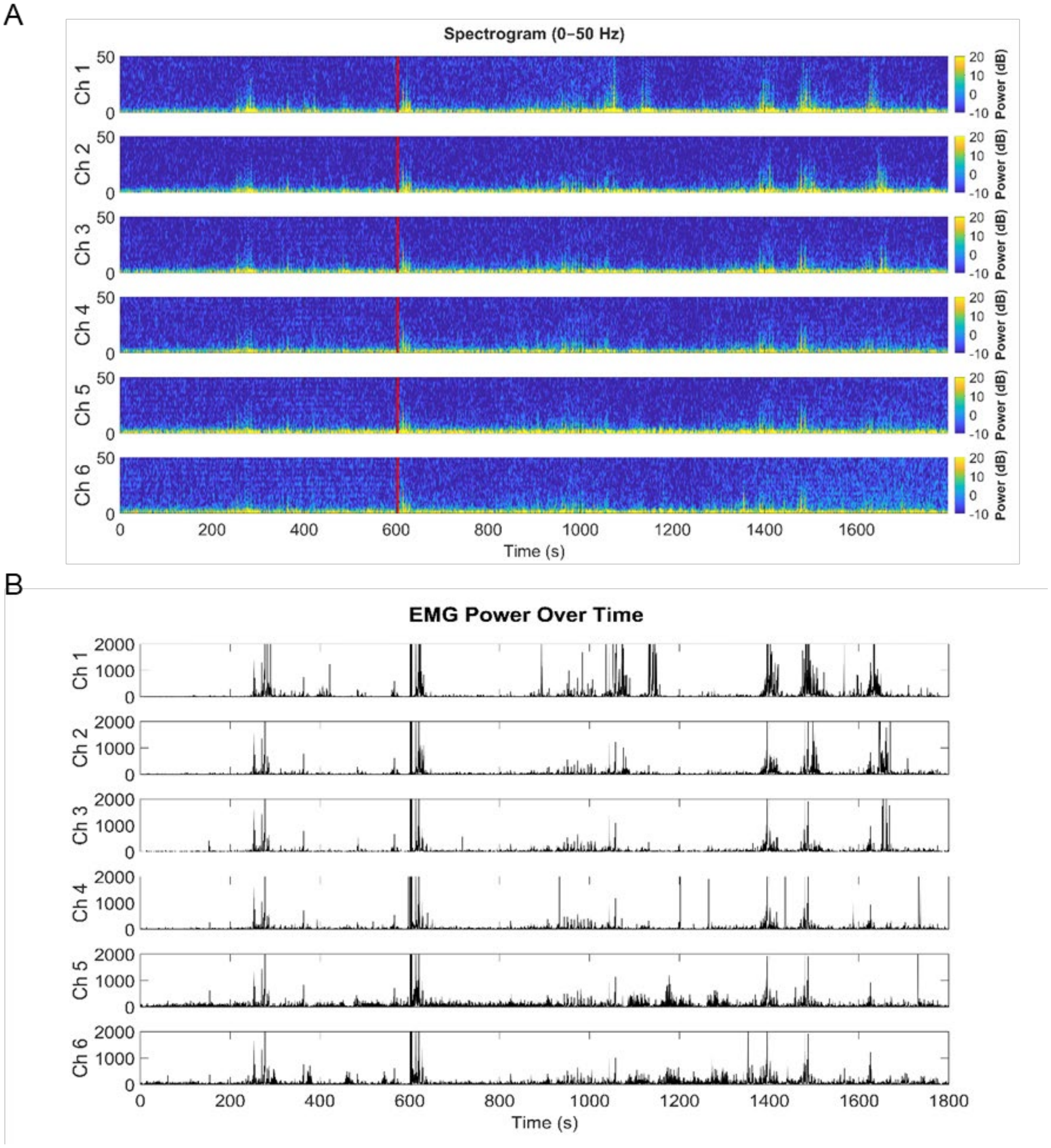


**Fig. S32. Analysis of pharmacological stimulation in Animal 3 (channels 1–6).** (**A**) EMG-band spectrogram (0–50 Hz). (**B**) EMG power over time. Red and black lines mark injection time.

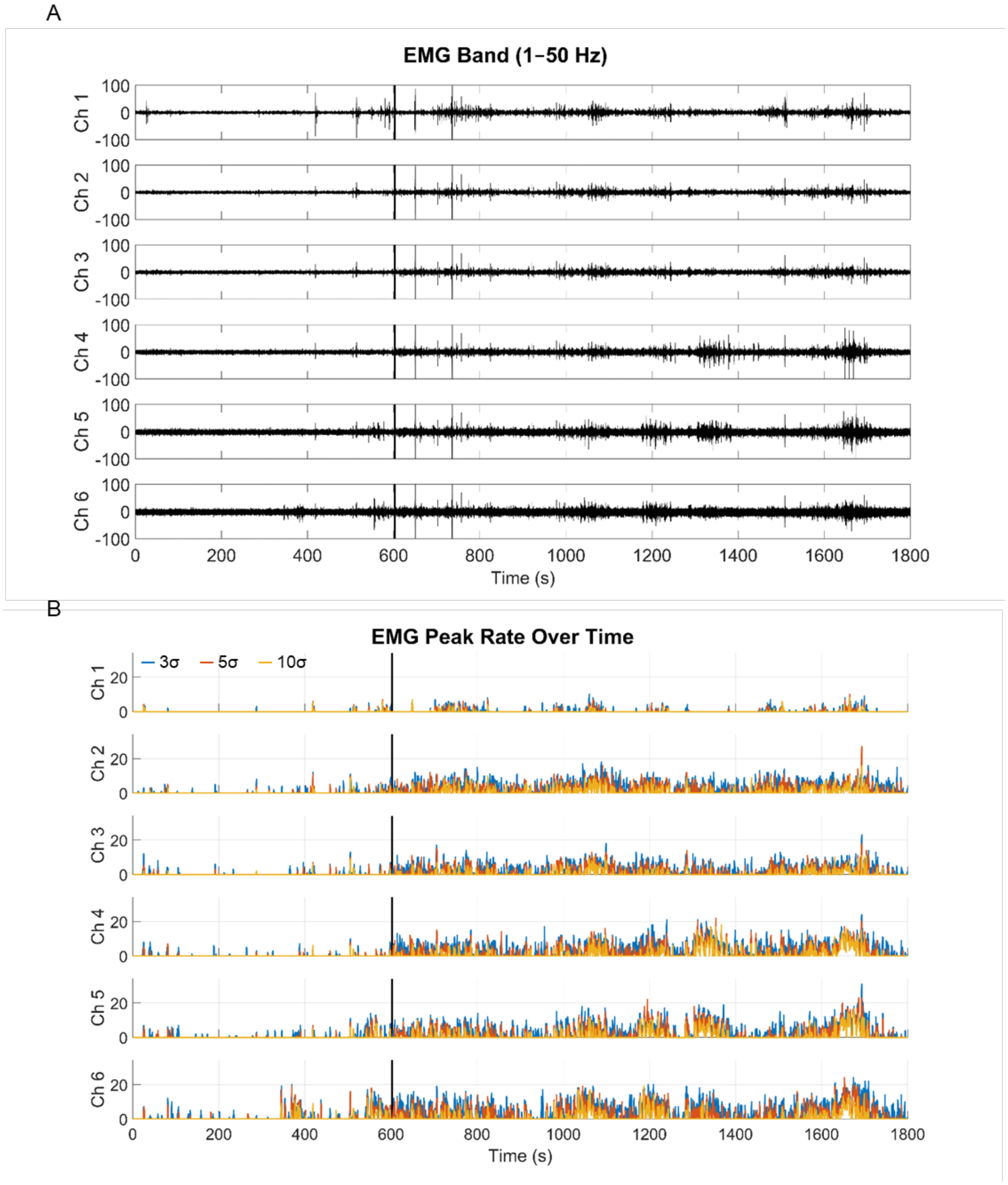


**Fig. S33. Electrophysiological recordings during pharmacological stimulation in Animal 4 (channels 1–6).** (**A**) EMG-band (1–50 Hz) signals before and after drug injection. (**B**) EMG peak rate across time. Black region marks drug delivery.

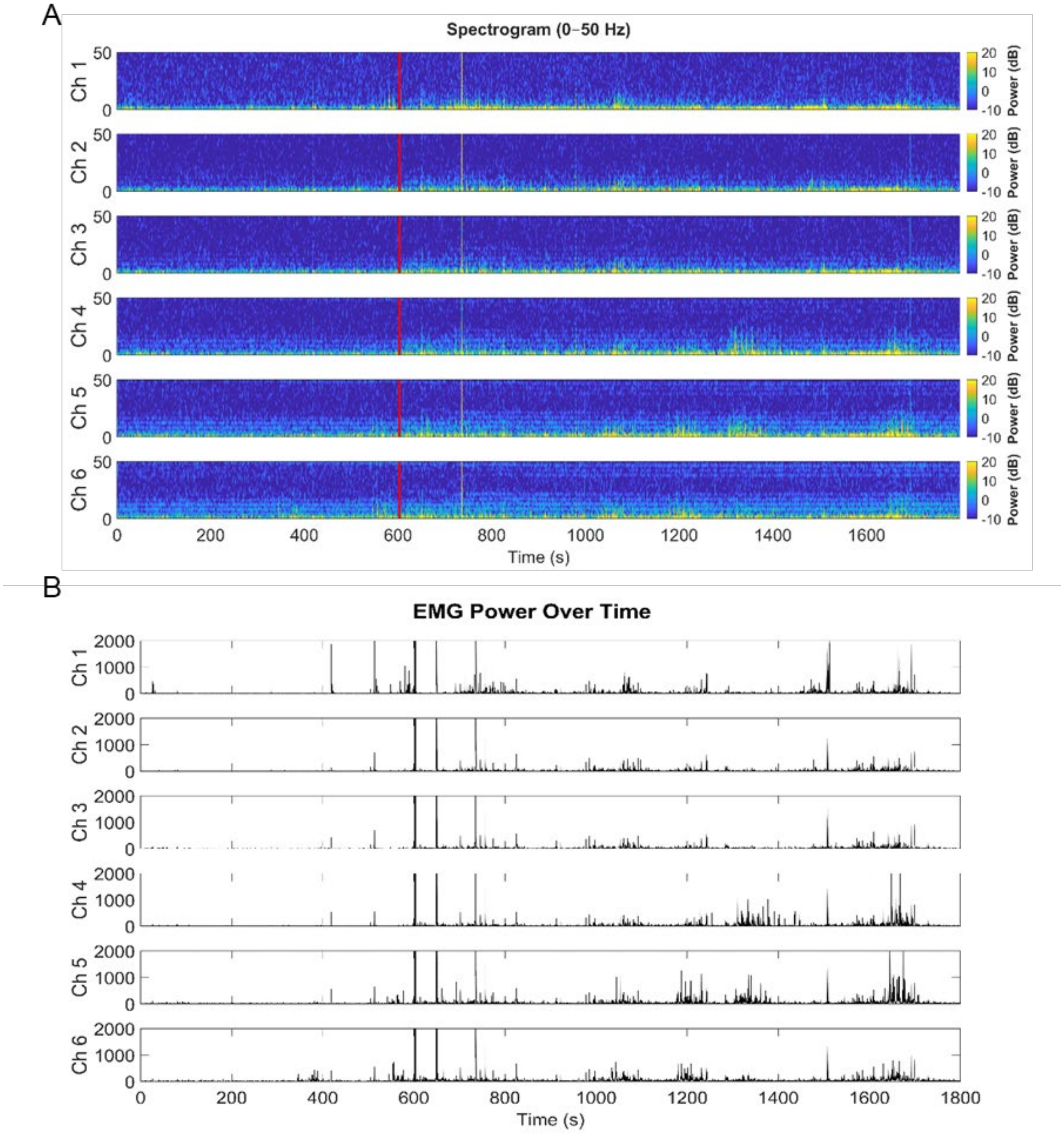


**Fig. S34. Analysis of pharmacological stimulation in Animal 4 (channels 1–6).** (**A**) EMG-band spectrogram (0–50 Hz). (**B**) EMG power over time. Black and red lines mark injection time.

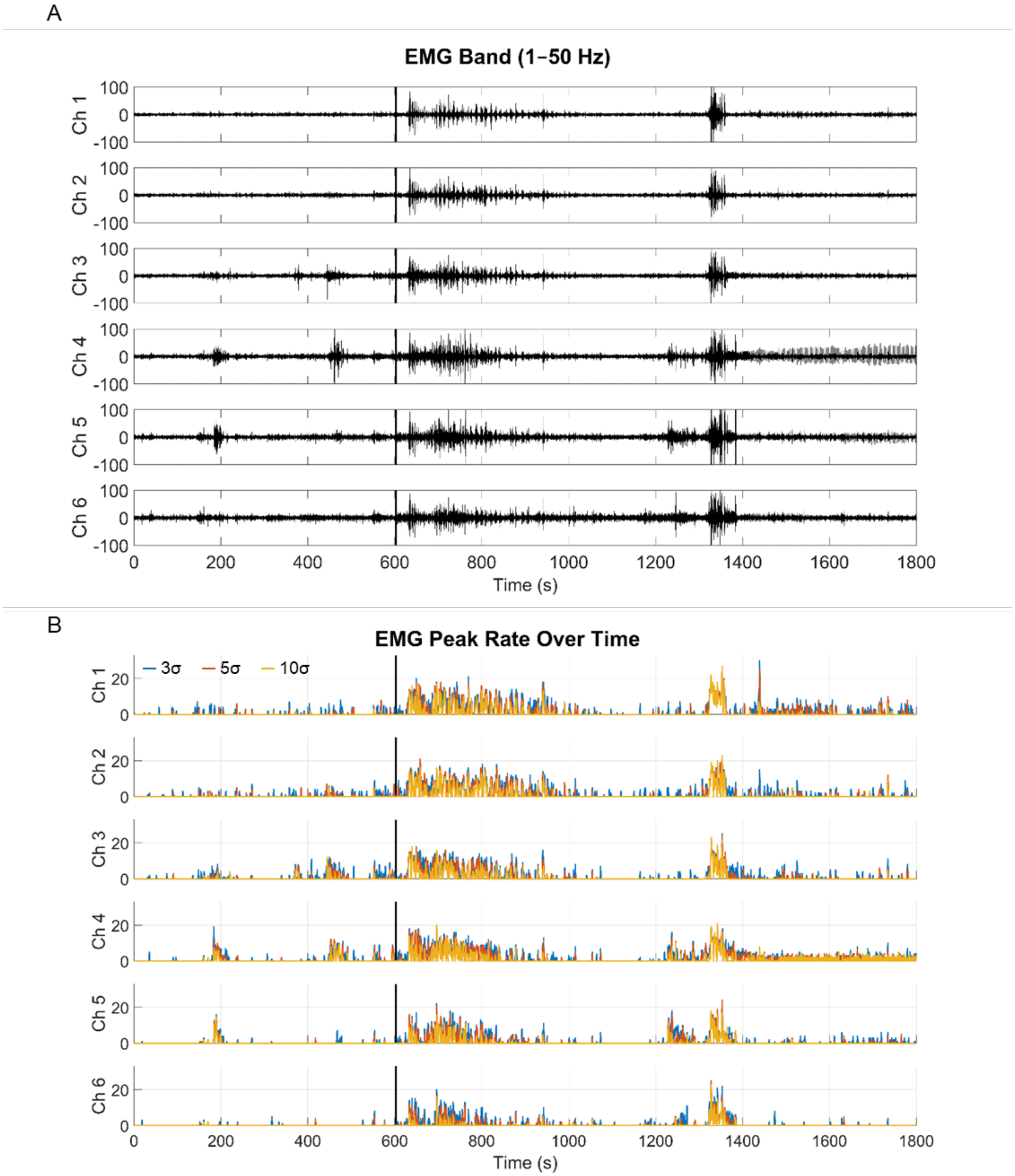


**Fig. S35. Electrophysiological recordings during pharmacological stimulation in Animal 5 (channels 1–6).** (**A**) EMG-band (1–50 Hz) signals before and after drug injection. (**B**) EMG peak rate across time. Black region marks drug delivery.

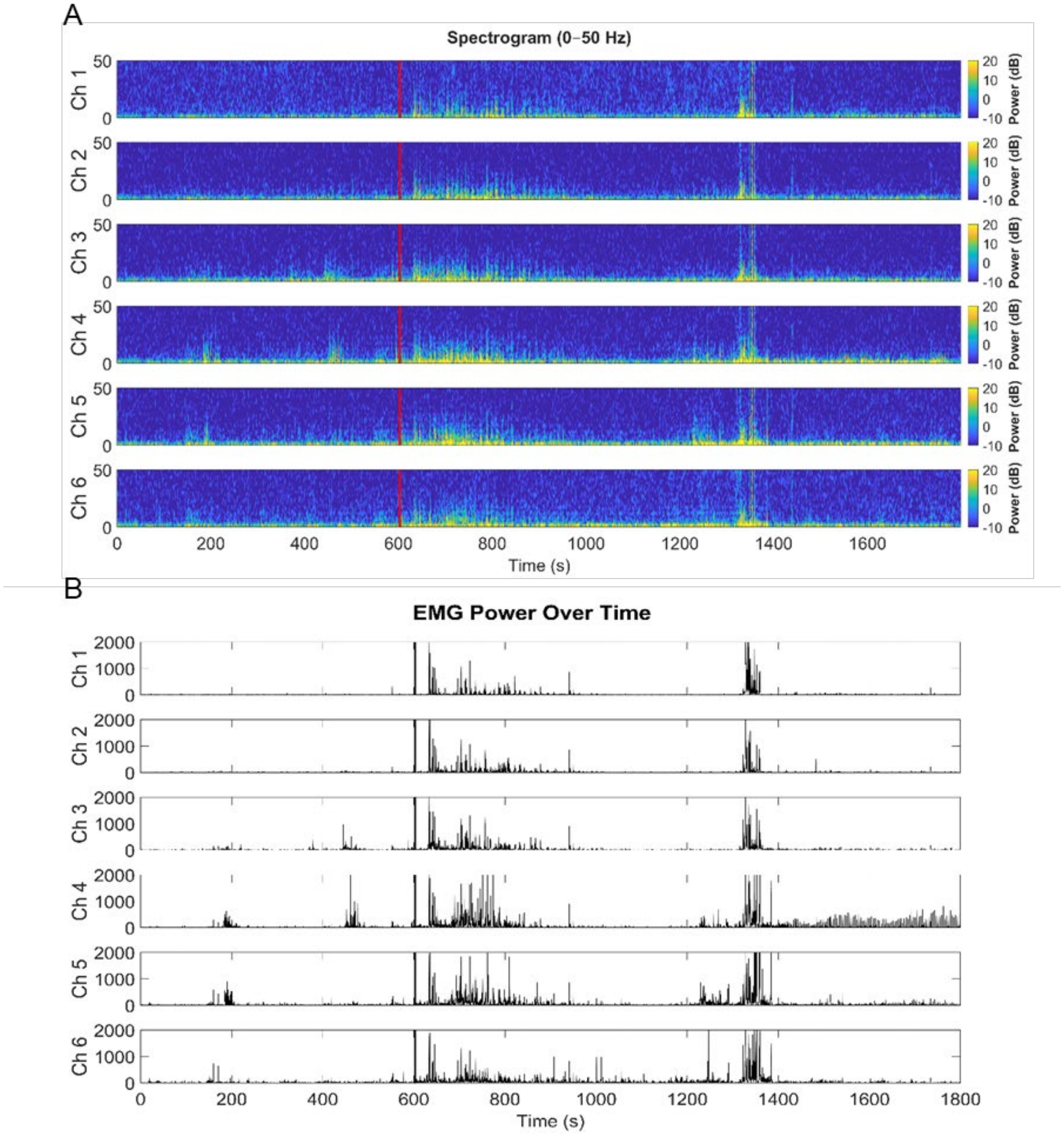


**Fig. S36. Time–frequency analysis during pharmacological stimulation in Animal 5 (channels 1–6).** (**A**) EMG-band spectrogram (0–50 Hz). (**B**) EMG power over time. Black and red lines mark injection time.

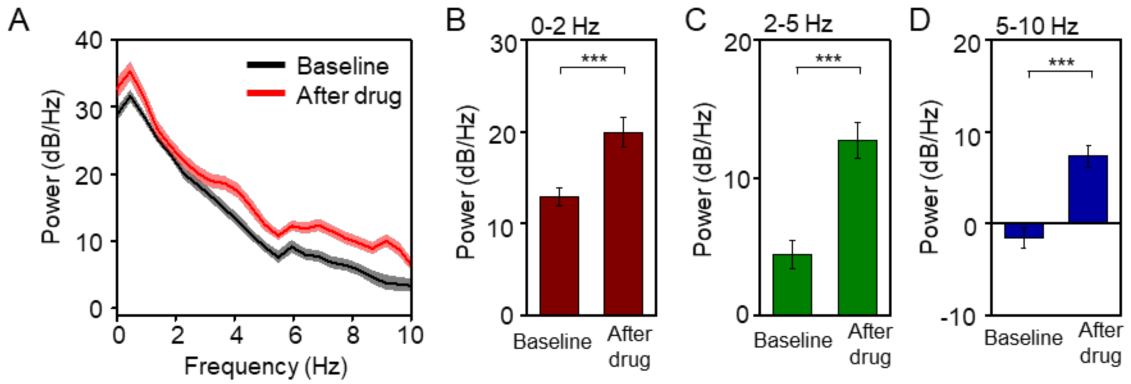


**Fig. S37. Power spectral density analysis of drug across animals 3-5.** (**A**) Power spectral analysis showing changes in frequency response before (baseline) and after (after drug) addition of 20 μL 1 mM bethanechol. Averaged power in the (**B**) 0–2 Hz, (**C**) 2–5 Hz, and (**D**) 5–10 Hz frequency bands before and after addition of 20 μL 1 mM bethanechol. ***$p < 0.001$. Lines and Bars denote mean; shaded regions and error bars denote ± s.e.m. (baseline: n = 18 segments; after drug: n = 18 segments).

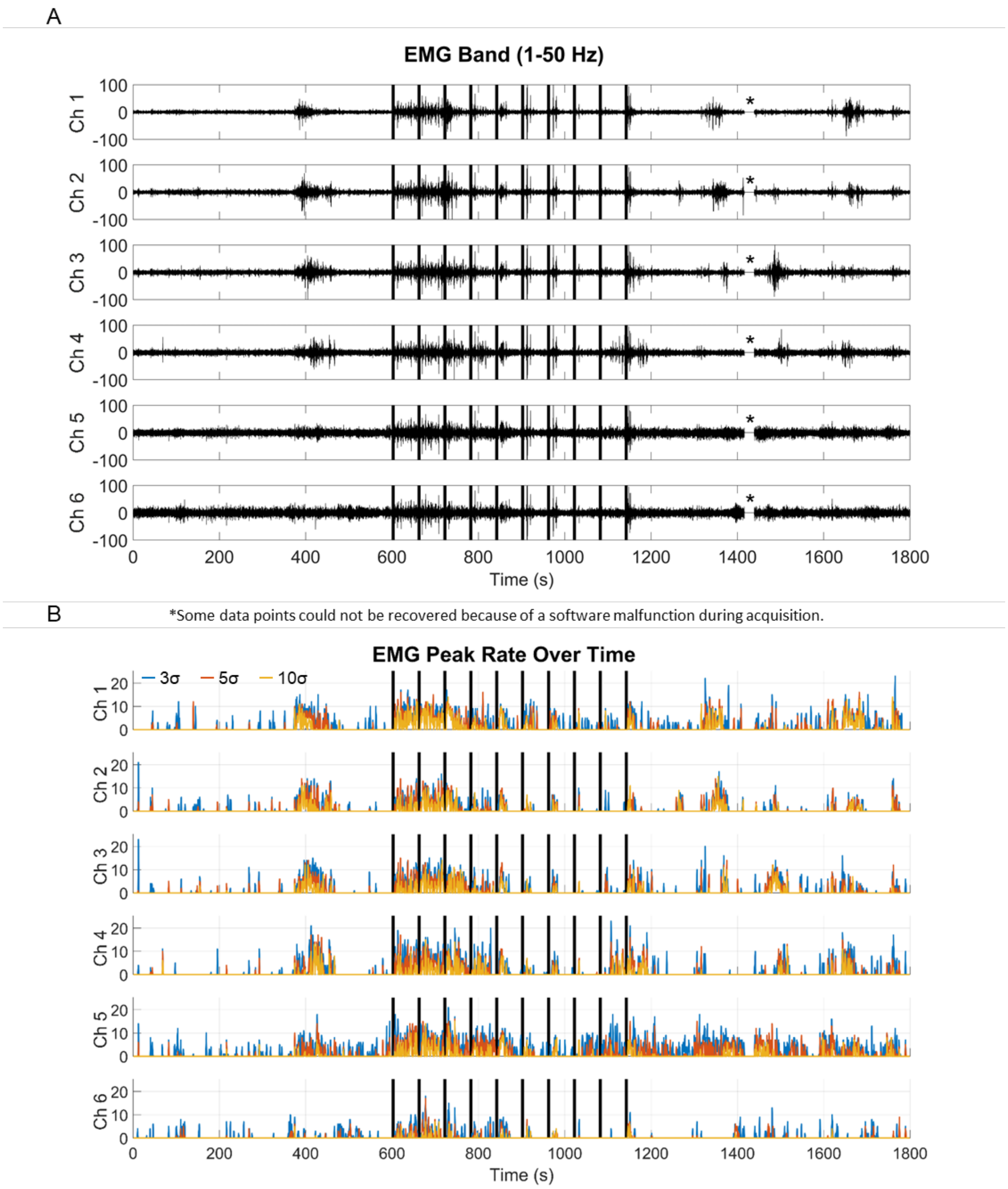


**Fig. S38. Electrophysiological recordings during electrical stimulation in animal 3 (channels 1–6).** (**A**) EMG-band (1–50 Hz) signals in response to pulse bursts (10 repetitions of 3 s of 60 biphasic pulses at 750 µA amplitude, 400 µs pulse width, 80 µs interval, 20 Hz). (**B**) EMG peak rate over time. Black bars indicate stimulation epochs.

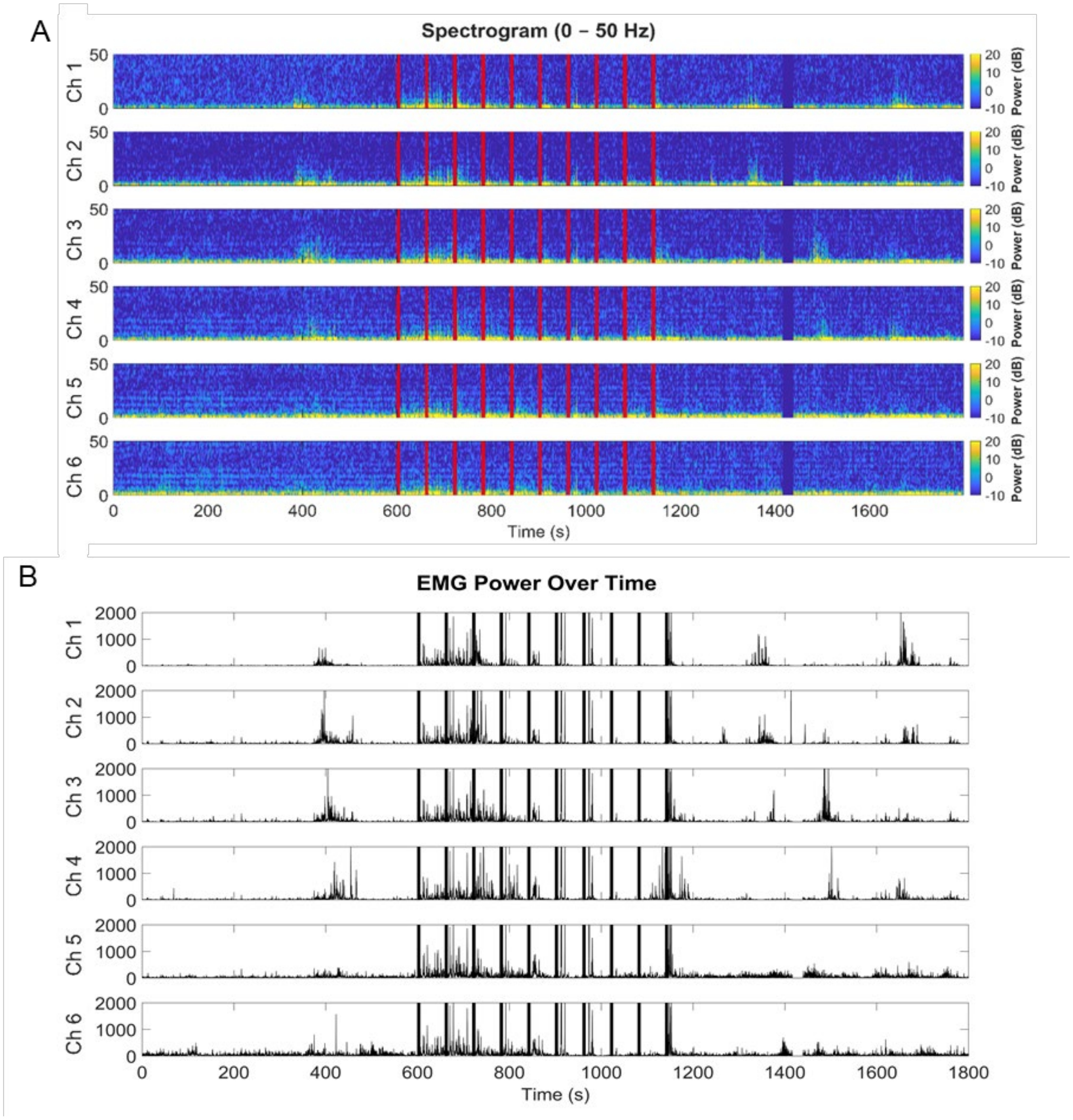


**Fig. S39. Analysis of electrical stimulation in animal 3 (channels 1–6).** (**A**) EMG-band spectrogram (0–50 Hz). (**B**) EMG power over time. Red and black bars indicate stimulation epochs.

A

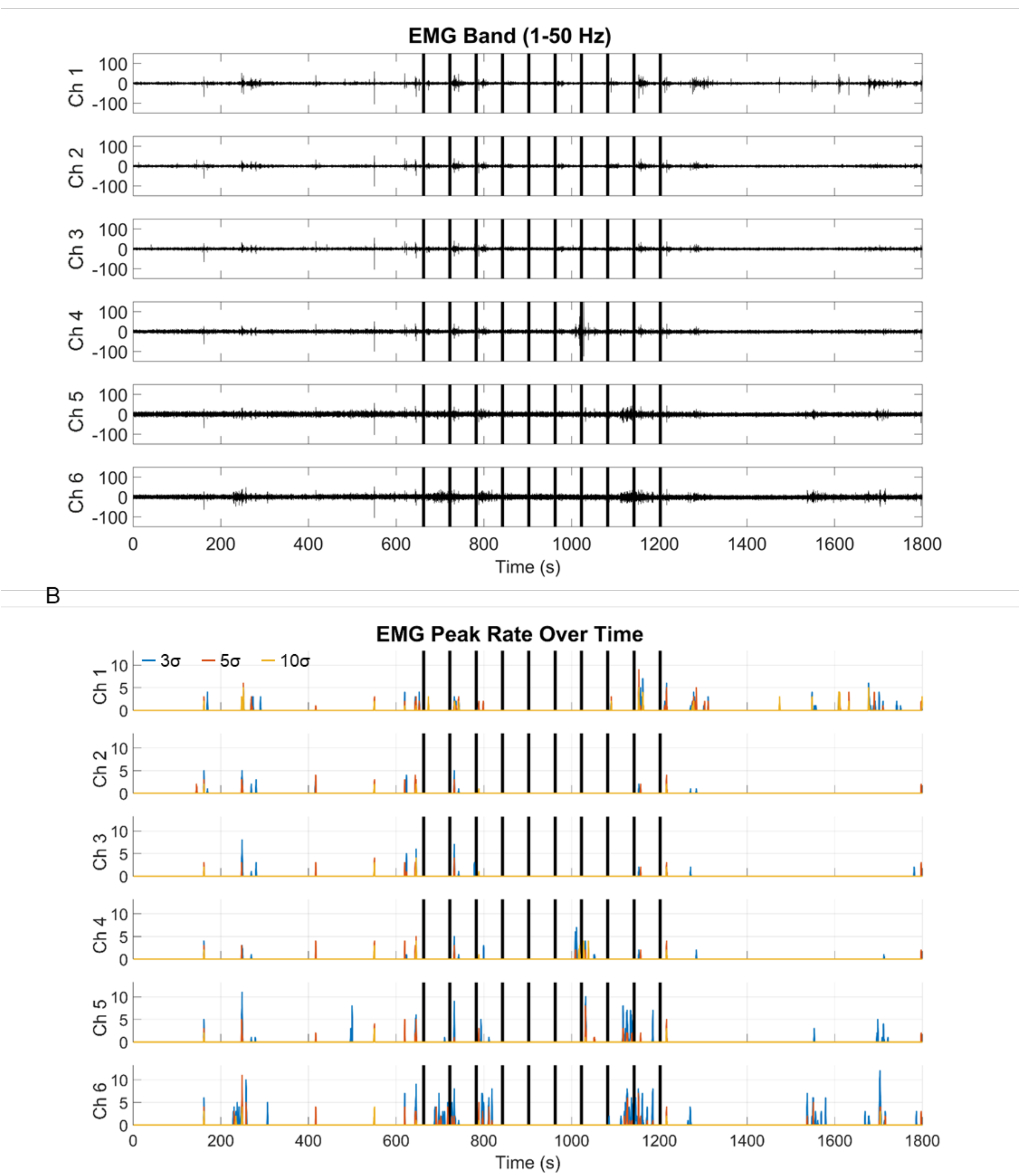


**Fig. S40. Electrophysiological recordings during electrical stimulation in animal 4 (channels 1–6).** (**A**) EMG-band (1–50 Hz) signals in response to pulse bursts (10 repetitions of 60 bipolar pulses). (**B**) EMG peak rate over time. Black bars indicate stimulation epochs.

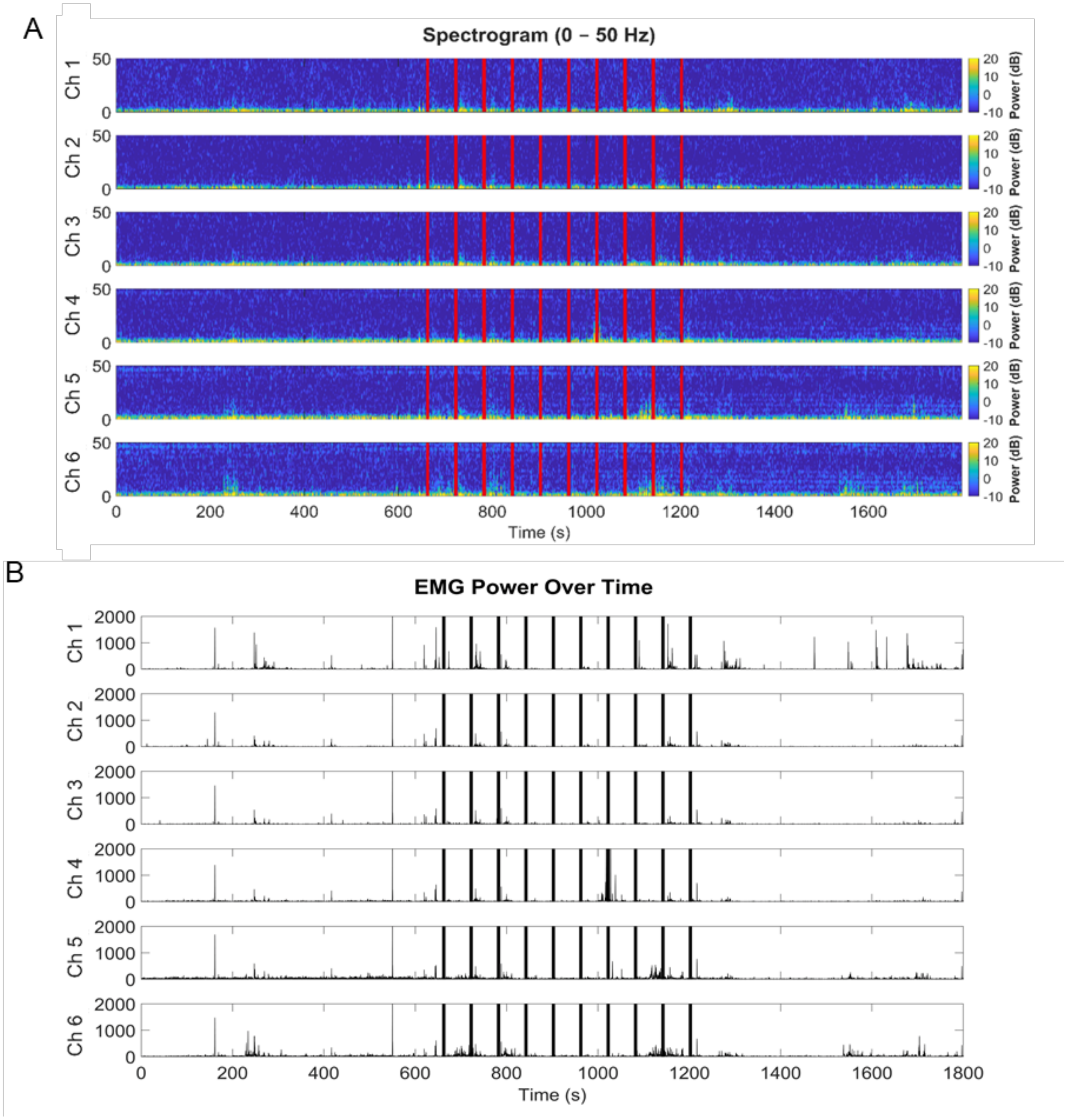


**Fig. S41. Time–frequency analysis during electrical stimulation in animal 4 (channels 1–6).** (**A**) EMG-band spectrogram (0–50 Hz). (**B**) EMG power over time. Red and black lines mark stimulation epochs.

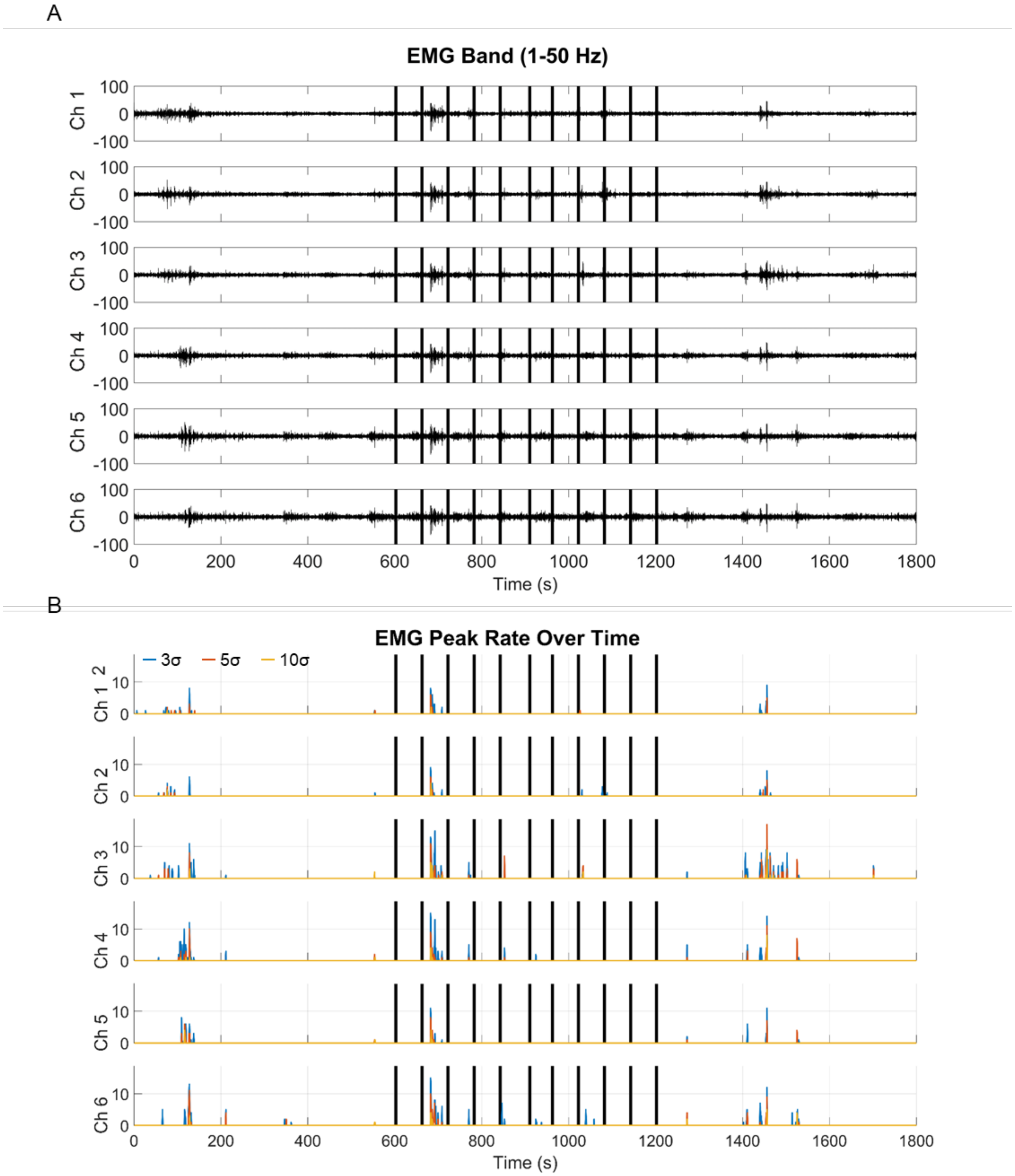


**Fig. S42. Electrophysiological recordings during electrical stimulation in animal 5 (channels 1–6).** (**A**) EMG-band (1–50 Hz) signals in response to pulse bursts (10 repetitions of 60 bipolar pulses). (**B**) EMG peak rate over time. Black bars indicate stimulation epochs.

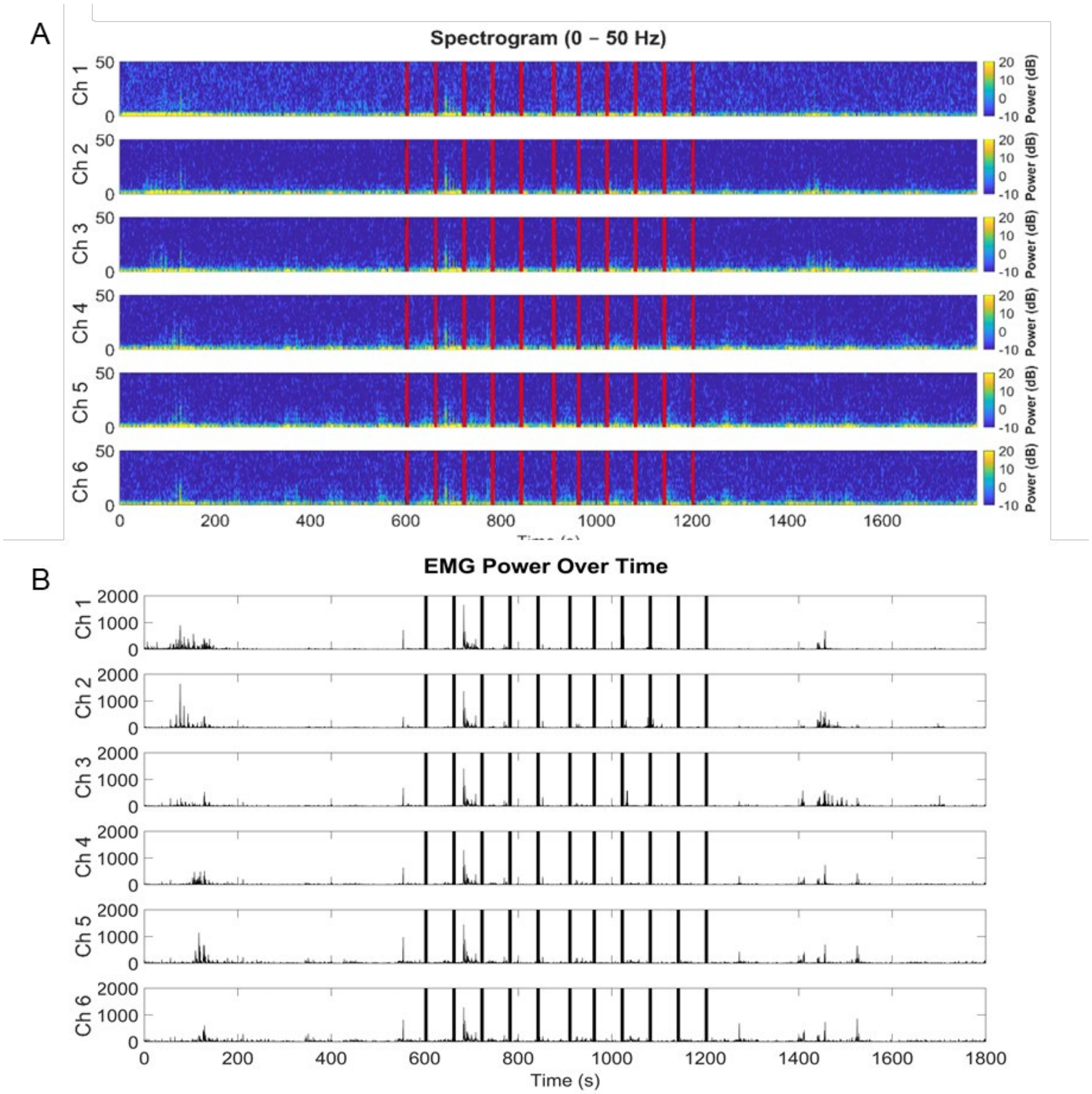


**Fig. S43. Time–frequency analysis during electrical stimulation in animal 5 (channels 1–6).** (**A**) EMG-band spectrogram (0–50 Hz). (**B**) EMG power over time. Red and black lines mark stimulation epochs.

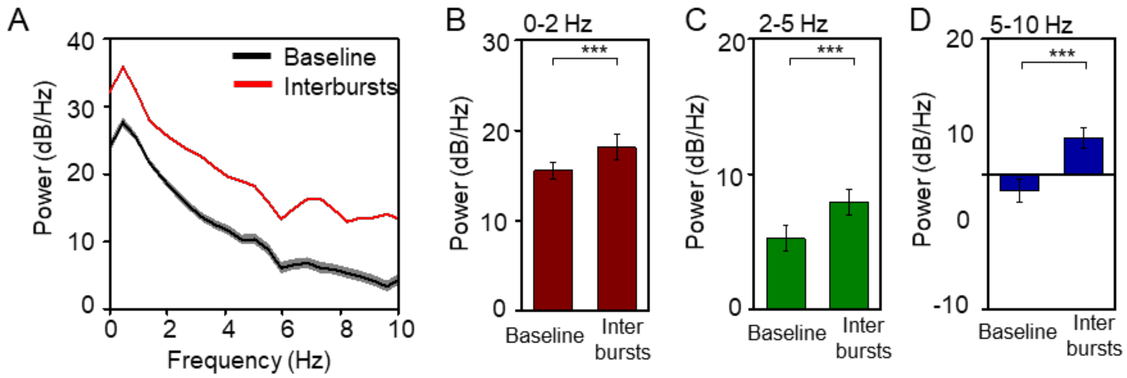


**Fig. S44. Power spectral density (PSD) analysis of electrical stimulation across Animals 3–5.** (**A**) Power spectral density (PSD) comparing baseline and interburst periods. Averaged power within the (B) 0–2 Hz, (C) 2–5 Hz, and (D) 5–10 Hz bands for baseline vs. interbursts. ***$p < 0.001$. Lines and Bars denote mean; shaded regions and error bars denote ± s.e.m. (n = 18).

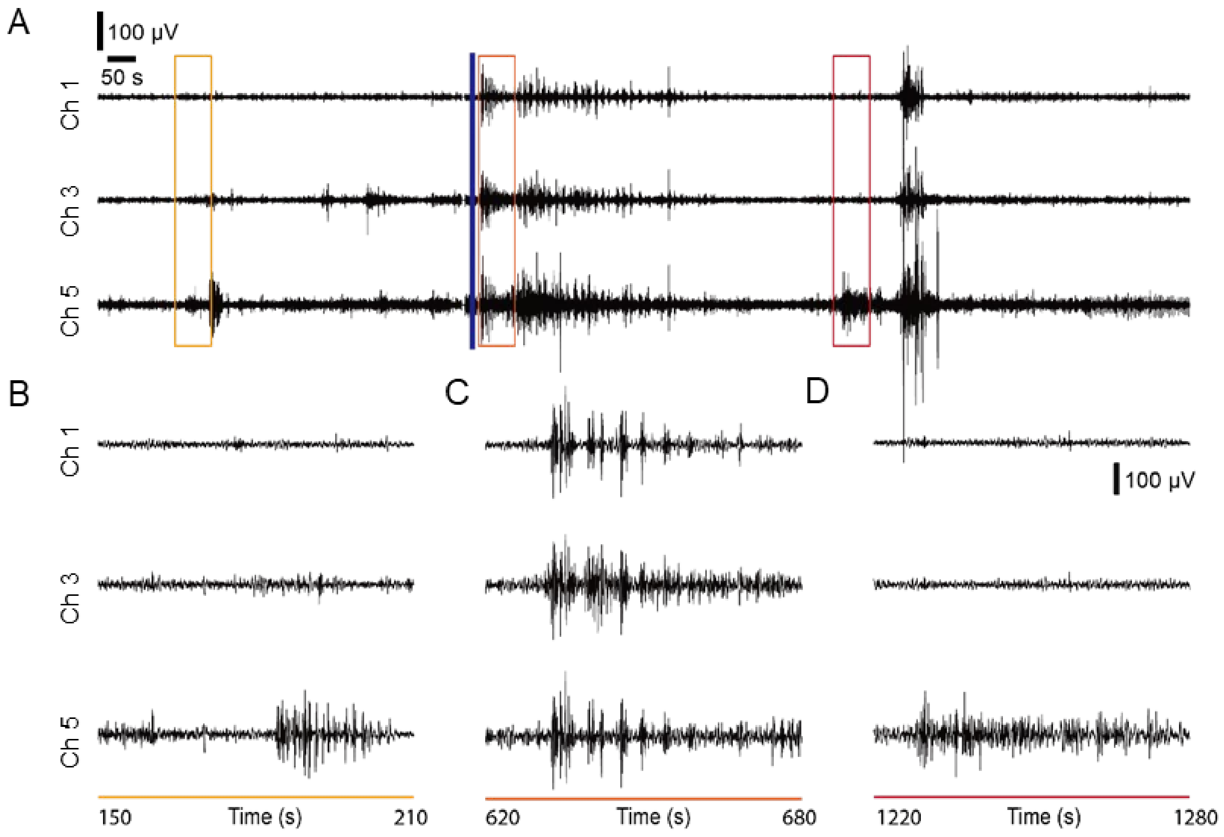


**Fig. S45. Local EMG recording in response to bethanechol in animal 5.** (**A**) EMG band (1–50 Hz) signals in response to bethanechol (1 mM). Blue line indicates the time of drug application. (**B** to **D**) Expanded views of the EMG traces shown in (A).

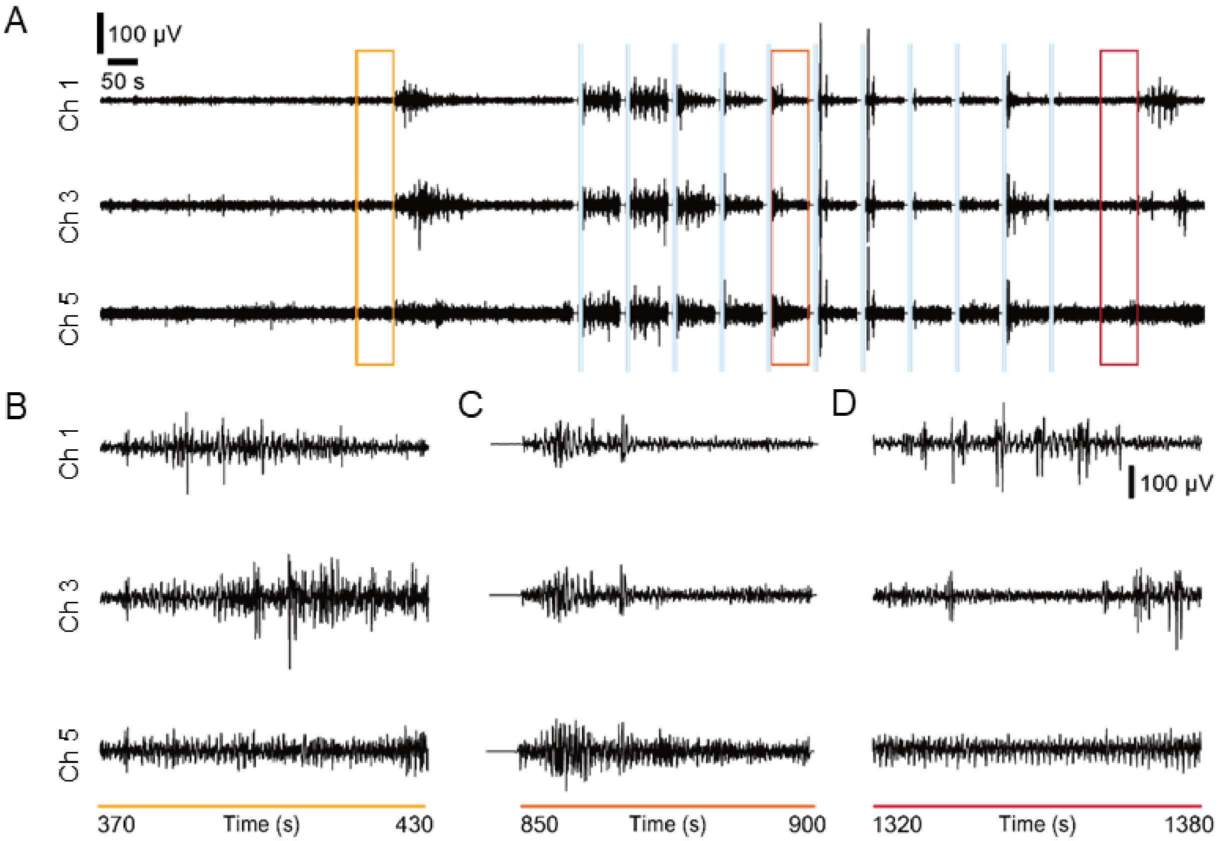


**Fig. S46. Local EMG recording in response to electrical stimulation in animal 3.** (**A**) EMG band (1–50 Hz) signals in response to pulse-burst stimulation. Blue lines indicate the timing of stimulation. (**B** to **D**) Expanded views of the EMG traces shown in (A).

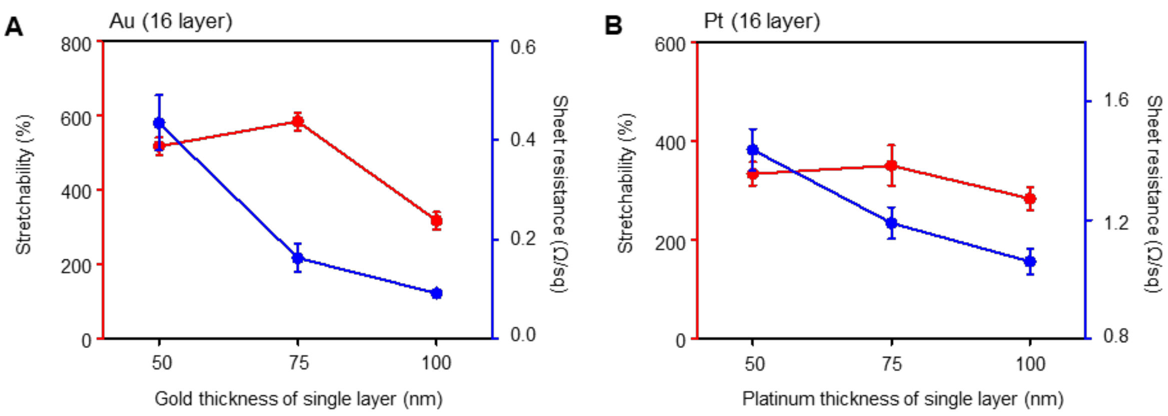


**Fig. S47. Stretchability and sheet resistance of 16-layer Au and Pt stacks as a function of individual metal nanomembrane (NM) thickness.** (**A** and **B**) Stretchability and sheet resistance measurements of 16-layer (A) Au and (B) Pt stacks fabricated with varying single-layer metal NM thicknesses. The PIB solutions used for NM preparation were 0.03 wt% for Au and 0.025 wt% for Pt, respectively. Markers and error bars represent mean ± s.d. (n = 3).

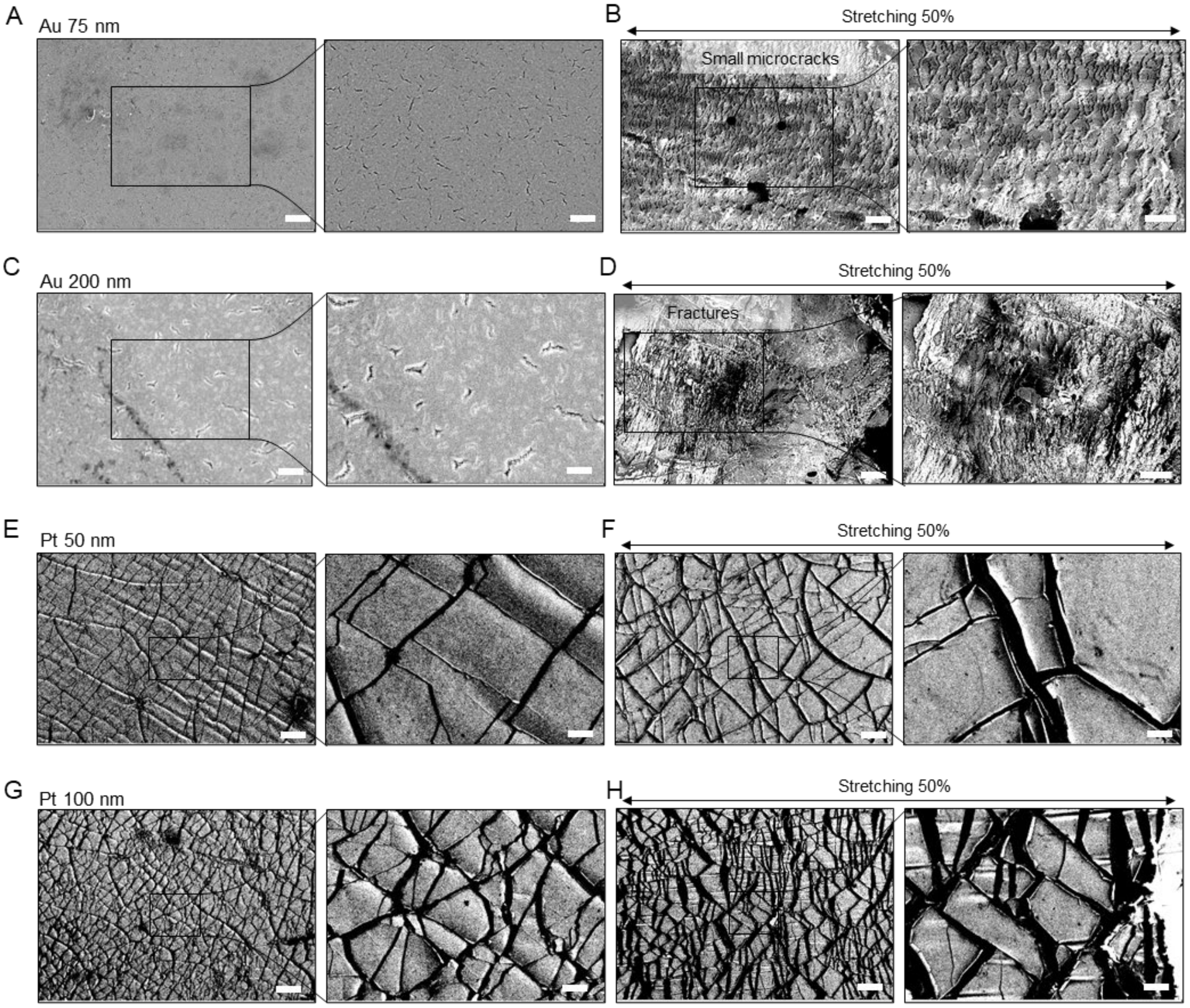


**Fig. S48. Structural limitations of conventional deposition-based metal films on elastomers.** (**A** to **D**) SEM images of Au with thicknesses of (A and B) 75 nm and (C and D) 200 nm, shown (A and C) before and (B and D) after 50% stretching. Scale bars: (A and C) (left) 2 µm, (right) 1 µm; (B) (left) 10 µm, (right) 5 µm; (D) (left) 200 µm, (right) 100 µm. (**E** to **H**) SEM images of Pt with thicknesses of (E and F) 50 nm and (G and H) 100 nm (E and G) before and (F and H) after 50% stretching. Scale bars: (left) 100 µm, (right) 20 µm. (**I**) Stretchability of Au and Pt films on SEBS elastomer depending on metal thickness.

| Figure | Panel | Condition | Comparison | Metric | Method | n | p | |
|---|---|---|---|---|---|---|---|---|
| Fig. 4 | G | Drug | Baseline vs Saline | Peak count | LMM (Satterthwaite DF) | n=12 (2 animals, 6 electrodes) | 0.001313417 | ** |
| | | | Baseline vs 10 μM | | | | 5.38E-06 | *** |
| | | | Baseline vs 100 μM | | | | 2.14E-14 | *** |
| | | | Baseline vs 1000 μM | | | | 1.25E-27 | *** |
| | K | Electrical (pulse width) | Baseline vs 200 μs | | | n=36 (2 animals, 6electrodes, 3 segments) | 0.625354101 | ns |
| | | | Baseline vs 400 μs | | | | 2.42E-18 | *** |
| | | | Baseline vs 800 μs | | | | 2.69E-15 | *** |
| | O | Electrical (amplitude) | Baseline vs 250 μA | | | | 0.736876771 | ns |
| | | | Baseline vs 500 μA | | | | 8.87E-08 | *** |
| | | | Baseline vs 750 μA | | | | 1.90E-22 | *** |
| fig. S22 | A | Drug | Baseline vs saline | Power (0-2 Hz) | | n=12 (2 animals, 6 electrodes) | 1.98E-16 | *** |
| | | | Baseline vs 10 μM | | | | 1.92E-19 | *** |
| | | | Baseline vs 100 μM | | | | 2.52E-28 | *** |
| | | | Baseline vs 1000 μM | | | | 1.40E-35 | *** |
| | B | | Baseline vs saline | Power (2-5 Hz) | | | 1.07E-40 | *** |
| | | | Baseline vs 10 μM | | | | 1.54E-45 | *** |
| | | | Baseline vs 100 μM | | | | 3.93E-68 | *** |
| | | | Baseline vs 1000 μM | | | | 2.97E-67 | *** |
| | C | | Baseline vs saline | Power (5-10 Hz) | | | 5.15E-67 | *** |
| | | | Baseline vs 10 μM | | | | 3.19E-65 | *** |
| | | | Baseline vs 100 μM | | | | 2.42E-97 | *** |
| | | | Baseline vs 1000 μM | | | | 1.59E-76 | *** |
| fig. S23 | C | Saline 100 μL | Baseline vs After saline | Power (0-2 Hz) | | | 0.006507754 | ** |
| | D | | | Power (2-5 Hz) | | | 0.874819404 | ns |
| | E | | | Power (5-10 Hz) | | | 0.073367606 | ns |
| fig. S27 | A | Electrical (pulse width) | Baseline vs 200 μs | Power (0-2 Hz) | | n=36 (2 animals, 6electrodes, 3 segments) | 0.000585971 | *** |
| | | | Baseline vs 400 μs | | | | 2.78E-08 | *** |
| | | | Baseline vs 800 μs | | | | 5.84E-08 | *** |
| | B | | Baseline vs 200 μs | Power (2-5 Hz) | | | 4.40E-05 | *** |
| | | | Baseline vs 400 μs | | | | 2.07E-38 | *** |
| | | | Baseline vs 800 μs | | | | 2.85E-34 | *** |
| | C | | Baseline vs 200 μs | Power (5-10 Hz) | | | 0.978336152 | ns |
| | | | Baseline vs 400 μs | | | | 1.25E-32 | *** |
| | | | Baseline vs 800 μs | | | | 1.83E-27 | *** |
| fig. S30 | A | Electrical (amplitude) | Baseline vs 250 μA | Power (0-2 Hz) | | | 0.001135418 | ** |
| | | | Baseline vs 500 μA | | | | 3.99E-18 | *** |
| | | | Baseline vs 750 μA | | | | 4.40E-19 | *** |
| | B | | Baseline vs 250 μA | Power (2-5 Hz) | | | 0.04415549 | * |
| | | | Baseline vs 500 μA | | | | 1.01E-25 | *** |
| | | | Baseline vs 750 μA | | | | 1.23E-36 | *** |
| | C | | Baseline vs 250 μA | Power (5-10 Hz) | | | 0.032227454 | * |
| | | | Baseline vs 500 μA | | | | 1.03E-14 | *** |
| | | | Baseline vs 750 μA | | | | 2.47E-22 | *** |
| fig. S37 | A | Drug | Baseline vs drug | Power (0-2 Hz) | | n=12 (2 animals, 6 electrodes) | 2.39E-28 | *** |
| | B | | | Power (2-5 Hz) | | | 1.44E-42 | *** |
| | C | | | Power (5-10 Hz) | | | 9.78E-66 | *** |
| fig. S44 | A | Electrical | Baseline vs interbursts | Power (0-2 Hz) | | n=18 (3 animals, 6 electrodes) | 0.000852064 | *** |
| | B | | | Power (2-5 Hz) | | | 4.58E-11 | *** |
| | C | | | Power (5-10 Hz) | | | 8.67E-26 | *** |

**Table S1. Statistical summary for all main and supplementary figures.**

**Movie S1. Hydrogel-assisted adhesion of the stretchable device to the mouse colon under tensile stress.**

**Movie S2. Nanomembrane electrode arrays record spontaneous colonic EMG signals corresponding to peristaltic activity.**

**Movie S3. Nanomembrane electrode arrays stimulate colonic contractions and record the corresponding EMG activity.**
